# Promoter evolution in HIV-1C establishes latent reservoirs highly resistant to reversal

**DOI:** 10.1101/2025.08.02.668273

**Authors:** Disha Bhange, Arun Panchapakesan, Swarnima Mishra, Swati Singh, Dolly Parihar, Chhavi Saini, Manisenthil Shanmugam, Hrimkar Buch, Jayendra Singh, Riya Manna, Manisha Sharma, Shobith Suresh, Narendra Nala, Siddappa N Byrareddy, Shesh Prakash Maurya, Mary Dias, Bimal Kumar Das, Kailapuri G Murugavel, Aylur K Srikrishnan, Tapas K Kundu, Udaykumar Ranga

## Abstract

Latent viral reservoirs remain a major barrier to curing HIV-1, with the long-terminal repeat (LTR) and Tat playing crucial roles in regulating viral transcription. Subtype-specific transcription factor binding site (TFBS) variations within the LTR significantly influence latency and reservoir stability. In earlier work, we identified HIV-1C LTR variants with duplicated TFBS motifs, including NF-κB, AP1, RBEIII, and TCF-1α/LEF-1. Using five cell models, including Jurkat and primary CD4⁺ T cells, we compared canonical R-LTR and variant R2-LTR strains. Across sub-genomic reporters, single-round infections, and full-length viral vectors, we found that the balance between RBEIII and NF-κB motifs governs stability of latency. The ‘two-viruses-one-cell’ system that normalized confounding environmental factors further revealed that latency is primarily controlled by intrinsic transcriptional circuits rather than external stimuli. In longitudinal studies of HIV-1⁺ individuals from acute and chronic infection phases, we observed dominant R strains during early infection and the spontaneous emergence of R2 strains in nearly half of chronic-phase subjects, a process accelerated by ART. Upon CD4⁺ T cell activation, R strains preferentially rebounded, while R2 strains showed strong resistance to reversal, even in subjects harbouring a co-infection. Together, these findings establish the clinical significance of LTR variation in latency regulation and identify the R2 phenotype as a critical determinant of reservoir persistence. These results underscore the importance of addressing reservoir heterogeneity in cure strategies, particularly in HIV-1C-prevalent regions.

## Introduction

Following integration, HIV-1 may either produce infectious particles or remain transcriptionally silent, forming a stable latent reservoir resistant to cure ^1^ ^2^. Several replication-competent latent proviral strains remain unresponsive to cellular activation, contributing to a highly stable and dynamic reservoir^3^. Reservoir stability is influenced by both intrinsic mechanisms and extrinsic factors, including epigenetic modifications at the integration site ^4^ ^5^ ^6^. Although antiretroviral therapy (ART) effectively suppresses viral replication, it fails to clear or significantly reduce the latent reservoir, even after long-term use ^7^. Current experimental strategies have achieved limited reactivation of latent strains, underscoring the need for deeper insights into reservoir maintenance to inform curative efforts.

HIV-1 regulates viral gene expression through a single promoter, the long terminal repeat (LTR), enabling centralized ON/OFF control of transcription. The LTR orchestrates two opposing functions: transcriptional activation and suppression, guided by the transcription factor-binding site (TFBS) profile, which recruits specific host factor complexes. The LTR contains densely arranged TFBS for various transcription factor families, including NF-κB, NF-AT, USF, TCFα/LEF, AP1, and SP1 ^8^. The composition of transcription factors at the LTR is dynamic and complex. The TFBS represent diverse families capable of forming homo- or heterodimer complexes with transcription-inducing or -suppressing properties. Although the core structure of the LTR is conserved across HIV-1 subtypes, subtype-specific TFBS variations may influence potential differences in transcriptional decision-making.

Molecular features unique to HIV-1 subtypes and their impact on reservoir stability remain underexplored. For example, HIV-1C typically contains three NF-κB motifs in its LTR enhancer, compared to two in most other subtypes (Figure S1). We also identified HIV-1C strains in India containing four NF-κB motifs, correlating with proportionately enhanced viral transcription ^9^. In a subsequent study, we reported the emergence of additional LTR-variant strains of HIV-1C containing increased copies of existing TFBS, particularly NF-κB (N) and RBEIII (R) ^10^. Variations in the number, sequence, and position of these TFBS create diverse LTR configurations, highlighting the evolving nature of HIV-1C LTRs and their potential impact on transcription and reservoir stability.

Among LTR variants, those with duplicated RBEIII (R vs. R2, with one vs. two copies) and/or NF-κB motifs (N3 vs. N4, with three vs. four copies) are especially noteworthy. The duplication of the RBEIII motif, termed the most frequent nucleotide length polymorphism (MFNLP), is found across all HIV-1 subtypes, including HIV-1C ^11^ ^12^ ^13^. In contrast, NF-κB motif duplication is exclusive to HIV-1C ^9^, making the co-duplication of NF-κB and RBEIII motifs a unique feature of this subtype. The TF-binding landscape of the LTR reflects the coordinated action of host factors supporting or suppressing viral transcription based on host cell activation. However, Individual TFBS may play a dominant role in either promoting or suppressing transcription. For instance, under activating conditions, NF-κB motifs enhance transcription in a copy number-dependent manner ^9^. In contrast, as shown in this work, in the absence of activation, RBEIII motif duplication, along with its associated factors, such as Ap1, appears to stabilize latency. The co-duplication of NF-κB and RBEIII motifs in HIV-1C is, therefore, of particular interest due to their potentially opposing effects on transcriptional regulation.

To this end, using five cell models, including long-term cultures of primary CD4 cells, we evaluated latency establishment and reversal in LTR-variant strains differing in RBEIII and NF-κB motif duplication. We found that RBEIII motif duplication, termed the ‘R2 genotype’, significantly raises the activation threshold required for latency reversal, here referred to as the ‘R2 phenotype’. Analyses of cross-sectional and longitudinal samples from acute and chronic infection stages, including pre- and post-ART and mixed R/R2 infections, consistently show that R2 strains resist latency reversal in vivo. Although the R2 genotype arises naturally, ART appears to accelerate its selection. Collectively, our findings demonstrate, for the first time, that the R2 phenotype in HIV-1C supports the establishment of more stable latency by requiring stronger activation signals for reactivation.

## Results

### A fine balance exists between positive and negative regulatory elements of the LTR

Preliminary data from our lab suggest that a precise balance between NF-κB (N; three or four copies) and RBEIII (R; one or two copies) motifs in the LTR is critical for regulating HIV-1 latency establishment, maintenance, and reversal. Disruption of this balance, such as the creation of an additional copy of one motif without the other, may alter latency dynamics. The emergence of multiple HIV-1C promoter variants ^10^ offered a valuable opportunity to test the hypothesis that differences in the TFBS profile may modulate latency kinetics. In particular, two strains, R2N3 and R2N4, differ in their TFBS composition relative to the canonical RN3 virus. R2N3 includes a duplicated RBEIII motif, while R2N4 carries additional copies of both RBEIII and NF-κB motifs (Figure S2). These variants provide a model to dissect how TFBS architecture influences latency kinetics. Of note, although for simplicity, we refer the sequence motif duplication by the identity of the majorly known TFBS, in reality, the duplicated sequences invariably contain binding sites for more than one TF family. For example, the RBEIII motif duplication of must be considered as the duplication of the core motif of eight base pairs flanked by additional sequence motifs co-duplicated for the binding of AP1, LEF-α, and/or NF-κB.

We constructed a panel of four LTR-variant viral strains differing in the copy number of the RBEIII (R Vs. R2) and NF-κB (N3 Vs. N4) motifs (Figure S2). The RN3-LTR corresponds to the canonical HIV-1C promoter, while the RN4-LTR variant emerged naturally over two decades ago ^14^. Unlike these canonical LTRs with a single RBEIII element in the modulator region, R2N3- and R2N4-LTRs carry two RBEIII copies in this region.

Latency establishment and reactivation were assessed using pseudotyped sub-genomic viral strains co-expressing Tat and d2EGFP, as illustrated (Figure 1). All four strains showed comparable downregulation of GFP over time, with ∼90% of infected cells silencing GFP by day 8 (Figure 1B, left panel) and minimal cell death (Figure S3). Reporter gene expression correlated strongly with Tat transcript levels at all time points (Figure S4). While overall latency profiles were similar, dual-RBEIII strains (R2N3 and R2N4) showed a trend toward faster latency establishment compared to canonical strains (RN3 and RN4), with statistically significant differences (Figure 1B, left panel). For instance, at day 4, 51.3 ± 8.6% of RN3-infected cells remained GFP⁺, versus only 20.5 ± 0.7% for R2N3.

**Figure 1:**
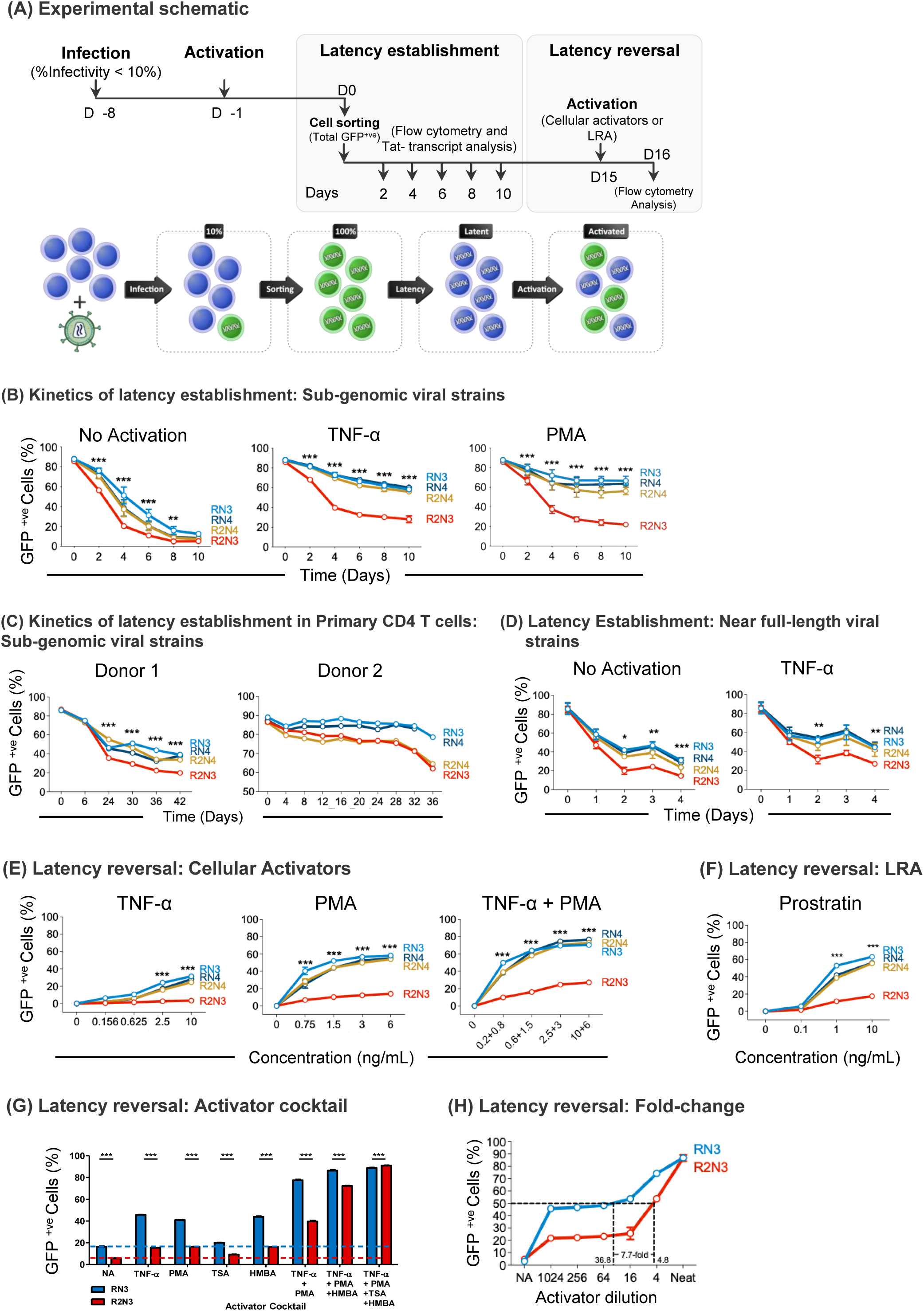
The latency profiles of the LTR-variant viral strains. (A) Experimental schematic of the latency evaluation in Jurkat T cells. A schematic of the workflow and the timeline of the experimentation are depicted (Top panel). Jurkat cells were infected with LTR-variant viral strains of the panel at an RIU of ∼0.1 or 10% infectivity, and the infected cells (GFP+ve) were sorted using FACS. The sorted cells approximately take 15 days to relax the gene expression. (B) The profiles of latency establishment of the sub-genomic promoter-variant viral strains in Jurkat T-cells. Latency establishment was monitored in the absence or presence of sub-optimal concentrations of TNF-α (10 ng/ml) or PMA (5 ng/ml), as illustrated. The data are representative of four independent experiments. (C) The kinetics of latency establishment of the sub-genomic viral strains in primary CD4 cells. Primary CD4 cells stably transduced with Bcl-2 expression were infected with the panel of sub-genomic promoter-variant strains described in supplementary figure 6. The percentage of GFP+ve cells was monitored. (D) The profiles of latency establishment of the near full-length promoter-variant viral strains in Jurkat T cells. Latency establishment was monitored in the absence or presence of TNF-α. (E) The latency reversal kinetics of the panel of sub-genomic viral strains. The latency reversal assays were performed in the presence of increasing concentrations of TNF-α and/or PMA. The data are representative of four independent experiments. (F) The latency reversal profile in Jurkat cells in the presence of increasing concentrations of Prostratin. (G) Differential Latency reversal properties of RN3- and R2N3-LTRs in response to activator cocktails. Jurkat cells were activated with single activators or cocktails of two, three, or four (TNF-α, PMA, TSA, and HMBA) activators as shown. The data are representative of two independent experiments. (H) Activation threshold difference between RN3 and R2N3 LTRs. Jurkat cells harbouring latent proviruses were exposed to a cocktail (Neat) of PMA (5 ng/ml) and HMBA (5 mM) or a serial four-fold dilution of the activation mixture. The data are representative of three independent experiments. Under all experimental conditions, the cell viability was not compromised. For all graphs, the statistical analysis represents a comparison between RN3 and R2N3 viral strains, while comparisons with the remaining viral strains are not marked on graphs. p=0.033(*), p=0.002(**), p<0.001 (***), Two-way ANOVA with Dunnett multiple comparisons test.

To accentuate differences in latency kinetics, viral strains were cultured under suboptimal concentrations of activators—TNF-α or PMA. Under these conditions, all four LTRs exhibited slower latency establishment (Figure 1B, middle and right panels); however, the R2N3-LTR showed a significantly faster silencing of gene expression compared to the other LTRs, including R2N4. Similar latency patterns were observed in primary CD4⁺ T cells from healthy donors (Figure 1C). Notably, this differential latency profile was consistent and reproducible with near-full-length viral strains (Figure 1D). For instance, by day 4, 31.2 ± 1.0% of RN3-infected cells were GFP⁺, whereas only 14.5 ± 1.2% of R2N3-infected cells remained GFP⁺— a statistically significant difference. In summary, the additional RBEIII motif in R2N3-LTR enhances latency establishment, whereas the co-duplication of NF-κB motifs in R2N4-LTR appears to counteract this effect.

To assess latency reversal, infected cells were cultured for 15–20 days, allowing >90% to downregulate GFP and establish latency. Cells were then stimulated with increasing concentrations of TNF-α (0-10 ng/ml), PMA (0-6 ng/ml), or both, and GFP⁺ cells were quantified by flow cytometry. The RN3, RN4, and R2N4 LTRs showed dose-dependent reactivation (Figure 1E), with R2N4 behaving similarly to the canonical LTRs despite its additional RBEIII motif. In contrast, the R2N3-LTR exhibited significant resistance to reactivation, even at the highest doses of TNF-α and PMA, alone or combined. For example, at 10 ng/ml TNF-α plus 6 ng/ml PMA, only 27.1 ± 0.5% of R2N3 cells were GFP⁺ vs. 70.5 ± 1.3% for RN3. No substantial cell death was observed under any condition. The R2N3-LTR remained unresponsive even to supraphysiological TNF-α (10,240 ng/ml) or PMA (1,280 ng/ml), which robustly reactivated RN3 and R2N4 (Figure S5).

Similarly, Prostratin reactivated RN3, RN4, and R2N4 efficiently, but not R2N3 (Figure 1F); at 10 μM, 63.3 ± 1.0% of RN3 cells vs. only 17.5 ± 1.3% of R2N3 cells were GFP⁺. Notably, R2N3 was fully reactivated to RN3 levels using a cocktail of four activators - TNF-α, PMA, TSA, and HMBA - confirming the absence of intrinsic transcriptional defects (Figure 1G). However, any combination of three activators induced only partial activation, and two were insufficient. A dual-activator combination of PMA (5 ng/ml) and HMBA (5 mM) was more effective in reactivating R2N3. Using a dilution series of this cocktail, we found that R2N3 required a 7.7-fold higher concentration than that of RN3 to achieve 50% reactivation (36.8-vs. 4.8-fold dilution; Figure 1H).

In summary, the additional RBEIII motif in the R2N3-LTR raises the activation threshold, rendering it significantly more resistant to latency reversal. Notably, co-duplication of the NF-κB motif, as in R2N4-LTR, offsets this repressive effect, restoring transcriptional responsiveness.

### Increasing NF-κB motif copies can reverse the suppressive effect of the RBEIII motif duplication on latency reversal

These findings support a finely tuned interplay between the transcriptionally activating NF-κB motifs and the repressive RBEIII motifs in HIV-1 LTR regulation. If accurate, reducing NF-κB motifs in the R2N4-LTR should progressively increase resistance to latency reversal. To test this hypothesis, we generated a panel of sub-genomic viral constructs by sequentially deleting NF-κB motifs from the R2N4-LTR (Figure S6), yielding four variants - R2N, R2N2, R2N3, and R2N4 - containing 1 to 4 NF-κB motif copies, respectively. The central κB site adjacent to the Sp1 motifs (C-κB site; ^15^ was preserved in all constructs. Of note, the NF-κB number variant strains RN2 and R2N2 appear naturally, although at a low frequency.

Latency establishment and reactivation profiles in Jurkat cells aligned with the working hypothesis. All LTR variants efficiently established latency without activators, regardless of NF-κB motif number. However, in the presence of sub-optimal TNF-α or PMA, latency kinetics varied based on NF-κB motif copy number (Figure 2A). LTRs with fewer NF-κB motifs (R2N and R2N2) downregulated gene expression more rapidly, irrespective of activator presence. In contrast, R2N3 and R2N4, containing more NF-κB motifs, exhibited slower latency establishment and incomplete silencing. For instance, by day 10 with 5 ng/ml PMA, GFP⁺ cell percentages were 10.0 ± 1.8% (R2N), 8.2 ± 1.0% (R2N2), 21.9 ± 1.5% (R2N3), and 56.2 ± 3.7% (R2N4) (Figure 2A, right panel). These results indicate an inverse relationship between latency establishment rate and NF-κB motif number when RBEIII copy number remains constant at two.

**Figure 2:**
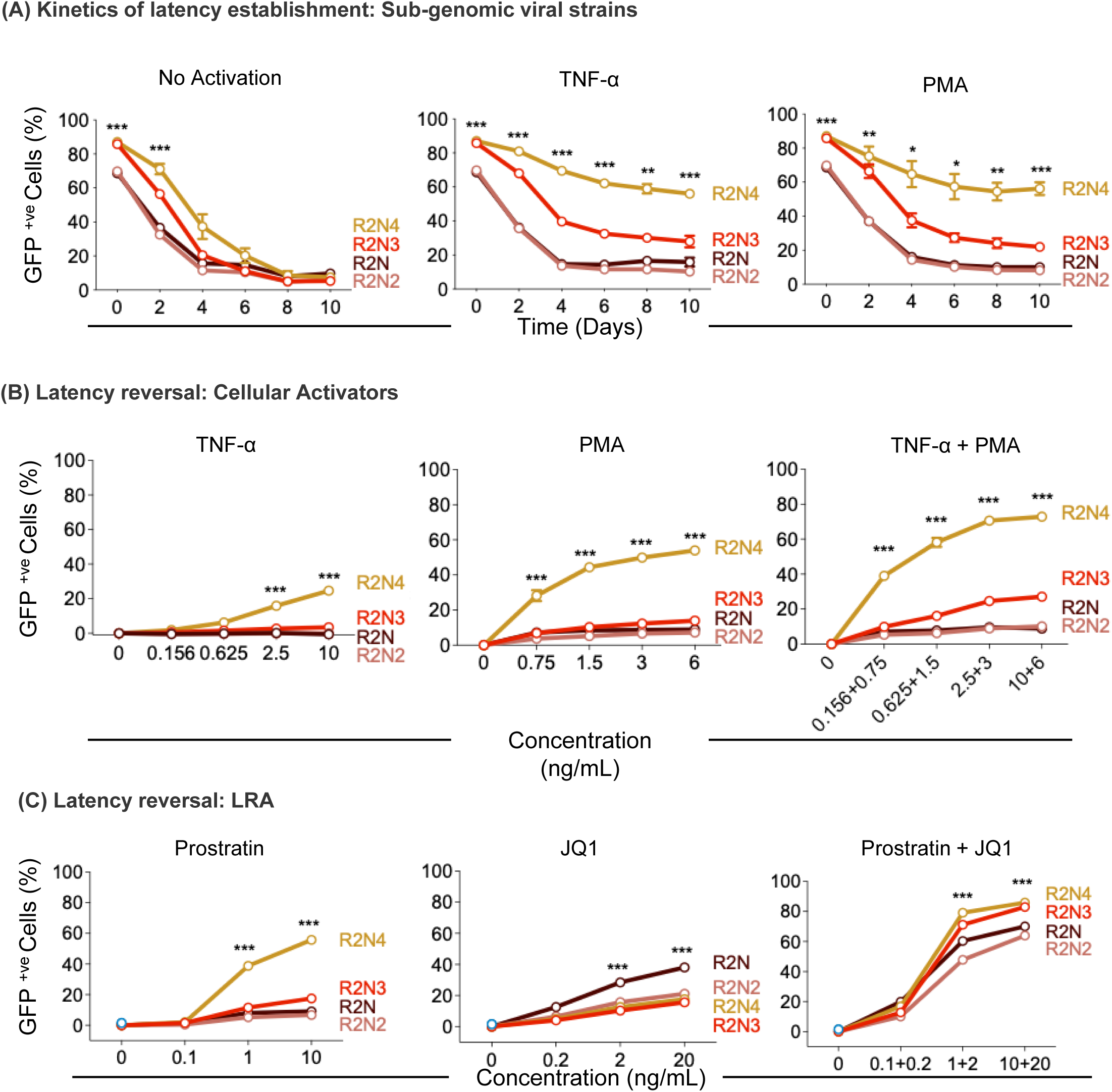
NF-κB motif duplication can reverse the negative impact of the RBEIII motif duplication. The latency profiles of the R2-LTR-variant viral strains.(A) The profiles of latency establishment of the sub-genomic promoter-variant viral strains in Jurkat T-cells. Latency establishment was monitored in the absence or presence of TNF-α, PMA, or both as illustrated. (B) The latency reversal profiles of the R2 viral panel. Induced GFP expression from the latent proviruses was monitored in the presence of increasing concentrations of cellular activators TNF-α and/or PMA, or (C) LRAs Prostratin and/or JQ1, as depicted. For all graphs, the statistical analysis represents a comparison between R2N4 and R2N viral strains while comparisons with remaining viral strains although similar are not marked on graphs. p=0.033(*), p=0.002(**), p<0.001 (***), Two-way ANOVA with Dunnett multiple comparisons test.

In contrast to latency establishment, the latency reversal profiles of the R2 viral strains showed a direct correlation with NF-κB motif copy number (Figure 2B). Under stimulation with TNF-α (10 ng/ml) and PMA (6 ng/ml), LTRs with fewer NF-κB motifs (R2N and R2N2) showed inefficient reactivation, with only 8.9 ± 0.7% and 10.1 ± 0.2% of cells, respectively, becoming GFP⁺. In comparison, R2N4 showed robust reactivation (72.9 ± 0.3% GFP⁺), while R2N3 exhibited intermediate activation (27.1 ± 0.3% GFP⁺). Crucially, latency reversal profiles of the panel were profoundly different in the presence of two conventional LRAs – prostratin and JQ1 (Figure 2C). Especially in the presence of both LRAs, the differences in the latency reversal of the panel nearly disappeared regardless of NF-κB motif difference, despite statistical significance (see Discussion).

Collectively, these findings highlight a delicate balance between the opposing regulatory roles of NF-κB and RBEIII motifs in HIV-1 latency reversal. In LTRs with two RBEIII copies, effective reactivation requires four NF-κB motifs; fewer NF-κB sites result in suboptimal or inefficient latency reversal. Notably, the impact of RBEIII duplication is more pronounced on latency reversal than on its establishment.

### The dual-RBEIII viral strains proliferate at a slower rate in primary CD4 cells and establish latency at a faster rate

We examined latency profiles of ‘near full-length’ LTR-variant viral strains, constructed using the Indie molecular clone (Figure 1C), in primary CD4 cells since these cells serve as the major target of HIV-1 in natural infection. However, efficient infection requires prior activation of these cells, which delays latency establishment and necessitates prolonged culture. Among various published long-term CD4 cell culture methods ^16^ ^17^ ^18^ ^19^ ^20^, we adapted a protocol based on Bcl-2 overexpression ^18^ and further optimized it as detailed below.

In the modified protocol, we used a lentiviral vector to co-express Bcl-2 and mouse CD8a (mCD8a) (Figure S7). Surface expression of mCD8a enabled rapid, positive selection of transduced cells via magnetic beads, reducing the experimental timeline by at least three weeks and improving cell yield. Co-expression of Bcl-2 and mCD8a was confirmed 48 h post-transduction using specific antibodies and flow cytometry (Figure S7C). Subsequently, mCD8a⁺ CD4⁺ T cells were enriched using magnetic beads. These Bcl-2-expressing primary CD4⁺ cells remained viable for 4–6 weeks, allowing latency profile assessments.

Bcl-2-expressing primary CD4⁺ T cells were infected with three sub-genomic HIV-1 strains (RN3, R2N3, and R2N4), and p24 levels in culture supernatants were monitored every three days until cell viability declined (Figure S8). Replication kinetics and culture duration varied across donors. In Donor 1, by day 45, RN3 showed significantly higher replication than R2N3, with R2N4 displaying intermediate levels. Similar trends were observed in Donor 2 on day 15 and in Donors 3 and 4 on their respective replication peaks (days 10 and 18). Overall, both dual-RBEIII strains (R2N3 and R2N4) exhibited markedly slower replication than RN3. Notably, latency establishment in primary CD4⁺ cells was slower and less complete than in Jurkat cells. Factors such as global T cell activation for infection, ectopic Bcl-2 expression potentially affecting LTR activity, and donor-specific host factors likely influenced these outcomes. Nonetheless, viral proliferation patterns in primary CD4⁺ cells generally mirrored latency trends observed in the T-cell line model.

### Within the confines of a single cell, the differential activation profiles of the RN3 and R2N3 LTRs are reproducible

The relative contribution of intrinsic proviral elements and extrinsic cellular factors in regulating HIV-1 transcription and latency remain debated. It is unclear whether latency is primarily driven by host activation status ^21^ ^22^ ^23^or by stochastic viral gene expression ^5^ ^6^ ^24^. Traditional latency models are constrained by the inability to normalize both intrinsic and extrinsic variability. The availability of HIV-1 LTR variants with distinct activation thresholds enabled the development of a model to disentangle these contributions. By introducing two LTRs with divergent activation requirements into a single cell, we normalize environmental variables, including activation signals. Evaluating gene expression at the population level further controls for integration site effects. Thus, we established two-viruses-one-cell models to normalize all experimental variables and assess the influence of viral factors on transcriptional activity.

We engineered sub-genomic HIV-1 vectors using two LTR variants—RN3 (simple-to-activate) and R2N3 (tough-to-activate), each driving expression of d2EGFP or d2mScarlet to independently track LTR activity (Figure 3A). Tat, the most potent latency-reversing agent, was co-expressed from only one LTR via an IRES, which can transactivate both LTRs regardless of its origin. Due to the distinct activation thresholds of the two LTRs, intracellular Tat levels differ depending on which LTR drives Tat expression (Figure S9). Under identical conditions, the RN3-LTR is expected to produce Tat earlier and at higher levels than the R2N3-LTR. Thus, when Tat is expressed from RN3, both LTRs should exhibit stronger and more comparable activation than when Tat is expressed from R2N3. To test this, we developed two ‘two-viruses-one-cell’ models (Figure S9) and constructed two additional ‘one-virus-one-cell’ models as controls (Figure S10). Stable Jurkat cell pools were generated in two sequential steps following the schematic in Figure 3B.

**Figure 3:**
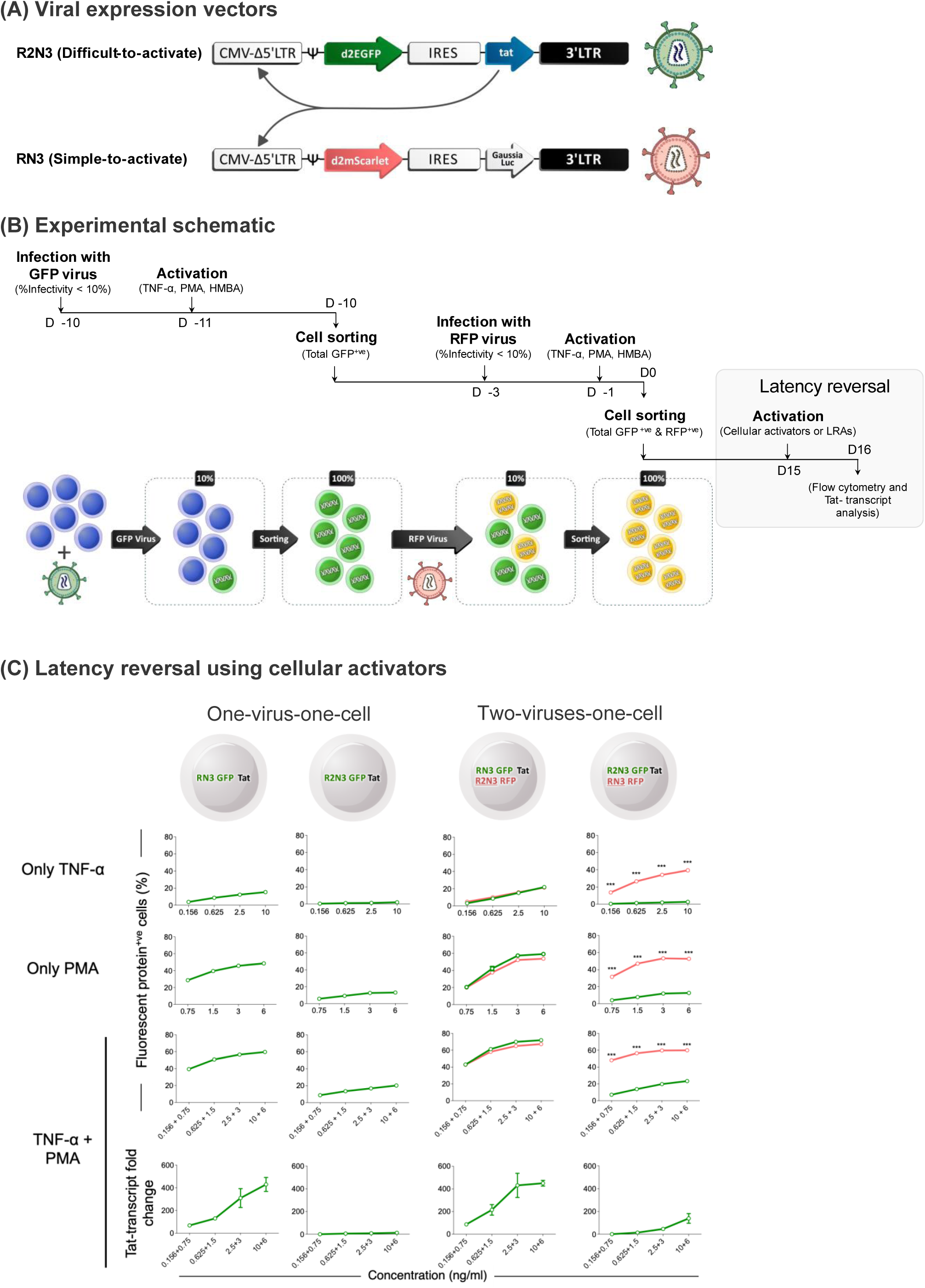
Latency reversal kinetics of RN3- and R2N3-LTRs using cellular activators in the two-viruses-one-cell models. **(A)** The sub-genomic viral vectors, where each LTR encodes a different fluorescent protein. Note that only one of the vectors encodes Tat under an IRES element. **(B)** The experimental workflow and a schematic of sequential infection and selection of cells are presented. Jurkat T-cells were first infected with the virus co-expressing d2EGFP and Tat. GFP+ve cells were sorted and re-infected with the d2mScarlet-expressing virus and selected for RFP expression. The yellow-colored cells represent the dually infected cell pool. **(C)** Latency reversal kinetics of the RN3- and R2N3-LTRs. The left panels represent mono-infections with RN3 or R2N3 strains, each co-expressing d2EGFP and Tat. The right panels represent the two-viruses-one-cell models. In one cell model, Tat is encoded by the RN3-LTR, and in the other by the R2N3-LTR. The activation status of each LTR in a cell model can be monitored by the expression status of the associated fluorescent marker. The latency reversal assays were performed in the presence of increasing concentrations and a combination of activators, TNF-α and/or PMA. The red and green curves represent the expression profiles of d2EGFP and d2mScarlet, respectively. Tat transcript levels were determined using qPCR (bottom panel). The data are representative of three independent experiments.

Cells were stimulated with increasing concentrations of TNF-α and/or PMA, and the expression of the two fluorescent reporters was analyzed by flow cytometry to assess the latency reversal profiles of each LTR. Tat transcript levels were quantified at various time points using real-time PCR (Figure 3C). In mono-infected cells with the canonical RN3 provirus, Tat and d2EGFP were co-expressed in a dose-dependent manner, correlating with the concentration of TNF-α and/or PMA (Figure 3C, left panels), and Tat transcripts were robustly induced. In contrast, mono-infection with the R2N3 strain yielded low d2EGFP expression and Tat transcript levels under identical conditions. For example, at 10 ng/ml TNF-α and 6 ng/ml PMA, RN3-infected cells showed a ∼400-fold increase in Tat transcripts over baseline, while R2N3-infected cells showed only a ∼13-fold increase. Thus, GFP expression from each LTR mirrored the Tat transcript levels, which were consistent with earlier findings. We next assessed the co-expression of the two reporters when the cells harboured both viral strains.

In the ‘two-viruses-one-cell’ model, cell activation led to robust intracellular Tat production from the RN3-LTR, efficiently reversing latency at both LTRs and resulting in strong expression of both d2EGFP and d2mScarlet (Figure 3C, right panels). In contrast, when Tat was driven by the R2N3-LTR, even the highest inducer concentrations yielded only minimal Tat, leading to weak activation. For example, 67.3 ± 0.9% of cells expressed d2mScarlet when Tat was under the RN3-LTR, compared to only 23.4 ± 0.7% expressing d2EGFP when Tat was under the R2N3-LTR. Notably, low levels of Tat were sufficient to activate the RN3-LTR encoding d2mScarlet but not the tougher R2N3-LTR encoding d2EGFP. Similar patterns were observed in mono-infected cells (Figure 3C, left panels) and under Prostratin treatment, but not with JQ1 (Figure S9, see Discussion).

Data from the two-viruses-one-cell model clearly demonstrate that duplication of the RBEIII motif substantially impairs latency reversal. Importantly, these findings highlight the central role of the HIV-1 master transcriptional circuit, comprising the LTR and Tat, in governing the ON/OFF states of viral gene expression, though the contribution of cellular environmental cues cannot be entirely excluded.

### R2 strains emerge spontaneously and dominate the latent reservoir in the chronic phase

The results from T-cell lines and primary CD4 cells indicate that adding an extra RBEIII motif to the LTR markedly suppresses latency reversal, an effect counteracted by the simultaneous addition of an NF-κB motif, as in R2N4. However, the engineered viral clones may not reflect the extensive genetic diversity of viral strains in natural infection. To address this, we analyzed the latent viral reservoir in seropositive individuals, examining the impact of co-added RBEIII and/or NF-κB motifs using cross-sectional and longitudinal samples from both acute and chronic phases of the infection.

Peripheral blood samples were collected from subjects in the chronic phase of HIV-1 infection. While some were drug-naïve at baseline, despite unknown dates of infection, others were ART-experienced. Drug-naïve individuals were initiated on ART following sample collection, and all subjects were followed longitudinally. Co-infection with the canonical RN3 and variant R2N3 or R2N4 strains was identified via LTR-genotyping using the Illumina NovaSeq platform (Figure S11). Studying latency reversal in co-infected individuals enabled the normalization of experimental variability. Cell-associated RNA (caRNA) from enriched CD4+ T cells, with or without activation, was subsequently analyzed (Figure S11).

At least five subjects were identified with co-infection involving the canonical RN3 strain and one or more R2 variants (R2N2, R2N3, and/or R2N4). The clinical profiles of these participants are summarized (Table S1). To assess preferential reactivation, we compared the inducibility of co-existing proviral strains within the latent reservoir of enriched CD4+ T cells from these individuals.

Among the five co-infected subjects, only for VFSJ15 multiple clinical samples spanning several follow-up time points were available. Samples from M0, M6, and M12 corresponded to the ART-naïve phase, while ART was initiated at M18, making M24 and M36 ART-exposed time points. In VFSJ15, at M0, only the canonical RN3 strain was detected in both proviral DNA and caRNA from enriched CD4+ T cells, irrespective of activation (Figure 4A, Table S2). The R2N4 strain first appeared at M6 and persisted thereafter in both the proviral DNA and caRNA compartments under all tested conditions. Its relative abundance in the proviral DNA progressively increased: 9.1%, 23.0%, 35.0%, and 65.9% at M6, M12, M24, and M36, respectively (Table S2). Under ‘no-activation’ conditions, caRNA proportions mirrored those of genomic DNA. At M36, RN3, R2N3, and R2N4 contributed 26.7%, 2.1%, and 71.2% of caRNA reads, respectively, indicating increasing dominance of R2N4, especially post-ART. Upon activation, R2N4 transcripts consistently predominated across all conditions at later time points. This dominance was most pronounced at M24 and M36, where nearly all caRNA reads represented R2N4. All NGS analyses were conducted in biological duplicates, which showed strong concordance, confirming the reproducibility of the findings (Figure S12).

**Figure 4:**
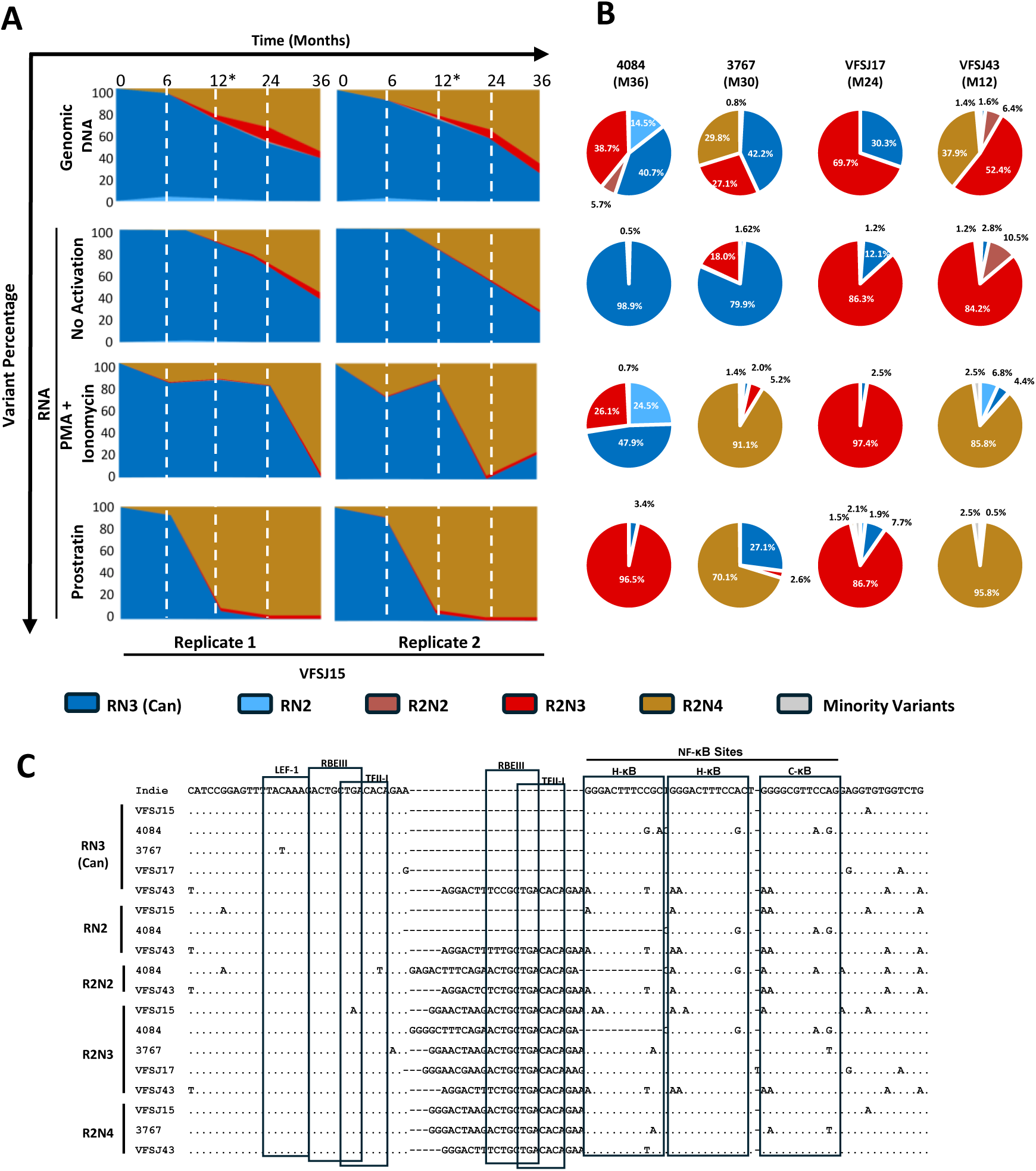
The R2 phenotype is manifested in natural infection. (A) ART administration enhances the rate of R2 switch. Stacked area plots showing the percentage prevalence of different viral variants (colour code as indicated) from the patient VFSJ15 across time points (shown as white dashed lines) in two replicate experiments. The time of ART initiation is depicted with an asterisk. (B) HIV-1C latent reservoirs harbour a higher proportion of R2 viruses after ART. Pie charts depicting the percentage prevalence of promoter variant viral strains circulating in four different patients post-ART in genomic DNA as well as cell-associated RNA. (C) Sequence Alignments of LTR variants. Representative sequences of the major LTR variants observed in the subjects of panels A and B, aligned with the HIV-1C reference sequence of Indie. Dots represent sequence identity, and dashes represent deletions. The major transcription factor binding sites are indicated.

Clinical samples from four additional ART-exposed subjects (3767, 4084, VFSJ17, and VFSJ43), all in the chronic phase, were available at a single follow-up time point (Table S1). NGS analysis of the proviral DNA revealed co-infection with the canonical RN3 and one or more R2 variants in all cases (Figure 4B, Table S3). For instance, in subject 4084, RN3, R2N3, RN2, and R2N2 accounted for 40.7%, 38.7%, 14.5%, and 5.7% of reads, respectively, indicating that R2 variants (including RN2) made up 58.9% of the total, compared to 40.7% for RN3. A similar dominance of R2 variants was observed in subjects 3767 (70.0%), VFSJ17 (69.7%), and VFSJ43 (96.7%). This trend also extended to the caRNA compartment, with or without activation, although correlations between proviral DNA and caRNA profiles were only partial. Notably, biological replicates showed strong concordance across all samples, confirming the reliability of the NGS data (Figure S12).

Collectively, our data provide key insights into HIV-1C LTR variation and latency. First, in the acute phase, as seen in subject VFSJ15, only the canonical RN3 strain is detectable in both proviral DNA and caRNA, with no evidence of R2 variants. Second, R2 strains can emerge rapidly after infection, even without ART. Third, ART appears to accelerate the emergence of R2 strains. Fourth, in the chronic phase, multiple R2 variants co-exist and tend to dominate the latent reservoir. These findings suggest that prolonged infection and/or ART promote the emergence and dominance of R2 variants in latency—a phenomenon we refer to as the ‘R2 phenotype’. The R2 phenotype is defined as the enhanced resistance of HIV-1 LTR-variant strains to latency reversal, primarily due to either a reduced number of NF-κB motifs or duplication of the RBEIII motif—with or without NF-κB reduction.

### R2 phenotype is rare in the early phase of HIV infection

Since the emergence of the R2 phenotype likely involves positive selection over time, its frequency is expected to be lower in the early/acute phase compared to the chronic phase. To test this, we analyzed stored samples from subjects in early and chronic phases, drawn from two clinical cohorts at YRGCARE, India. These individuals were p24 and viral RNA negative three months prior but tested positive at baseline while remaining negative for integrase antibodies, classifying them as early phase infection. All participants were ART-naïve at baseline, initiated on ART, and followed for 2–3 years.

Plasma and PBMC samples collected at 0, 1, and 3 months from four early-phase subjects (Table S4) were analyzed for LTR sequences in proviral DNA and cell-associated RNA (caRNA) using NGS, with all analyses conducted in biological duplicates. As expected, the canonical RN3 strain predominated in the proviral DNA of all subjects, with only minimal representation of R2 variants (Figure 5A). At baseline, RN3 constituted 92.3%, 98.0%, 94.3%, and 94.4% of total reads in AEHI001, AEHI006, BEHI001, and BEHI002, respectively (Table S5). caRNA from AEHI001 and AEHI006 also showed exclusive RN3 representation across all activation conditions. In contrast, BEHI002, and to a lesser extent BEHI001, showed caRNA reads mapping to R2 variants, particularly R2N3 and RN2, which were also detectable in the proviral DNA of BEHI002 at 4.0% and 0.5%, respectively. These R2 forms emerged primarily under activation conditions. Similarly, plasma viral RNA from AEHI001, AEHI006, and BEHI001 showed no detectable R2 variants at any time point. However, BEHI002 exhibited a high baseline R2 presence (83.1%), which declined sharply to 3.6% and 20.9% at months 1 and 3, respectively. These findings indicate that RN3 dominates in early infection, while R2 variants emerge sporadically, especially under cellular activation.

**Figure 5:**
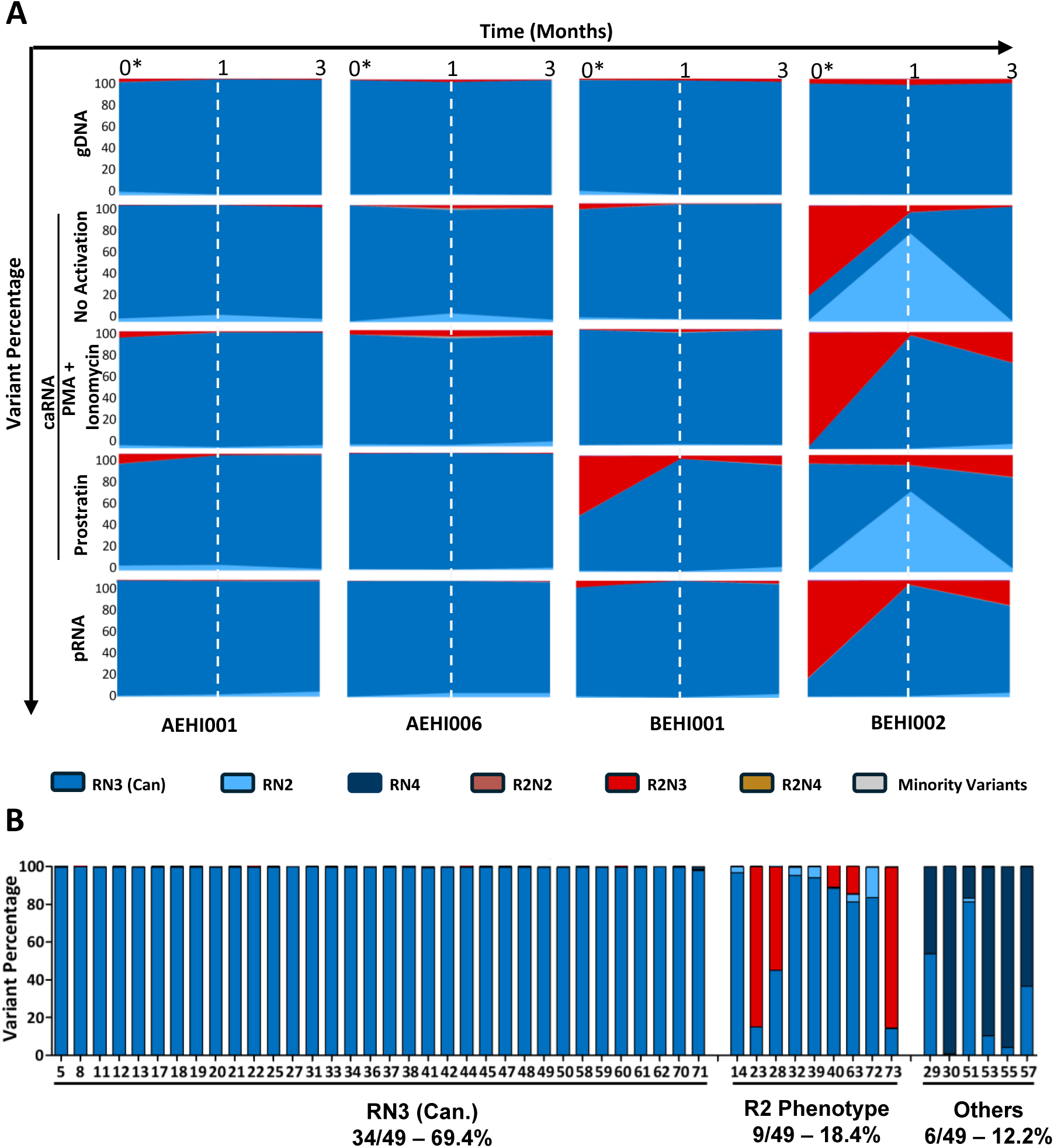
(**A) Early phase of HIV-1C infection is dominated by the RN3 viral strains.** Stacked area plots of the NGS reads of the LTR demonstrate the LTR variant forms in four HIV-1C subjects belonging to the early phase who were initiated on ART at the baseline. These subjects were drawn from the Indo-Dutch clinical cohort. **(B) R2 phenotype is seen only in a small minority of the early-phase subjects.** The NGS profile of the LTR-variant strains in 49 early-phase subjects, classified as per the major circulating variant. The colour code is consistent across all figures. caRNA: cell-associated RNA; gDNA: genomic DNA; pRNA: plasma viral RNA

To broadly estimate the frequency of R2 strains during the acute phase, we analyzed 49 cross-sectional, ART-naïve samples from the YRGCARE repository. LTR genotyping revealed that 34 of 49 subjects (69.4%) harbored only the canonical RN3 strain in their proviral DNA compartment (Figure 5B, Table S6). R2 variants were detected in nine subjects (18.4%) at varying levels, while six subjects (12.2%) showed the presence of RN4 along with RN3. These findings, consistent with both longitudinal and cross-sectional analyses, indicate that the latent reservoir in early HIV-1C infection is predominantly composed of the canonical RN3 strain, with spontaneous emergence of R2 variants in a subset of individuals.

### In the chronic phase, the R2 phenotype is manifested in approximately half of subjects

We conducted NGS analysis on four additional ART-naïve subjects in the chronic phase from the Indo-Dutch cohort using stored plasma and PBMC samples at 0, 6, and 12 months (Table S7). In CHI009 and CHI010, the proviral DNA at baseline was overwhelmingly dominated by the RN3 strain, comprising 97.6% and 96.6% of reads, respectively (Figure 6A, Table S8), with only trace levels of variant LTRs. caRNA in both subjects, across all activation conditions, was also exclusively RN3. In contrast, CHI004 and CHI008 showed a pronounced presence of R2 variants in both proviral DNA and caRNA. Notably, CHI008 exhibited high proportions of R2N3 reads—71.6%, 96.6%, and 90.8%—at 0, 6, and 12 months, respectively (Figure 6A, Table S8).

**Figure 6:**
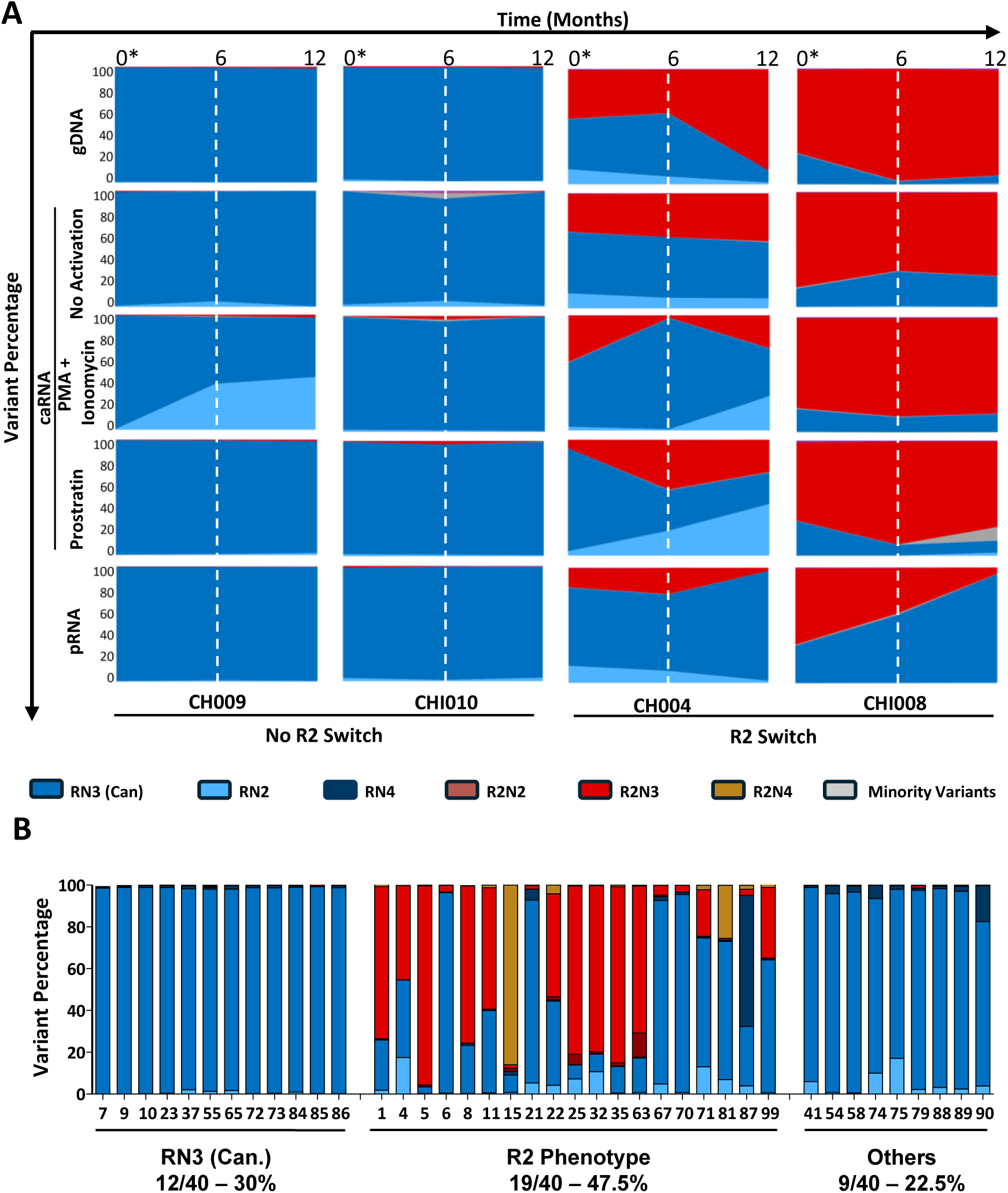
(**A) Half of Chronic HIV-1C patients demonstrate the R2 switch.** Stacked area plots of the NGS reads of the LTR demonstrating the prevalence of R2 strains in two of four chronic HIV-1C patients who were initiated on ART at the baseline. These subjects were drawn from the Indo-Dutch clinical cohort. **(B) The prevalence of the R2 Switch in ART-naïve subjects of YRGCARE belonging to the chronic phase**. The colour code is consistent across all figures.

In CHI009 and CHI010, plasma viral RNA was almost exclusively composed of the canonical RN3 strain. In contrast, although CHI004 and CHI008 harbored proviral reservoirs enriched with R2 variants, R2 RNA was detectable in plasma at lower levels. In CHI004, R2 RNA constituted 16.5% of plasma reads at baseline, increased to 22.6% at month 6, and declined to 2.5% at month 12. CHI008 showed a similar trend, with R2 reads comprising 65.6% at baseline, decreasing to 39.6% and 5.4% at months 6 and 12, respectively. These results suggest that, despite the dominance of R2 variants in the proviral reservoir, the transcriptionally active plasma virus population remains largely composed of the more readily inducible RN3 strain.

In a cross-sectional analysis of 40 ART-naïve subjects in the chronic phase (Table S9), 12 (30.0%) harbored only the canonical RN3 strain, while 19 (47.5%) carried one or more R2 variant strains. Thus, nearly half of the chronic-phase subjects exhibited the R2 phenotype. Comparatively, the prevalence of the R2 phenotype increased from 18.4% in the early phase to 47.5% in the chronic phase, indicating a time-associated enrichment of R2 variants in drug-naïve individuals.

### The R2 phenotype is associated with a larger proviral load constituting the bulk of the non-inducible HIV reservoir

Total HIV DNA was quantified by ddPCR using a TaqMan probe targeting the LTR enhancer region (Figure S14A). At baseline, R2-phenotype subjects (CHI004, CHI008) showed markedly higher HIV DNA levels than canonical R-infected subjects (CHI009, CHI010), with 770 and 28,626 vs. 572 and 222 copies per 10⁶ CD4⁺ T cells, respectively. ART led to substantial reductions in all cases, yet R2 subjects consistently retained higher DNA levels at both follow-ups. For instance, CHI004 had 338 and 420 copies at 6 and 12 months, compared to 160 and 157 in CHI010 (Figure S14B). These results suggest greater stability of latent proviruses in R2 infections.

To assess HIV-1 reservoir inducibility, we evaluated the latent reservoirs in two subjects each harboring either the canonical R or the variant R2 LTR phenotype using the Tat/rev Induced Limiting Dilution Assay (TILDA). Note that R2 phenotype is invariably associated with the co-presence of the R strain. R2-infected individuals consistently exhibited higher levels of spontaneous and stimulated viral reactivation, reflecting a larger but less inducible latent reservoir compared to R phenotype cases.

At baseline, without stimulation, R-infected individuals exhibited significantly lower frequencies of spontaneous induction events: while CHI009 and CHI010 (R phenotype) showed 10.6 and 6.92 events/million CD4⁺ T cells, respectively, CHI004 and CHI008 (R2 phenotype) showed 127.9 and 320.2 events (Figure 7A). Post-ART, frequencies declined across all subjects but remained higher in R2 cases at 6M (CHI009: 9.64; CHI010: 9.02 vs. CHI004: 9.02; CHI008: 21.04) and at 12M (CHI009: 2.57; CHI010: 8.58 vs. CHI004: 3.36; CHI008: 12.9).

**Figure 7:**
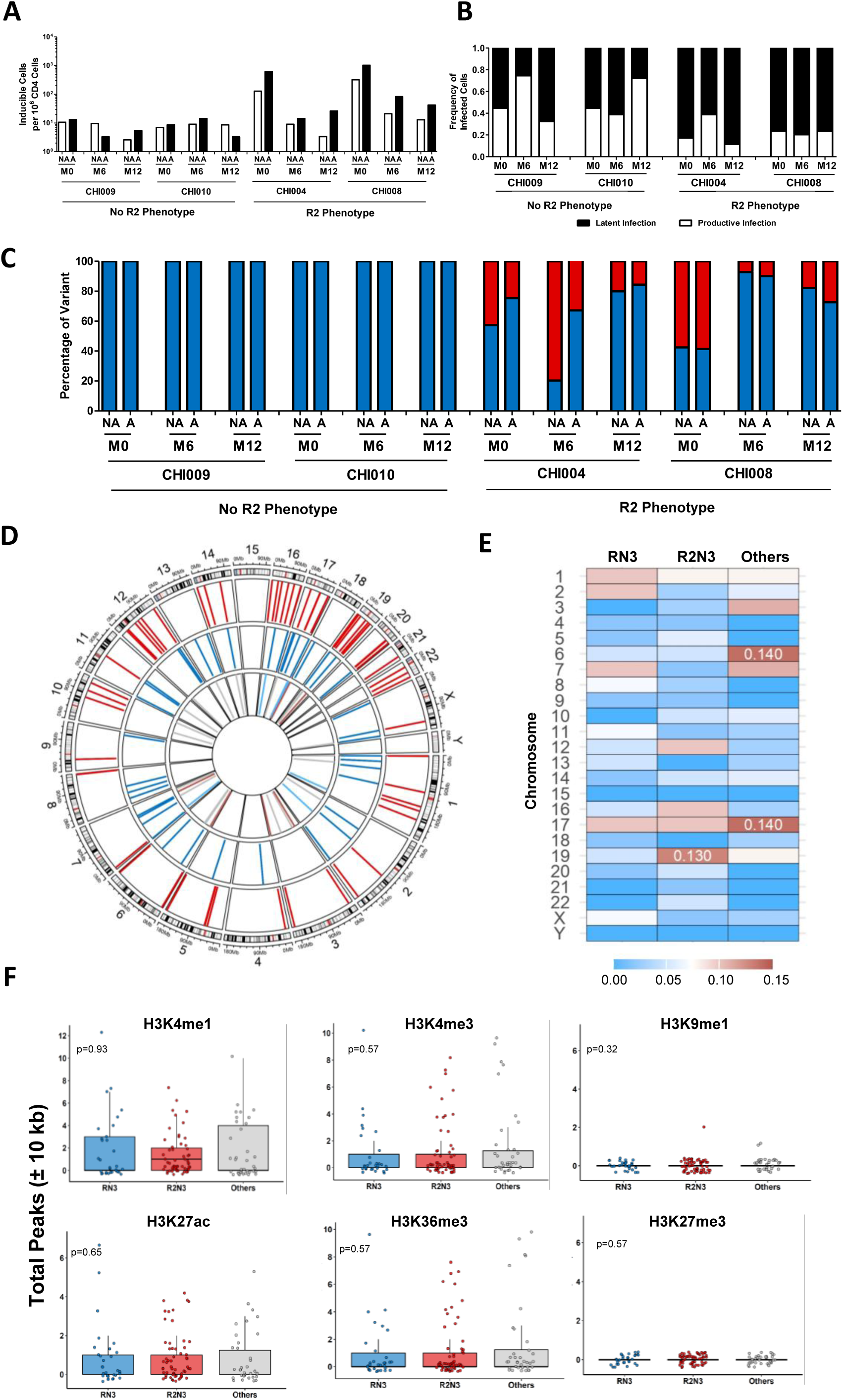
Latent reservoir analysis of R and R2 strains. **(A)** The frequency of CD4+ T cells producing msRNA spontaneously (No activation) or after 12 h of stimulation (activation) in R and R2 subjects. **(B)** Stacked bar graphs depicting the frequencies of CD4+ T cells in four subjects infected productively or latently. **(C)** Stacked bar graphs demonstrating the proportion of variants that are induced in the TILDA assay. The data are normalized with the size of the viral reservoir presented in figure 6A. **(D)** Circos plot demonstrating integration sites of viral variants from the four subjects. The three concentric rings depict integration sites of R2 (red), R (blue) and other (grey) variants, respectively. **(E)** Heatmaps depicting the proportions of integration into each chromosome compared among the LTR-variant strains. **(F)** Number of peaks for key histone marks in a 10-kb region flanking the proviral integration sites shown in (D). Data are represented as Mean± SD, and analysed using the Kruskal-Wallis Test. The colour code is consistent with previous figures.

Stimulation with PMA-ionomycin markedly increased inducibility. At baseline, CHI004 and CHI008 had 614.62 and 1028.8 events/million, versus 13.11 and 8.58 in CHI009 and CHI010. This trend persisted at 6M (CHI004: 14.25; CHI008: 82.48 vs. CHI009: 3.3; CHI010: 14.25) and 12M (CHI004: 26.1; CHI008: 42.17 vs. CHI009: 5.38; CHI010: 3.3). Sporadic deviations occurred, with spontaneous activation exceeding induced responses in CHI009 at 6M (9.64 vs. 3.3) and CHI010 at 12M (8.58 vs. 3.3). On average, baseline spontaneous activation in R phenotype subjects (mean: 8.76) was ∼25-fold lower than in R2 cases (mean: 224.05). Although ART reduced this gap, it persisted at 6M (R: 9.33 vs. R2: 15.03) and 12M (R: 5.58 vs. R2: 8.16). Under stimulation, the disparity was greater, R2 subjects showed ∼60-fold higher inducibility at baseline (R2: 821.71 vs. R: 13.68), with differences maintained at 6M (48.36 vs. 8.77) and 12M (34.17 vs. 4.34). Productive-to-latent infection ratios, derived from TILDA data (Figure 7B), were stable over time but differed by phenotype. R subjects (CHI009, CHI010) showed a near 1:1 ratio (0.51:0.49 and 0.52:0.48), while R2 subjects (CHI004, CHI008) had skewed ratios (∼1:3.5), indicating that most proviruses in R2 infections remained latent even with stimulation.

We further characterized LTR variant contributions to inducibility, both absolute and relative, via LTR-PCR and next-generation sequencing on TILDA-positive wells. In R subjects, only the RN3 promoter was detected (Figures 7C, S15). In R2 subjects, CHI004 showed near-equal induction of RN3 and R2N3 (R:R2 = 1:1.08), while CHI008 showed a 3.5-fold excess of R2 transcripts. These proportions were consistent across spontaneous and induced conditions. Relative inducibility analysis, normalized to reservoir size (Figure 6A), revealed a disproportionate contribution from RN3 despite a smaller reservoir size. In CHI004 and CHI008, R:R2 transcript ratios shifted from 1:1.08 to 1.78:1 and 1:3.5 to 2.36:1, respectively (Figure 7C). This pattern held across spontaneous and induced settings.

In summary, these findings indicate that although R2 transcripts are more abundant in absolute terms in individuals with the R2 phenotype, this effect is primarily due to the relatively larger size of their reservoir. The actual number of R2-infected cells that are induced is relatively modest in comparison to the transcripts that arise from the much smaller pool of RN3-infected cells (Figures 7B, 7C, S15).

### The site of integration and epigenetic landscape do not influence the R2 phenotype

To assess whether the host chromatin landscape influences the R2 phenotype, we mapped proviral integration sites from the four chronic-phase subjects described above. Genomic DNA was sonicated, adapter-ligated, PCR-amplified, and subjected to Nanopore sequencing. A total of 139 unique sites were identified: 41 for RN3, 62 for R2N3, and 36 for minority variants (R2N2, RN2) (Figure 7D). Integration sites were widely distributed across the genome, with no significant differences among variants. Consistent with prior reports, integrations were enriched on chromosomes 16, 17, and 19 and absent on chromosomes 15 and Y (Figures 7D, 7E).

Further, to evaluate the impact of local chromatin context, we analyzed ChIP-seq data from primary CD4⁺ T cells, available in the ROADMAP database, focusing on six histone marks: H3K4me1, H3K4me3, H3K9me1, H3K27ac, H3K36me3, and H3K27me3. No variant-specific enrichment was detected near integration sites (Figure 7F). Likewise, no correlation was found between integration orientation relative to the nearest transcription start site and variant type (Figure S16). Collectively, these results suggest that the promoter configuration primarily drives the enhanced transcriptional silencing characteristic of R2 variants in clinical samples.

## Discussion

### The TFBS profile of the LTR influences viral transcription in contrary ways

Our study demonstrates that subtle variations in the TFBS profile of the HIV-1 LTR significantly influence viral transcription and latent reservoir stability of LTR-variant strains. Notably, the LTR and Tat alone can govern ON/OFF transcriptional decisions, independent of other viral elements, highlighting the functional impact of strain- or subtype-specific sequence differences. Here, we investigated HIV-1C-specific TFBS variants and their effects on latency regulation.

Our data support the model that the TFBS landscape of the HIV-1 LTR controls transcriptional activation or silencing through context-dependent recruitment of host factors. As the sole essential cis-acting element, the LTR serves as a scaffold for both activating and repressing complexes, with transcriptional outcomes shaped by the specific TFs bound, their interactions, and regulatory modifications. The bimodal ON/OFF transcriptional function of the LTR, without intermediate states, requires two key conditions: TF families must include both activating and repressing members, and transitions between states must involve coordinated remodeling of TF complexes to ensure all TFBS function in concert toward a single transcriptional fate. The major TF families, including TCF-α/LEF-1, RBF-2, AP-1, NF-κB, NFAT, and Sp1, satisfy these conditions.

For instance, NF-κB motifs enhance transcription when bound by p50:p65 heterodimers during cellular activation ^25^ but repress it when occupied by p50 homodimers that recruit HDAC1 ^26^, Bcl-3 ^27^, or CtBP ^28^ in resting cells. Additionally, NFAT binds the 3’ end of some NF-κB motifs, further modulating transcription, especially in primary CD4+ T cells ^29^.

Similarly, the RBEIII motif upstream of the enhancer is bound by the RBF-2 complex (USF1, USF2, TFII-I) ^30^ ^31^ ^32^. RBEIII overlaps a 7-bp AP-1 site (5’-TGACACA-3’) ^33, 34^, facilitating cooperative AP-1–RBF-2 interactions. In resting cells, hypo- or non-phosphorylated RBF-2 represses transcription via chromatin modification, while activation-induced phosphorylation enables transcription ^30^. TFII-I, a key RBF-2 component, also modulates host gene expression contextually, including transcriptional repression ^35^. AP-1 family members (e.g., c-Jun, Fos) bind sequences overlapping RBEIII and interact with adjacent NF-κB, NFAT, or RBF-2 motifs. Their transcriptional effects depend on dimer composition and surrounding TFBS context ^36^. Similarly, Sp1 elements in the core promoter can either activate (via Sp1-dominant complexes) or repress (via Sp3-containing complexes) transcription ^37^.

### A fine balance exists between the transcription-enhancing and -inhibiting functions of regulatory elements in the LTR

We show for the first time that the R2 genotype, duplication of the RBEIII core motif, and flanking MFNLP sequence strongly represses transcription in HIV-1C (RN3 vs. R2N3), promoting deeper latency. In contrast, duplicating only the NF-κB motif and flanking regions (RN3 vs. RN4) enhances transcription, while co-duplication of both elements (RN3 vs. R2N4) balances their individual effects. These findings suggest that NF-κB and RBEIII motifs function as activators and repressors, respectively, in a cell activation-dependent manner. Our data further indicate that a precise NF-κB: RBEIII motif ratio, three: one in canonical HIV-1C, is critical for establishing, maintaining, and reversing latency. Disrupting this balance alters transcriptional dynamics, explaining how the addition of an extra RBEIII motif without NF-κB co-addition (R2N3) increases resistance to rebound.

Strong support for this model comes from testing constructs with constant RBEIII duplication (R2) and varying NF-κB motif numbers (1-4). As NF-κB copies decreased, reactivation from latency declined, with R2N2 and R2N1 showing no reactivation (Figure 2). Importantly, HIV-1 variants with differing NF-κB motif numbers occur naturally ^10^. Collectively, our findings position the HIV-1 LTR as a transcriptional control hub, balancing opposing TF forces in response to cell activation and TF availability.

### Why is the R2 phenotype highly suppressive for HIV-1C transcription?

Although the MFNLP duplication, encompassing the RBEIII core motif and flanking sequences, has been reported across multiple HIV-1 subtypes, including HIV-1C ^10^, its biological effects have been primarily studied in the context of transcription in HIV-1B ^38^ ^39^ ^12^ ^40^, with limited insight into its role in latency. Reports on the impact of MFNLP on transcription are inconsistent: some showed reduced activity upon RBEIII duplication ^38^ ^41^ ^42^ ^43^, while others observed no effect ^44^ ^12^ ^45^or even enhancement ^46^ ^47^. These discrepancies likely reflect differences in experimental systems and the sequence heterogeneity of RBEIII-cluster duplications. In contrast to previous reports, we show that the R2 phenotype in HIV-1C exerts strong and reproducible transcriptional suppression across several different viral formats. Patient-derived CD4+ T cell reactivation assays further validate this suppression, underscoring the biological relevance of the R2 phenotype in HIV-1C.

The RBEIII core motif (5′-ACGTCGTA-3′), located upstream of the enhancer, is thought to activate or repress transcription depending on the activation status of the T cell ^48^. The RBF-2 complex, comprising USF1, USF2, and TFII-I, binds this motif and adjacent bases ^30^. Components of RBF-2 can act as transcriptional activators or repressors. USF1/2 heterodimers are generally associated with repression of specific cellular genes ^49^ ^50^ ^51^ ^52^. While USF can activate HIV-1 transcription in Jurkat cells, it suppresses it in HeLa cells regardless of viral subtype ^53^, though epithelial cell models may not fully reflect HIV latency.

TFII-I is particularly suppressive in certain contexts, recruiting HDAC1/3 to cellular promoters^35^ ^54^. Among its splice variants, TFII-Iβ is especially repressive at the c-fos promoter in uninduced fibroblasts ^55^. HDAC3 may augment repression by recruiting HDAC1, 4, 5, and 7 into multi-protein complexes ^56^ ^57^. TFII-I also recruits additional co-repressors, including LSD1 and polycomb repressor complex (PRC) components ^58^ ^59^. Thus, in unstimulated cells, RBF-2 may assemble a potent transcriptional repressor complex.

Despite detailed characterization of R2 strain latency kinetics, the exact mechanism of their resistance to latency reversal is yet to be determined. Variants like R2N3 and R2N4 differ from the canonical RN3 promoter not only by RBEIII motif duplication but also by subtle changes in NF-κB site spacing and flanking sequences. The conserved RBEIII motif and its association with RBF-2-mediated repression suggests a role, but current data do not support it as the sole cause. Altered NF-κB positioning may affect cooperative binding of p50:p65 or p50:HDAC complexes, while flanking sequences may influence chromatin structure or nucleosome positioning. Given the potential for small sequence changes to impact transcription factor binding and nucleosome occupancy, the latency resistance likely stems from combined, interdependent sequence alterations. Further analysis of the TF complexes associated with R2 and non-R2 LTRs is warranted. Additionally, given the subtype-specific differences in RBEIII flanking regions (Figure S18), evaluating the R2 phenotype’s suppressive effect across different HIV-1 subtypes is essential.

### NF-κB can function as a pioneer transcription factor

Despite its high thermodynamic propensity for nucleosome formation, the ∼150 bp DNase Hypersensitive Site-1 (DHS1) of the LTR, containing TFBS, remains nucleosome-free or bound by a low-affinity nucleosome in proviral DNA. BAF, a SWI/SNF chromatin remodeler, can reposition Nuc-1 downstream, blocking the transcription start site (TSS) while keeping NF-κB motifs accessible in a transcriptionally repressed LTR ^60^. Notably, NF-κB functions as a pioneer transcription factor, capable of binding target DNA within nucleosomes or heterochromatin and initiating chromatin remodeling ^61^.

For example, in unstimulated dendritic cells, c-Rel binds inactive inflammatory gene promoters, such as *IL12b* and *Mdc*, recruits the histone demethylase Aof1, and removes the repressive H3K9me2 mark, allowing further TF binding, including NF-κB ^62^. In vitro, NF-κB can unwrap nucleosomes by 30–50 bp ^61^, displace the H1 linker histone, and invade unstable nucleosomes ^63^, enhancing chromatin accessibility. Thus, NF-κB binding sites in DHS1 remain accessible even in suppressed proviruses and are poised to trigger viral reactivation. This shift in TF occupancy correlates with RBF-2 upregulation via kinase signaling ^64^. We propose that RBEIII establishes a repressive TF profile in resting cells, whereas NF-κB promotes transcriptional activation upon stimulation, as represented schematically (Figure S18).

This model is further supported by leads obtained with latency-reversing agents. The R2N3-LTR, typically resistant to activation, was reactivated as efficiently as RN3-LTR when treated with JQ1 plus Prostratin but not with PMA and TNF-α (Figure 2B, C). PMA and TNF-α activate NF-κB-dependent signaling that facilitates P-TEFb-Tat interaction by destabilizing the P-TEFb-BRD4 complex ^65^. Their efficacy depends on how robustly cell surface signaling is induced. In contrast, JQ1 bypasses cellular signaling by directly releasing P-TEFb from the 7SK snRNP complex. This suggests that R2 LTRs are blocked at the level of P-TEFb recruitment, a step highly dependent on signaling through transcription factors.

Based on these observations, we propose that R2 LTRs are less responsive to cellular activation due to tighter chromatin packaging, likely from enhanced HDAC recruitment or occlusion of NF-κB sites. Consequently, R2 variants fail to recruit the transcriptional machinery via natural signaling cues but remain susceptible to chromatin-modifying agents like JQ1.

### Can the R2 phenotype confer a clinical benefit to subjects?

The R2 phenotype, which promotes a stable and non-reactivating latent reservoir, may lower plasma viral load, slow CD4+ T cell decline, and reduce T-cell exhaustion, potentially delaying disease progression. However, due to the limited clinical sample size, we could not assess this directly. The observed positive association between the R2 genotype and ART exposure raises the possibility of clinical benefits, especially in patients lost to follow-up (LFU) after receiving ART. LFU remains a significant barrier to HIV care, affecting 25–80% of patients, particularly in resource-limited regions like India and even in high-income countries ^66^ ^67^ ^68^ ^69^ ^70^. While viral rebound and drug resistance are known LFU outcomes, our findings suggest the R2 genotype as an alternative pathway that warrants further exploration.

This is particularly relevant in the context of post-treatment controllers (PTCs), individuals who maintain viral suppression after ART discontinuation ^71^ ^72^ ^73^. PTCs, comprising 5–12% of ART-treated individuals, have been documented in diverse cohorts, including US, European^74^, and African populations ^75^ ^72^. Unlike elite controllers, PTCs typically require early ^76^ ^71^ ^73^ and/or prolonged ^77^ ART.

Latency stability in PTCs is associated with smaller reservoir size, viral defects, and integration site chromatin states ^78^ ^79^ ^80^. Viral integration into euchromatin facilitates reactivation under selective pressure, while over time, reservoirs shift toward more stably silent proviruses in heterochromatin. Although ART does not reduce reservoir size, it promotes reservoir stabilization via chromatin and epigenetic changes ^81^ ^82^. PTCs also show enhanced immune function, including stronger T-cell responses and reduced exhaustion ^83^ ^72^, though these features remain incompletely characterized ^74^.

Importantly, the latency reversal resistance observed in R2 variants fundamentally differs from that in PTCs. In PTCs, latency is shaped by extrinsic factors, such as the viral integration site and chromatin context, while in R2 variants, it is governed by intrinsic changes in TFBS architecture within the LTR, although epigenetic factors may still contribute. Notably, in co-infected subjects, we found no significant differences in integration site profiles or epigenetic marks between RN3 and R2 strains (Figure 7D - F), indicating that extrinsic factors do not drive resistance in R2. Additionally, while PTCs typically require early or prolonged ART, our cohort comprised ART-naïve individuals in the chronic phase, reinforcing the conclusion that the R2 phenotype represents a distinct mechanism of latency stabilization.

Given the dominance of HIV-1C in India, we could not assess whether the R2 phenotype similarly affects latency in other subtypes. Experimental studies are needed to determine whether subtype-specific variations in sequences flanking the RBEIII motif (Figure S19) modulate the observed resistance to latency reversal.

### The transcription ON/OFF decision-making appears to be hard-wired into the viral master regulatory circuit

HIV-1 latency was initially attributed to the transition of infected CD4⁺ T cells into a quiescent state ^21^. Over time, two main models emerged to explain latency: deterministic and stochastic^84^. While latency has been associated with factors like cellular activation, promoter methylation, and integration into transcriptionally silent regions ^21^ ^85^ ^86^ ^87^, these deterministic mechanisms fail to fully explain latency in patient-derived CD4⁺ cells ^88^. Instead, increasing evidence supports a stochastic model of latency regulation ^24^ ^5^.

The HIV-1 variants identified in this study provide a powerful model to dissect the relative roles of viral and cellular factors in latency regulation. By using LTRs with distinct activation thresholds, we normalized for external influences. This unique ‘two-viruses-one-cell’ system enabled us to investigate the intrinsic function of the viral master transcription-regulatory circuit (MTRC), comprising the LTR and Tat. Our findings suggest that the ON/OFF transcriptional state is primarily virus-driven (Figure 3), though environmental factors may still play a significant role.

Given the distinct but contrasting biological advantages of R2N3 and R2N4, it is difficult to predict which may ultimately outcompete the canonical RN3 strain if that happens. The R2N4 variant likely confers a selective edge through rapid latency establishment and stable transcriptional silencing via the added RBEIII motif, combined with efficient reactivation due to an extra NF-κB motif, potentially enabling higher virion production upon activation. In contrast, R2N3 may gain evolutionary traction through its elevated activation threshold, promoting durable latency and favoring clonal expansion via homeostatic proliferation, like HTLV-1 ^89^. Further studies are needed to define the evolutionary trajectories of these emerging variants.

### Summary

This study reveals that duplication of the RBEIII motif in the HIV-1 LTR significantly influences latency establishment, maintenance, and resistance to reactivation, particularly when contrasted with NF-κB motif duplication, which enhances transcription. These findings highlight two opposing regulatory hubs within the LTR: an activation-promoting NF-κB cluster and a latency-enforcing RBEIII cluster. Despite a limited sample size and unaccounted variation in motif positioning, the data suggest that ongoing LTR evolution may select for architectures that confer a latency advantage. Importantly, under suppressive ART, we identify a novel mechanism of proviral enrichment, distinct from integration site bias, where the virus itself acquires cis-acting regulatory changes that stabilize latency. This suggests that HIV-1 can evolve promoter configurations that resist transcriptional reactivation, reflecting an adaptive trajectory toward deeper, more persistent latency.

## Materials and methods

### Construction of viral vectors and panels of promoter-variant viral strains

We constructed three analogous panels – sub-genomic-reporter, near-full-length-reporter, and full-length - of four LTR-variant HIV-1 strains - RN3, RN4, R2N3, and R2N4. Additionally, we also constructed the sub-genomic reporter panel contains R2N2- and R2N-LTRs. The strains of each panel differed from one another in the number of copies of the RBEIII (R, R vs. R2) and NF-κB (N, N3 vs. N4) motifs they contained in the LTR (Figure S2 and S6). For example, the RN3-LTR, the canonical viral promoter, contained a single RBEIII motif and three copies of the NF-κB motifs. Except for the defined difference in the TFBS composition, the viral vectors of a panel are identical in the viral vector backbone. The 3’ LTRs of all three parental vectors were engineered to construct the TFBS-variant viral panels.

The sub-genomic viral vector LTR-d2EGFP-IRES-Tat (p932.RN3) was modified from a previously reported pcLdGIT backbone ^90^ that co-expresses GFP and Tat. The pcLdGIT vector contains the 3’-LTR of Indie.C1 molecular clone (Accession no. AB023804.1), a short-lived GFP reporter (d2EGFP), and Tat of HIV-1C (BL4-3, Accession no. FJ765005.1) expressed under an IRES element. The pcLdGIT vector lacked the polypurine tract (PPT). To generate p932.RN3, its 3’LTR was replaced using XhoI and PmeI sites with a PCR-amplified LTR using primers N2685 and N2690 (Supplementary file 2), where the forward primer (N2685) introduced the 23 bp PPT upstream of the 3’LTR in the final construct. Compared to pcLdGIT, p932.RN3 vector containing the restored PPT element demonstrated ∼10 times superior GFP MFI (Data not shown).

The RN4-, R2N4-, R2N3-, R2N2-, and R2N-LTRs were constructed as follows. Each vector contains a unique restriction enzyme (RE) site at the end of the 3’-LTR, introduced via the reverse primer, for unambiguous identification (Supplementary file 2). p932.RN4 vector was generated by amplifying the 3’-LTR from p902.FHHC vector ^90^ using primers N2685 and N2691 and replacing the 3’LTR of p932.RN3 via AscI and SacII digestion.

The R2N4-LTR was created by amplifying RN3-LTR with the primers N2886 and N2692 to introduce a duplication of the κB-like (N) motif and RBEIII cluster (R), along with a spacer (5’-GAAGGGACTTTCAAGACTGCTGACACA-3’), then replacing the 3’-LTR in p932.RN3 using BspEI and SacII enzymes.

For p932.R2N3 vector, and intermediate p932.RN2 vector was first generated by deleting the first NF-κB motif (5’-GGGACTTTCC-3’) and its downstream spacer (5’-GCT-3’) from RN3-LTR using the primer pair N2881-N2693, followed by RN2-LTR cloning using BspEI and SacII enzymes. The R2N3-LTR was then constructed by duplicating the κB-like (N) motif, RBEIII cluster (R), and a spacer (5’-GAAGGGACTTTCAAGACTGCTGACACA-3’) into RN2-LTR using primers N2888 and N2693 and cloning the final R2N3-LTR into p932.RN3 with BspEI and SacII; thus, creating the final p932.R2N3 vector.

The R2N2-LTR was created using overlap PCR with primer pairs N4283-N4284 (Round 1), N4285-N2889 (Round 2), and N4283-N2889 (Round 3), then inserted into p932.RN3 with BspEI and SacII sites, thus resulting in p932.RN2 vector.

Lastly, the R2N-LTR was amplified from p932.RN2 using primers N4286-N2691, where primer N4286 deleted the NF-κB motif (5’-GGGACTTTCC-3’) upstream of the last NF-κB motif (5’-GGGGCGTTCC-3’) (Figure S6). This fragment replaced the RN3-LTR in p932.RN3 using BspEI and SacII to generate p932.R2N.

Note that R2N4, R2N2, and R2N variations are found in natural infection. All viral vectors were verified by Sanger sequencing, the d2EGFP expression in HEK293T cells, and virus infectivity in Jurkat cells and primary CD4 T-cells.

A full-length virus panel was generated using the p1000.RN3 vector. First, the p1000.RN3 vector was generated by inserting a nine bp sequence into the Nef reading frame between amino acid residues 90 and 94, thus grafting a recognition site for AscI. This manipulation inserted three additional amino acids into Nef immediately upstream of the PPT element. The AscI site was introduced using an overlap PCR using primer pairs N2710-2711 for Round 1 and N2712-N2713 for Round 2. The final overlap product was amplified using primers N2710 and N2713 (Supplementary file 2). The PCR product was introduced into the pIndie.C1 vector using PacI and SacII. p1000.RN3 remains infectious and encodes all the viral proteins. Next, two additional promoter-variant viral vectors (R2N3 and R2N4) with a variable number of RBEIII (R) and NF-κB (N) motifs were constructed in the full-length viral backbone. The RE sites, AscI and SacII, flanking the 3’-LTR were used to transfer the 3’-LTRs from the p932.R2N3 and p932.R2N4 vectors of the sub-genomic virus panel to construct a complete panel of variant LTRs in the full-length backbone. The panel of full-length promoter-variant viral vectors was used to evaluate replication kinetics in PBMC.

The near-full-length virus panel was constructed using the p1001.RN3 vector (unpublished, derived from p1000.RN3) encodes the reporter protein d2EGFP and all viral proteins except the envelope. In p1001.RN3 vector, a portion of the gp120 coding region was replaced with d2EGFP. The p1001.RN3 vector was constructed in two steps. First, a point mutation in codon 25 of gp120 was introduced into p1000.RN3 vector to create a unique SphI site at amino acid positions 24-25, generating the intermediate vector pIndie.C1.AS.RN3. The SphI site was introduced by overlap PCR using primer pairs N2710 and N3082 (Round 1) and N3083 and N3084 (Round 2). The final overlap product (Round 3) was amplified using N2710 and N3084 (Supplementary file 2). The PCR product was introduced into the p1000.RN3 backbone using PacI and StuI. Second, the d2EGFP gene was amplified from p932.RN3 plasmid using the primer pair N3085 and N3086 (Supplementary file 2). The envelope sequence between SphI and StuI of the intermediate vector pIndie.C1.AS.RN3 was replaced by d2EGFP, creating the final p1001.RN3 vector. Next, three promoter-variant viral vectors (RN4, R2N3, and R2N4) were generated by transferring the 3’-LTR of p1001.RN3 using AscI and SacII from the p932 panel, thus generating corresponding members of the p1001 LTR panel. The panel of near-full-length promoter-variant viral strains was used to examine the latency establishment kinetics in the Jurkat T-cells and primary CD4 T-cells.

### Cell-culture

Jurkat T-cells were maintained in RPMI 1640 medium (Cat. No. R4130, Sigma-Aldrich, USA) supplemented with 10% fetal bovine serum (FBS) (Cat. No. 04-121-1A, Life Technologies, India), 2 mM Glutamine (Cat. No. G8540, Sigma-Aldrich, USA), 100 units/ml Penicillin G (Cat. No. P3032, Sigma-Aldrich, USA) and 100 g/ml Streptomycin (Cat. No. S9137, Sigma-Aldrich, USA). The human embryonic kidney cell lines HEK293T cells were cultured in Dulbecco’s modified Eagle’s medium (Cat. No. D1152, Sigma-Aldrich, USA) supplemented with 10% FBS, 2 mM Glutamine, 100 units/ml Penicillin G, and 100 μg/ml Streptomycin. All the cells were incubated at 37°C in the presence of 5% CO2.

### Preparation of the viral stocks

Sub-genomic reporter viral strains: Pseudotyped reporter viral strains of the sub-genomic panel were generated in HEK293T cells by co-transfecting each viral vector along with the third-generation lentiviral packaging vectors using the standard calcium phosphate protocol ^91^. Briefly, a plasmid DNA cocktail consisting of 10 μg of the viral, 5 μg psPAX2 (Cat. No. 11348; NIH AIDS reagent program, USA), 3.5 μg pHEF-VSVG (Cat. No. 4693; NIH AIDS Reagent Program, USA) and 1.5 μg pCMV-Rev (Cat. No. 1443; NIH AIDS Reagent program, USA) was used to transfect HEK293T cells at 40% cell confluency in a 100 mm dish. pTdTomato (0.05 μg) was used as a transfection control. Six hours post-transfection, the medium was replenished with DMEM. Culture supernatants were harvested at 48 h post-transfection, centrifuged at 1,000 rpm for 5 min, and stored in 0.5 ml aliquots in a deep freezer for future use.

Near-full-length reporter viral strains: The near-full-length viral strains were generated similarly to the sub-genomic viruses, except that the packaging vectors psPAX2 and pCMV-Rev were not used during the transfection. A pool of 15 μg of the viral vector (promoter-variant viral vectors mentioned in Figure 1 B) and 5 μg pHEF-VSVG was used to transfect HEK293T cells at 40% cell confluency in a 100 mm dish. Further protocol for the harvest and storage of the viruses is as described in the previous section (Preparation of viral stocks: Sub-genomic reporter viral strains).

Full-length viral strains: The promoter-variant full-length viral strains were generated by transfecting the HEK293T cells with the individual viral vector using the standard calcium phosphate protocol. A plasmid DNA of each variant vector (20 μg) (promoter variants as depicted in Figure 1 B) was transfected individually in a 100 mm dish seeded with HEK293T at 40% cell confluence. pTdTomato (0.05 μg) was used as a transfection control. Further protocol for the harvest and storage of the viruses is as described in the previous section (Preparation of viral stocks: Sub-genomic reporter viral strains).

### Estimation of relative infectious units (RIU) of two panels of viral vectors

The viral strains of the sub-genomic and near-full-length panels encode d2EGFP, permitting the titration of viral stocks using flow cytometry. To this end, 3×10^5^ Jurkat cells in each well of a 12-well tissue culture plate were infected with viral stocks serially diluted 5-fold (from 10 to 250xd) in a total volume of 1 ml of the RPMI 1640 medium containing 25 μg/ml of DEAE-Dextran and supplemented with 10% FBS, 2 mM Glutamine, 100 units/ml Penicillin G, and 100 g/ml Streptomycin. Six hours post-infection, the cells were washed and replenished with 1 ml of the RPMI 1640 medium supplemented with 10% FBS, 2 mM Glutamine, 100 units/ml

Penicillin G, and 100 μg/ml Streptomycin. After 48 h of infection, the cells were activated with a cocktail of 5 ng/ml of PMA (Cat. No. P8139, Sigma-Aldrich, USA), 10 ng/ml of TNF-α (Cat. No. T0157, Sigma-Aldrich, USA), and 5 mM of HMBA (Cat. No. 224235, Sigma-Aldrich, USA). After 24 h of activation, the percentage of GFP^+ve^ cells was analyzed by flow cytometry (BD FACSAriaIII sorter, BD Biosciences, USA). Following this, we constructed titration curves and determined the 5-10% infectivity range of the cells by regression analysis, which would correspond to ∼0.05-0.1 RIU. The flow cytometry data were analyzed using FCS Express versions 4 and 6 (De Novo Software, USA).

### The kinetics of latency establishment in Jurkat T-cells

Jurkat T-cells were infected with individual viral strains and sorted by GFP expression. Pseudotyped viral stocks of promoter variants (Figure S2 and S6) were added to Jurkat T-cells (6×10^5^) at an RIU of ∼0.05-0.1. Jurkat T-cells were present in a total volume of 2 ml of complete RPMI 1640 medium supplemented with 25 μg/ml DEAE-Dextran, 10% FBS, 2 mM Glutamine, 100 units/ml Penicillin G, and 100 μg/ml Streptomycin in a 6-well tissue culture dish. Six hours post-infection, the cells were washed and replenished with 2 ml of RPMI 1640 medium supplemented with 10% FBS, 2 mM Glutamine, 100 units/ml Penicillin G, and 100 g/ml Streptomycin and maintained under standard cell-culture conditions. The infected cell pools were expanded over seven days and then induced with a cocktail of T-cell activation agents (5 ng/ml PMA + 10 ng/ml TNF-α + 5 mM HMBA). After 24 h of activation, GFP^+ve^ cells were sorted using a cell sorter (BD FACSAriaIII cell sorter, BD Biosciences, USA). An aliquot of the sorted cell population was validated by flow cytometry to confirm the purity of the GFP^+ve^ cells. Typically, >88-90% of cells were GFP^+ve^ following flow sorting. The sorted cell pools were divided into three groups each. Individual fractions were cultured in the absence of an activator or the presence of TNF-α (10 ng/ml) or PMA (5 ng/ml), and the GFP expression was monitored by flow cytometry up to 10 days at 2-day intervals. The sorted cell pool with stable GFP expression represented a random population of proviral integrants.

### Evaluation of latency reversal in Jurkat T-cells

About 15 days following the sorting of the GFP^+ve^ cells, a maximum proportion of cells, approximately 88-96%, downregulated GFP expression depending on the TFBS profile of the variant LTR. Latency reversal assays were performed at this stage. For the assay, 2×10^5^ Jurkat cells/well seeded in a 48-well tissue culture plate were activated with a single or combination of cellular activators or LRAs. After 24 h of activation, GFP expression was monitored using a flow cytometer (BD FACSAriaIII cell sorter, BD Biosciences, USA).

### Long-term CD4 culture by Bcl-2 over-expression

We adapted the long-term primary CD4 T-cell culture model from Kim *et al.* ^18^ with technical improvisation. The model involves the transduction of primary CD4 T-cells with the B cell lymphoma 2 (Bcl-2) gene ^92^. Over-expression of Bcl-2 extends the life span of CD4 cells in culture up to 4-6 weeks, sufficient for examining HIV-1 latency. We modified the existing protocol by introducing the co-expression of mouse CD8a (mCD8a) to permit the rapid selection of transfected cells by flow sorting. The surface expression of mCD8a allows for the rapid positive selection of the infected cells using paramagnetic beads. This improvisation of the protocol shortened the duration of the experiment by at least three weeks and enhanced cell yield. CD4 T-cells isolated from the peripheral blood of healthy volunteers were activated with anti-CD3 and anti-CD28 antibodies. Activated CD4 cells were transduced with a lentivirus co-expressing Bcl-2 and mCD8a receptor. The transduced cells were positively enriched using a mCD8a commercial kit. The expression level of Bcl-2 in the infected cells is approximately tenfold higher than that of freshly isolated CD4 cells. Enriched CD4 cells overexpressing Bcl-2 were used to evaluate HIV-1 latency.

### Construction of the lentiviral vector co-expressing Bcl-2 and mCD8a

The Bcl-2 expression vector, EBFLV, and a lentiviral packaging vector, pCHelp, were a kind gift from Dr. Robert Siliciano (Johns Hopkins University, USA). In the parental pEBFLV vector, Bcl-2 expression was regulated by an EF-1α promoter. We placed the expression of mCD8a under an IRES element following the Bcl-2 gene to generate the pEBFLV-mCD8a. The engineering of the IRES-mCD8a cassette was performed as follows. The IRES element was amplified from the p932.RN3 plasmid by PCR using the primer pair N4115 and N4116 (Supplementary file 2). The mCD8a gene was amplified from the mCD8-GFP pUAST (Addgene no.17746) plasmid by PCR using the primer pair N4117 and N4118 (Supplementary file 2). The IRES-mCD8a cassette was constructed by an overlap PCR using the primer pair N4115 and N4118. Subsequently, the IRES-mCD8a cassette was introduced into the pEBFLV backbone using the Bsu36I enzyme. The expression of Bcl-2 and mCD8a from the EBFLV-mCD8a vector was verified in Jurkat T-cells and primary CD4 cells by flow cytometry.

### Preparation of the Bcl-2-mCD8a-expressing viral stocks

Bcl-2-mCD8a expressing virus was generated in HEK293T cells by transfecting the pEBFLV-mCD8a vector with the third-generation lentiviral packaging vectors using the standard calcium phosphate protocol. Briefly, a plasmid DNA cocktail consisting of 7 μg of the pEBFLV-mCD8a, 7 μg of pcHelp, and 7 μg of pHEF-VSVG (Cat. No. 4693; NIH Reagent Program, USA) was transfected in a 100 mm dish seeded with HEK293T at 40% cell confluence. pTdTomato (0.05 μg) was used as a transfection control. Six hours post-transfection, the medium was replaced with fresh complete DMEM supplemented with 10% FBS, 2 mM Glutamine, 100 units/ml Penicillin G, and 100 μg/ml Streptomycin. The subsequent procedures of harvesting and storing the viral stocks were as described previously.

### Bcl-2-mCD8a transduction of healthy donor CD4 cells and the selection and expansion of mCD8a+ cells

PBMC were isolated by Ficoll density gradient centrifugation of 40 ml of the peripheral blood collected from healthy donors. CD4 T-cells were enriched from the PBMC by negative selection using magnetic beads (Cat. No. 19052, Stemcell Technologies, Canada). From 40 ml of blood, we typically collected 35-85×10^6^ PBMC and 12-22×10^6^ CD4 T-cells after the negative selection. Subsequently, enriched CD4 cells were resuspended at a density of 1×10^6^ cells/ml in the RPMI 1640 medium supplemented with IL-2 (100 U/ml), 1 μg of each anti-CD3 and anti-CD28 antibodies, 10% FBS, 2 mM Glutamine, 100 units/ml Penicillin G, and 100 g/ml Streptomycin. The enriched CD4 T-cell suspension was distributed equally in a 12-well plate (2 ml of cell suspension per well). Each well was previously coated with 1 μg of each anti-CD3 and anti-CD28 antibodies, resuspended in PBS for 2 h. After 2 h of antibody coating, the wells were washed with PBS, and the enriched CD4 cell suspension was added to each well, as explained before. The cells were incubated for 72 h under the conditions of activation. Following the activation, approximately 5×10^6^ CD4 cells were transferred to a 6-well tissue culture plate and suspended at the density of 2×10^6^ cells/ml in the RPMI 1640 medium supplemented with IL-2 (100 U/ml), 10% FBS, 2 mM Glutamine, 100 units/ml Penicillin G, and 100 g/ml Streptomycin. To each well, the Bcl-2-mCD8a viral stock was added to obtain an infection rate of approximately 10-40% infected cells. The final volume of cells and virus stock was adjusted to 3 ml per well of a 6-well plate. The plate was centrifuged at 1,200 rpm for 2 h to improve transduction efficiency by spinoculation. Six hours post-transfection, the medium was replaced with the RPMI 1640 medium supplemented with 100 U/ml of IL-2, 10% FBS, 2 mM Glutamine, 100 units/ml Penicillin G, and 100 μg/ml Streptomycin. Two days post-infection, the transduced cells were enriched by mCD8a positive selection using magnetic beads (Cat. No. 18953, Stemcell Technologies, Canada). We typically recovered approximately 3-4×10^6^ cells at this level following infection and some cell death. The transduced and enriched CD4 cells were incubated for 7– 10 days in RPMI 1640 medium without IL-2 and supplemented with 10% FBS, 2 mM Glutamine, 100 units/ml Penicillin G, and 100 μg/ml Streptomycin. We observed a 1.3-1.4 log enhancement in cell numbers by the end of the expansion. At this stage, the cells may be stored in a liquid nitrogen container for later use.

### Sub-genomic HIV-1 infection of Bcl-2-CD4 cells

The expanded Bcl-2-CD4 cells were activated by the TCR engagement, as explained above. Subsequently, the cells were infected with the viral strains of the sub-genomic panel by spinoculation (1,200 rpm for 2 h) as described in section. Six hours post-transduction, the medium was replaced with the RPMI 1640 medium supplemented with 100 U/ml of IL-2, 10% FBS, 2 mM Glutamine, 100 units/ml Penicillin G, and 100 μg/ml Streptomycin. Two days post-infection, the GFP^+ve^ cells were sorted by flow cytometry (BD FACSAriaIII cell sorter, BD biosciences, USA), and the sorted cells were maintained in the RPMI 1640 medium without IL-2 and supplemented with 10% FBS, 2 mM Glutamine, 100 units/ml Penicillin G, and 100 g/ml Streptomycin. The latency establishment profile was monitored for 36 days at four- or six-day intervals.

### The two-viruses-one-cell model

A sub-genomic viral vector, p1004.RN3, was generated by deleting the Tat gene from the p932.RN3 backbone. The p1004.RN3 vector co-expresses d2mScarlet and Gaussia luciferase from an HIV-1C LTR, where luciferase expression is under the control of an IRES element. The half-life of the reporter protein mScarlet is approximately 2 h. The p1004.RN3 vector was constructed in three successive steps, as described below. First, the destabilization domain was amplified from p932.RN3 using the primer pair N4111 and N4112, and the fragment was inserted between NotI and EcoRI of a vector pCL-TdTomato-IGLuc-RRE plasmid ^93^, thus replacing the TdTomato gene from the vector. During this manipulation, simultaneously, recognition sites for two unique RE, AgeI and BlpI, were introduced upstream of the destabilizing domain through primer N4111. In the second step, the ORF of mScarlet was amplified from vector p1011 using the primer pair N4113 and N4114 and placed directionally between the two unique RE AgeI and BlpI upstream of the destabilizing domain, creating the vector pCL-d2mScarlet-IGluc-RRE. In the third step, the d2mScarlet-IRES-GLuc cassette was amplified using the primer pair N4243 and N4244 and placed between the BamHI and XhoI of the p932.RN3 vector replacing the original d2EGFP-IRES-Tat cassette, thus generating the final vector p1004.RN3. Subsequently, the 3’-LTR of p1004.RN3 vector was substituted with the R2N3-LTR, using AscI and SacII RE sites, to generate p1004.R2N3 vector.

We constructed Jurkat T-cell pools harboring combinations of two different proviral strains, each strain encoding a different fluorescent protein, and both viruses containing the same or different LTRs (RN3 and R2N3). Of note, Tat was encoded from only one of the two LTRs in the cells. The bi-reporter Jurkat T-cell model was generated in two sequential steps. The cells were first infected with the Tat-encoding viral strain and then sorted using the co-expressed fluorescent protein. The single-reporter cells were subsequently infected with the second viral strain expressing the second fluorescent protein, and the cells co-expressing both the reporter proteins were sorted and expanded (Figure 3B).

For example, Jurkat T-cells were transduced first with the virus co-expressing d2EGFP and Tat at an expected infectivity level of ≤10% (RIU ∼0.05-0.1). Six hours post-infection, the medium was replenished with the RPMI 1640 medium supplemented with 10% FBS, 2 mM Glutamine, 100 units/ml Penicillin G, and 100 μg/ml Streptomycin. After 48 h of the infection, the cells were activated with a cocktail of activators containing TNF-α (10 ng/ml), PMA (5 ng/ml), and HMBA (5 mM). After 24 h of activation, GFP^+ve^ cells were sorted (BD FACSAriaIII sorter, BD Biosciences, New Jersey, USA). Cells were allowed to expand for 8-10 days before the second transduction. Subsequently, the GFP^+ve^ cells were transduced with the second viral strain that co-expressed mScarlet and Gaussia luciferase. The procedures, infection parameters, and activation conditions were similar to the first round of infection described above. The double-positive cells (GFP and RFP positive cells) were sorted using a FACS machine. The cells were cultured for 15-20 days to relax the expression of both fluorescent proteins. Once a large number of cells (approximately >85%) downregulated GFP and RFP expression, latency reversal profiles were analyzed.

### RT-PCR to determine Tat transcripts

Total RNA was extracted from 1×10^6^ infected Jurkat T-cells using TRIzol (Cat. No., Sigma-Aldrich, USA). Briefly, 2 μg of RNA was treated with DNase I (Cat. No. 18068015, New England Biolabs, USA) at 37°C for 5 min, and the DNase I was inactivated by incubating the sample at 65°C for 10 min in the presence of 2.5 mM EDTA. The complementary DNA (cDNA) was synthesized from 1 μg of RNA using the pool of HIV-1-specific and Oligo dT primers N4101, N4102, and N4103 (Supplementary file 2) and a commercial kit (OneScript cDNA synthesis kit, Cat. No. G234, Applied Biological Materials Inc., Canada). The reaction vials containing RNA and primer mix were incubated for 5 min at 65°C and 2 min on ice. Next, the enzyme mix was added, and the vials were incubated for 15 min at 50°C. The reaction was terminated by incubating the samples at 85°C for 5 min. The cDNA was used for the qPCR analysis of Tat transcript and GAPDH using the primer pairs N4262-N4263 and N4415-N4416, respectively (Supplementary file 2).

### Replication kinetics

PBMCs were isolated from 10 ml of blood of healthy donors by Ficoll density gradient centrifugation. The PBMC were CD8 cell-depleted using a commercial kit (Cat. No. 15663, Stemcell Technologies, Canada) and activated by TCR engagement as described previously before infection with viral strains. Viral titers of the stocks were determined in the TZM-bl cells by quantifying luciferase activity. A viral stock equivalent to luciferase activity of 150,000 RLU was used for the infection. PBMC were incubated with full-length promoter-variant viral strains (Figure S2) for six hours in the RPMI 1640 medium supplemented with 10 μg/ml of DEAE dextran, 10% FBS, 2 mM Glutamine, 100 units/ml Penicillin G, and 100 μg/ml Streptomycin. Six hours post-infection, the cells were washed twice with PBS and incubated with RPMI 1640 medium supplemented with 10% FBS, 2 mM Glutamine, 100 units/ml Penicillin G, and 100 g/ml Streptomycin. The cells were monitored, and the medium was replenished every three days. The cultures were fed every 15 days with autologous and activated PBMC. The secretion of p24 into the culture medium was monitored by ELISA using a commercial kit (Cat. No. IR232096, J. Mitra & Co. Pvt. Ltd., India). The viral growth curve for each viral strain was constructed to determine the replication profile.

### Flow cytometry and statistical data analysis

Flow cytometry data were analyzed using FCS Express 6 software (De Novo Software, Los Angeles, CA). All data in figures 1-3 and S3-5, S8, and S10 were plotted using GraphPad Prism software (versions 9 and 10). Statistical evaluation for all the experiments was performed using the same software. The statistical tests performed for each experiment are depicted in the corresponding figure legends along with the P values.

### Next-generation sequencing: HIV-1 promoter analysis from activated HIV-1^+ve^ primary CD4 T-cells

Activation of the primary CD4 T-cells, isolation of the gDNA and cellular RNA, and RT-PCR: Stored PBMC of all study subjects were thawed at 37°C, and the CD4 T-cells were enriched by negative selection using magnetic beads (Cat. No. 19052, Stemcell Technologies, Canada). Next, the CD4 T-cells were divided into fractions for three experimental conditions as follows: No activation, PMA (40 ng/ml)+ Ionomycin (1 μM), and Prostratin (10 μM). Approximately 0.1-0.3×10^6^ CD4 cells/assay were activated for 24 h, except for the TCR-mediated activation (Dynabeads Human T-Activator CD3/CD28, Cat. No. 11131D, Thermo Fisher Scientific, USA), where the cells were activated for 72 h. After the activation, both the gDNA and cellular RNA fractions were purified from the cells using a commercial kit (Allprep DNA/RNA Micro Kit, Cat. No. 80284, Qiagen, Germany). cDNA was synthesized from the cellular RNA, as described above. The gDNA and cDNA preparations were used to amplify the HIV-1 LTR.

Amplification of the LTR region from gDNA and cellular RNA, library preparation, and sequencing: The amplification of the U3 region (∼247-272 bp) from gDNA and cDNA was performed by PCR using the primers with a unique sequence barcode at the 5’-end as presented in the supplementary file 2, Patient samples-A and -B. Each sample was amplified using primers containing a unique eight-bp barcode sequence specific for each sample (Supplementary file 2, Patient samples-C). The concentration of the purified PCR product was determined using the QubitTM dsDNA BR assay kit (Cat. No. Q32850, Invitrogen, California, United States). All the samples were pooled in equal amounts (10 ng) and were processed together. All the sample pools sequenced were pulsed with a defined quantity (10 ng) of the LTR amplicon of Indie.C1, a reference HIV-1C molecular clone, as an internal control for the sequencing quality.

The PCR products were sequenced using the NovaSeq platform. The DNA was quantified using the QUBIT 3 Fluorometer and a dsDNA HS Dye. Library preparation was performed using the Qiagen FX DNA library preparation kit. (Cat.No.180175, Qiagen, Germany). The adapters used in this protocol include dual index barcodes required for the multiplexing. Since the index barcodes were provided with platform-specific adapters, the PCR Enrichment and indexing PCR steps were excluded during the library preparation. The library was sequenced using the NovaSeq system following the manufacturer’s instructions. X A custom Python script was developed to analyze the data and count the NF-κB and RBEIII sites. The script has been deposited athttps://github.com/hivaidslab.). The same script was used to analyze intact genomes reported by Garcia-Broncano et. al, 2019; Lee et. al, 2019.

### The Tat/Rev Induced Limiting Dilution Assay (TILDA)

TILDA was performed as described previously ^94^. Briefly, CD4+ cells were enriched from stored PBMCs using a negative magnetic selection kit (EasySep Human CD4+ T cell Isolation Kit, #19052, Stemcell Technologies Inc., Canada). The enrichment was confirmed via flow cytometry using PE Mouse Anti-Human CD4+ antibody (#561844) and PE Mouse IgG1, k Isotype Control (#555749, both from BD Biosciences, USA). The cells were resuspended at 2 × 10⁶ cells/ml in RPMI 1640 medium with 10% fetal bovine serum, 100 U/ml penicillin G, 2 mM L-glutamine, and 100 μg/ml streptomycin, and rested at 37°C for 3–5 h. Activation was performed using 100 ng/ml PMA (#P1585, Sigma-Aldrich, USA) and 1 μg/ml Ionomycin (#I0634, Sigma-Aldrich, USA) for 12 h. Activated cells were counted and diluted to 27 × 10⁶, 9 × 10⁶, and 3 × 10⁶ cells/ml. One mlof each dilution, corresponding to 27,000, 9,000, and 3,000 cells, was added to 9 replicate wells containing 5 μL 2X reaction buffer, 0.1 μL RNase inhibitor, 0.2 μL of 25 μM primers (N2830 and N2831, reported previously,^94^, 0.5 μL of 2 μM HIV-specific cDNA primer pool, 2.8 μL H₂O, and 0.2 μL SuperScript III Platinum Taq (SuperScript III Platinum One-Step qRT-PCR Kit, #11732020, Invitrogen, USA). Amplification conditions were 50°C for 15 min, 95°C for 2 min, and 25 cycles (95°C for 15 s, 60°C for 4 min). PCR products were diluted with 40 μL TE, and 1 μL was used as a template for a second round of PCR with MyTaq™ DNA polymerase (#BIO-21105, Bioline, UK). This reaction included 2 μL 5X MyTaq Buffer, 2 μL each primer (N2832, N2833; 10 μM), 0.25 μL HIV probe (5 μM), and 3.4 μL H₂O in a 10 μL volume. The program included 95°C for 10 min, followed by 40 cycles (95°C for 10 s, 60°C for 30 s, 72°C for 30 s). Positive wells were counted, and the frequency of cells with inducible HIV msRNA was calculated using the maximum likelihood method. A subset of TILDA-positive wells was selected, and 1 μl of the first-round PCR product was subjected to the LTR-PCR described above to identify variants that are activated upon stimulation.

### Integration Site Analysis

One μg of genomic DNA was sonicated using the Bioruptor® Plus sonicator for 16 seconds over three cycles. The sonicated DNA was then subjected to end-repair and A-tailing, followed by adapter ligation using the NEBNext® Ultra™ II DNA Library Prep Kit (New England Biolabs, Cat. No. E7645). Adapter sequences were based on the specifications provided in the Lenti-X™ Integration Site Analysis Kit (Takara Bio, Cat. No. 631263). Following ligation, 200 ng of the adapter-ligated library was used as input for two separate nested PCR reactions targeting the 5′ and 3′ LTR regions. For the 5′ LTR, the HIV-specific primers were 5′-GTACAGGCGAAAAGCAGCTGCTTATATGC-3′ (outer round) and 5′-AGCGAAAAGCAGCTGCTTATATGCAGCATCTGAGGG-3′ (inner round). For the 3′ LTR, primers used were 5′-CCACTGACCTTTGGATGGTGCTTCAAG-3′ (outer round) and 5′-TGTGCCAGCATGGAATGGARGATGA-3′ (inner round). Adapter primer sequences were based on, but slightly modified from, the Lenti-X™ Integration Site Analysis Kit (Takara Bio, Cat. No. 631263). The adapter primers used were 5′-GTAATACGACTCACTATAGGGC-3′ for the outer round and 5′-ACTATAGGGCACGCGTGGT-3′ for the inner round. Two independent reactions were performed in duplicate for each subject. The PCR products were purified and subjected to Nanopore sequencing, and the resulting data were analyzed using a custom Python script.

### Data analysis

Data were analyzed using a custom pipeline as described below. First, the quality of the reads was assessed using FastQC (version 0.11.5). The sequencing adapters were removed using cutadapt (version 3.1) from the raw paired-end data. Second, the paired reads from both the forward and reverse files were merged using VSEARCH (version 2.17.1) with the minimum merge length set to 200 and an overlap of at least 10 base pairs. Around 95% of the reads were merged. Third, the merged reads were mapped to the LTR region from reference sequences of Indie.C1 (AB023804.1) and D24 (EF469243.2), using local alignment with Bowtie2 software. As a result, 99.8% of the merged reads were mapped to the references. Fourth, the mapped reads were demultiplexed using custom Python scripts into individual samples based on the combination of forward and reverse barcodes.

Further, the reads from each sample were sorted based on the number of RBEIII and NF-κB motifs using a Python script. Upon sorting in respective groups, the sequences were processed using a Python script to identify unique sequences and to provide the representative sequence from each group of every sample. Finally, the percentage prevalence of double- and single-RBEIII variants was calculated.

## Abbreviations

TFBS: Transcription factor binding site
HIV-1C: HIV-1 subtype C
LTR: Long Terminal Repeats
C-LTR: HIV-1C LTR
NGS: Next-generation sequencing
RT-PCR: Reverse transcription polymerase chain reaction
MTRC: Master transcription regulatory circuit
TFC: Transcription factor complex
PBMC: Peripheral Blood Mononuclear Cell
TILDA: Tat/Rev Induced Limiting Dilution Assay
gDNA: genomic DNA
cDNA: complementary DNA
pRNA: Plasma RNA
LFU: Lost-to-Follow-Up
PTC: Post-Treatment Control
ART: Anti- Retroviral Therapy

## Competing Interest Statement

The authors declare no competing interests.

## Author Contributions

D.B. performed research, analyzed data, and wrote the paper. A.P. designed research, analysed data, and co-wrote the paper. S.S., S.M., D.P., C.S., H.B., J.S., R.M., M.S., S.S., performed research and analyzed data. M.S. and J.S. performed the bioinformatics analysis. N.N. assisted in flow cytometry experiments and analysis. S.P.M, M.D, A.K.S, and K.G.M designed experiments and were involved in the establishment and procurement of clinical samples. S.N.B. and T.K.K contributed to the study design, review of work progress, and manuscript editing. U.R. designed the research and wrote the paper.

## Funding

This work was supported by funds from the Department of Biotechnology, Ministry of Science and Technology, Government of India (Sanction orders no. BT/PR7359/MED/29/651/2012 and BT/NETHERLANDS/RG/40/2015), Corporate Social Responsibility funds from Gennova Biopharmaceuticals Ltd., Maharashtra, India to UR. Funds of the Science and Engineering Research Board (Sanction order no. SPR/2021/000338-G) to TKK are acknowledged. Financial support to YRGCARE from the Department of Biotechnology, Ministry of Science and Technology, Government of India (Sanction order no. BT/NETHERLANDS/RG/40/2015) and the United States Agency for International Development (USAID, Cooperative Agreement No. AID-OAA-A-16-00032) to YRG CARE, have been used.

## Acknowledgments

We thank Dr. Robert Siliciano for kindly providing the plasmids used to improve long-term CD4 cell culture, which forms an important model for our study. We thank Dr Kushgra Bansal for the discussions during the NGS data analysis. We thank the study participants for their participation.

---

**Supplementary Figure 1:**
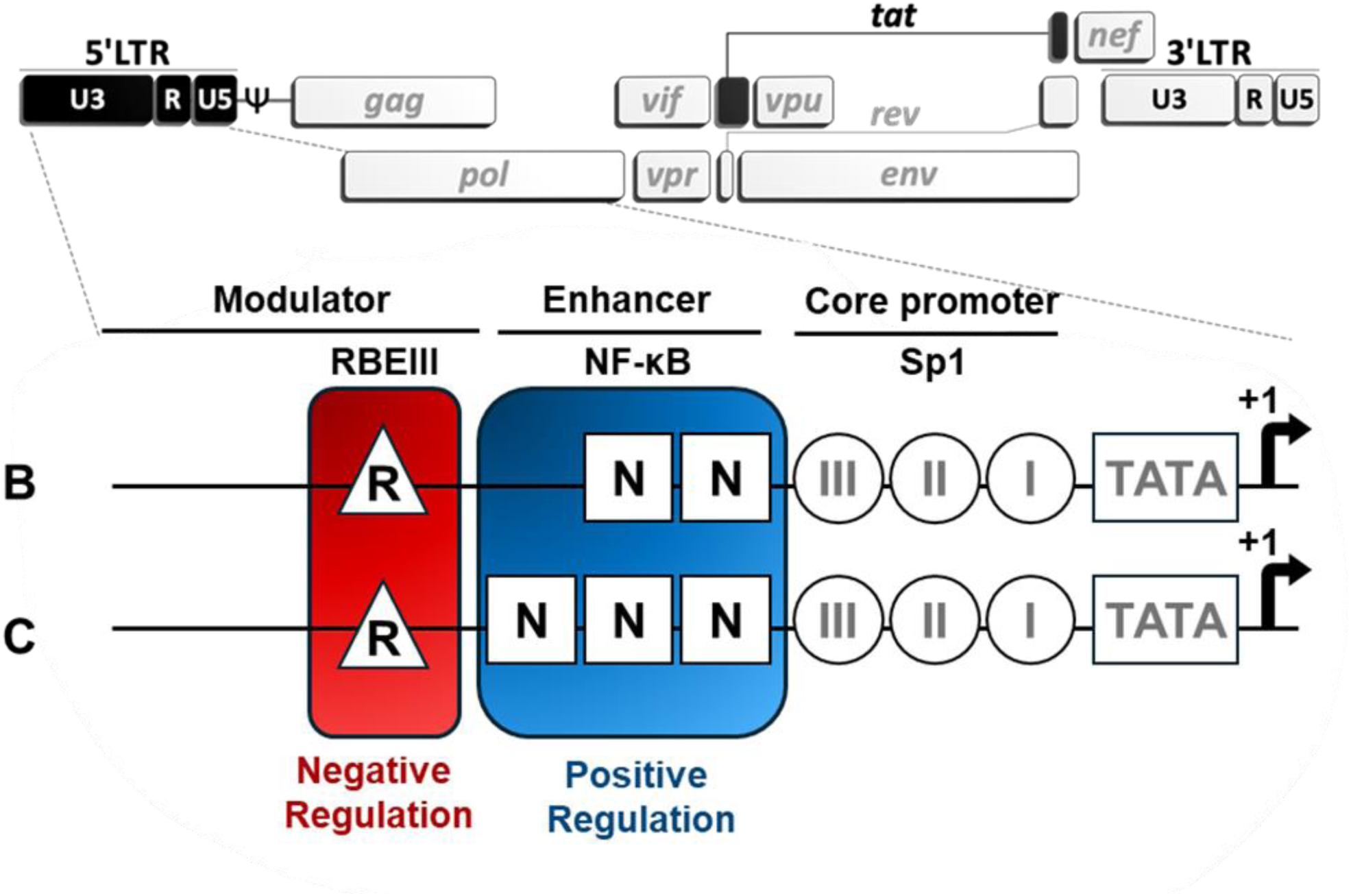
The TFBS profile of HIV-1 LTR. The genome organization of HIV-1 is depicted in the top panel. The U3 region of the LTR is enlarged and the arrangement of a few key TFBS in the modulatory, enhancer, and core promoter regions is portrayed. Note that HIV-1C LTR contains three NF-κB motifs, whereas that of HIV-1B and several other HIV-1 subtypes contains only two. The three Sp1 motifs (circles), the TATA box (rectangle), and the transcription start site (black arrow) are depicted downstream of the enhancer. Two key sequence clusters centered on the RBEIII (R, triangle) and NF-κB (N, square box) motifs are highlighted. Based on the data presented here, we propose that the NF-κB and RBEIII clusters are predominantly transcription-suppressing and transcription-enhancing under the conditions of absent (red) or present (blue) cellular activation, respectively.

**Supplementary Figure 2:**
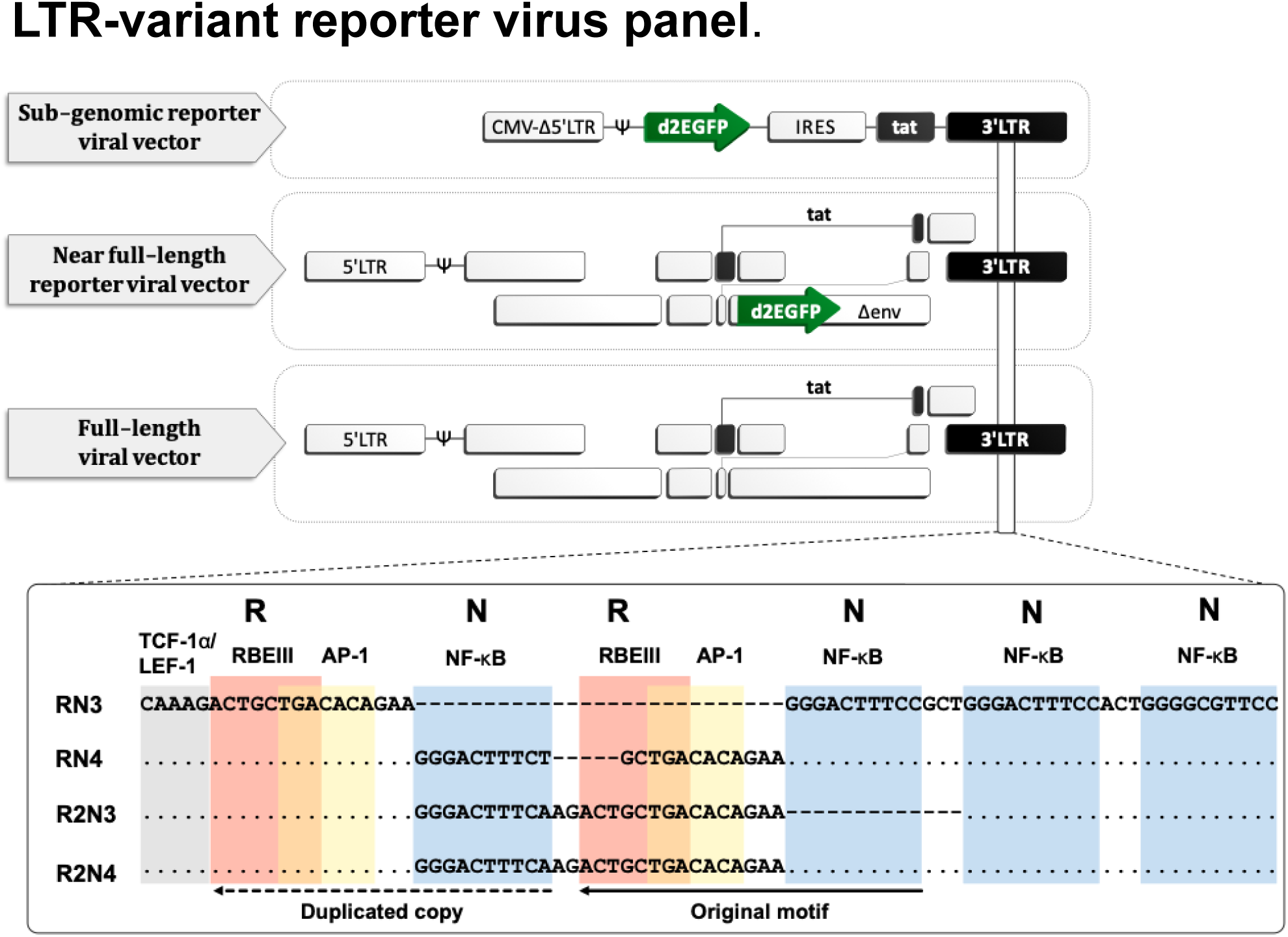
Construction of a panel of LTR-variant viral strains. Analogous panels of four variant LTRs (RN3, RN4, R2N3, and R2N4) were constructed using three different viral vector backbones. The sub-genomic viral strains co-express d2EGFP and Tat. The near full-length viral strains encode d2EGFP and are devoid of the envelope. The full-length viral strains produce replication-competent virus particles. The multi-sequence alignment of the promoter regions (the bottom panel) depicts the TFBS variations engineered into the 3’-LTR of the viral vectors. In the sequence alignment, dots and dashes represent sequence identity and deletion, respectively. Different TFBS are highlighted using different shades - the RBEIII motifs (R, red), NF-κB motifs (N, blue), TCF-1α/ E-1 sites (grey), and the AP-1 elements (yellow). The three base pairs that overlap between the RBEIII and AP-1 sites are shown in orange. The original and duplicated sequence motifs are represented by the solid and dotted arrows, respectively. The TFBS motifs used in vector engineering are representative of the sequences of primary viral strains.

**Supplementary Figure 3:**
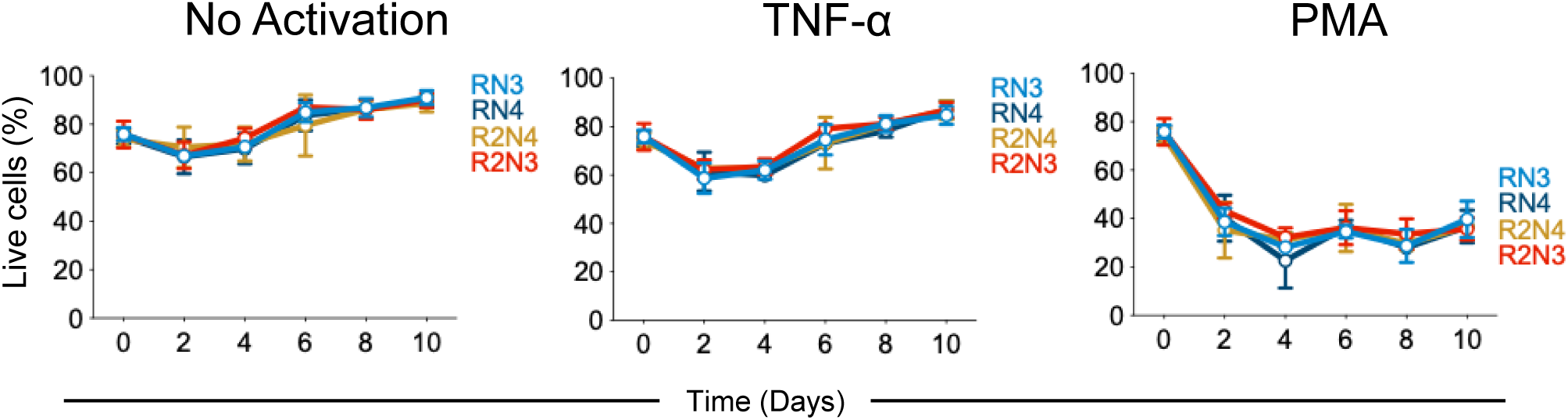
The percentages of live cells in Fig. 1E across the various activation conditions

**Supplementary Figure 4.**
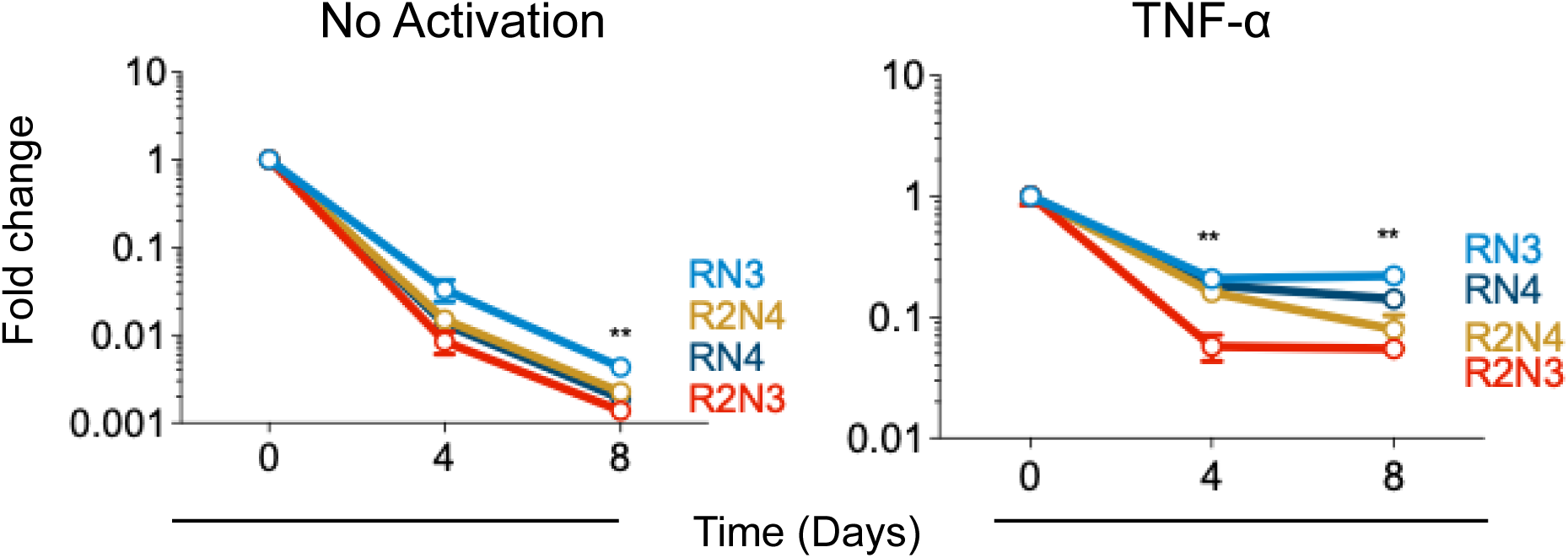
Tat transcript levels of the reporter virus panel shown in Fig. 1B. Total cellular RNA was isolated, and reverse transcribed, and Tat and GAPDH transcript levels were estimated at the 0, 4, and 8-day time points using qPCR. The data shown are Tat transcript levels normalized with GAPDH expression. The statistical analysis represents a comparison between RN3 and R2N3 viral strains, while comparisons with the remaining viral strains are not marked on the graphs. p=0.033 (*), p=0.002 (**), p<0.001 (***), Two-way ANOVA with Dunnett multiple comparisons test.

**Supplementary Figure 5:**
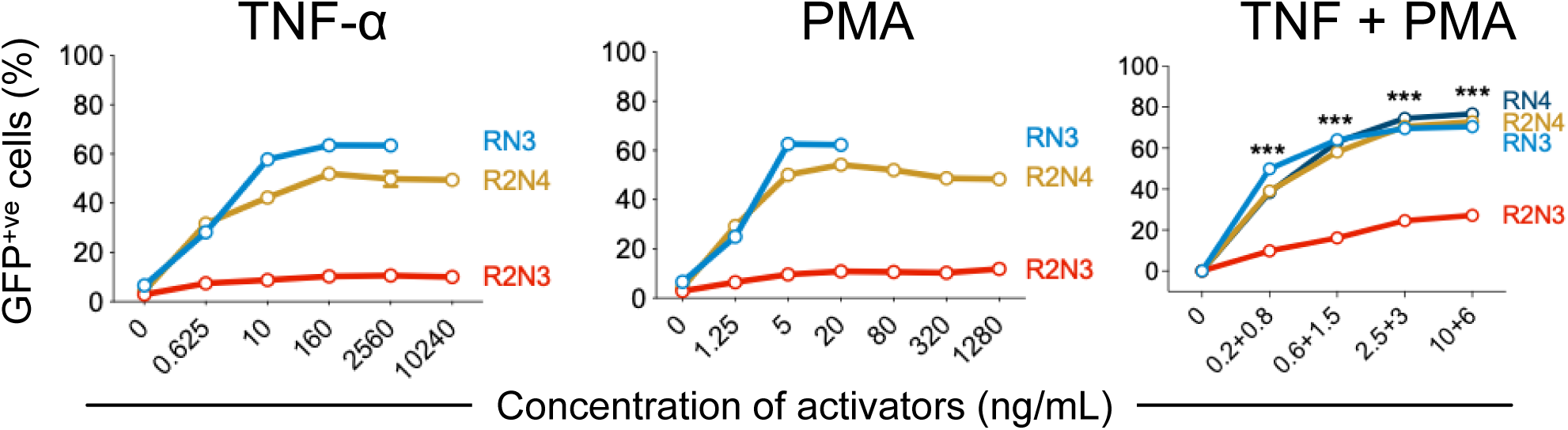
Latency Reversal profiles of LTR-variant sub-genomic reporter viral strains: The latency reversal assays were performed in the presence of increasing concentrations of TNF-α and/or PMA. The data are representative of two independent experiments. The statistical analysis represents a comparison between RN3 and R2N3 viral strains while comparisons with remaining viral strains are not marked on graphs. p=0.033(*), p=0.002(**), p<0.001 (***), Two-way ANOVA with Dunnett multiple comparisons test.

**Supplementary Figure 6:**
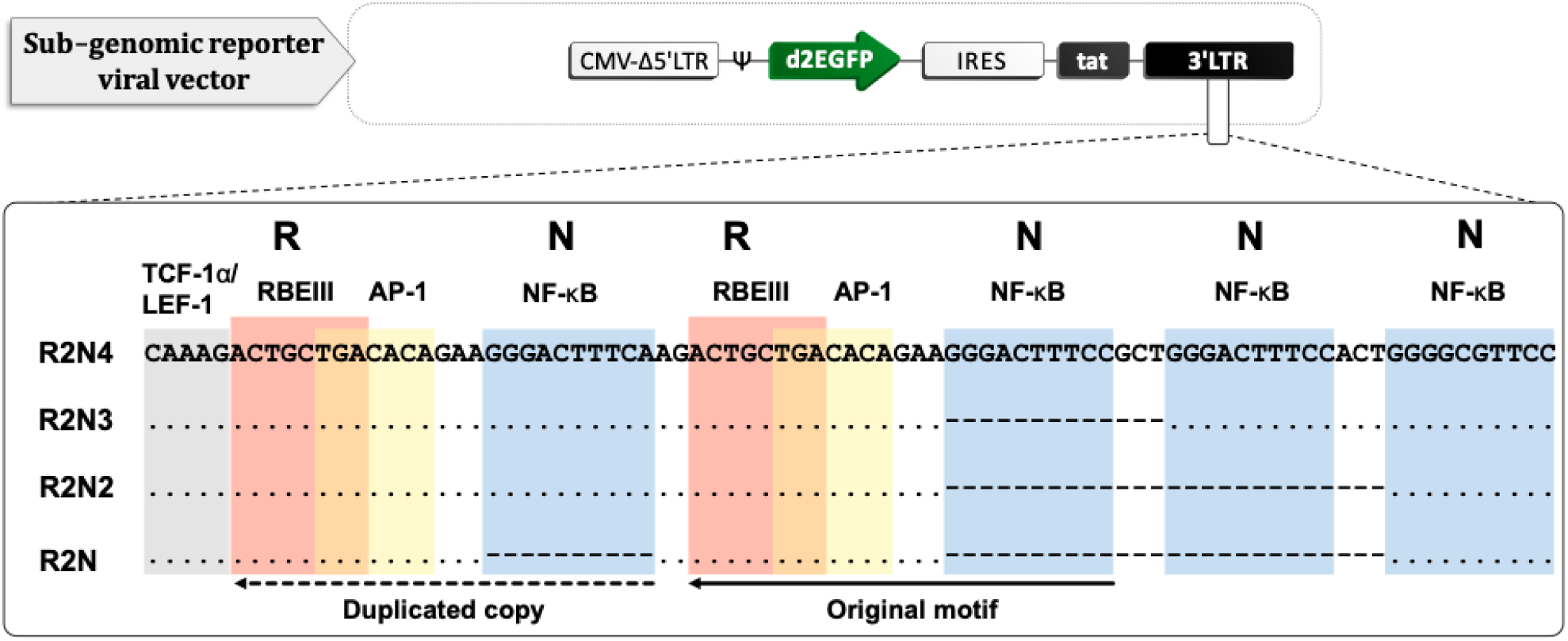
A panel of sub-genomic reporter R2 viral strains containing varying numbers of the NF-κB motif. The panel comprises four variant LTRs with progressively decreasing copies of the NF-κB motif, ranging from 4 to 1 (R2N4, R2N3, R2N2, and R2N, respectively). The multi-sequence alignment of the promoter regions (the bottom panel) depicts the TFBS variations engineered into the 3’-LTR of the viral vectors. In the sequence alignment, dots and dashes represent sequence identity and deletion, respectively. Different TFBS are highlighted using different shades - the RBEIII motifs (R, red), NF-κB motifs (N, blue), TCF-1α/ E-1 sites (grey), and the AP-1 elements (yellow). The three base pairs that overlap between the RBEIII and AP-1 sites are shown in orange. The original and duplicated sequence motifs are represented by the solid and dotted arrows, respectively. The TFBS motifs used in vector engineering are representative of the sequences of primary viral strains.

**Supplementary Figure 7:**
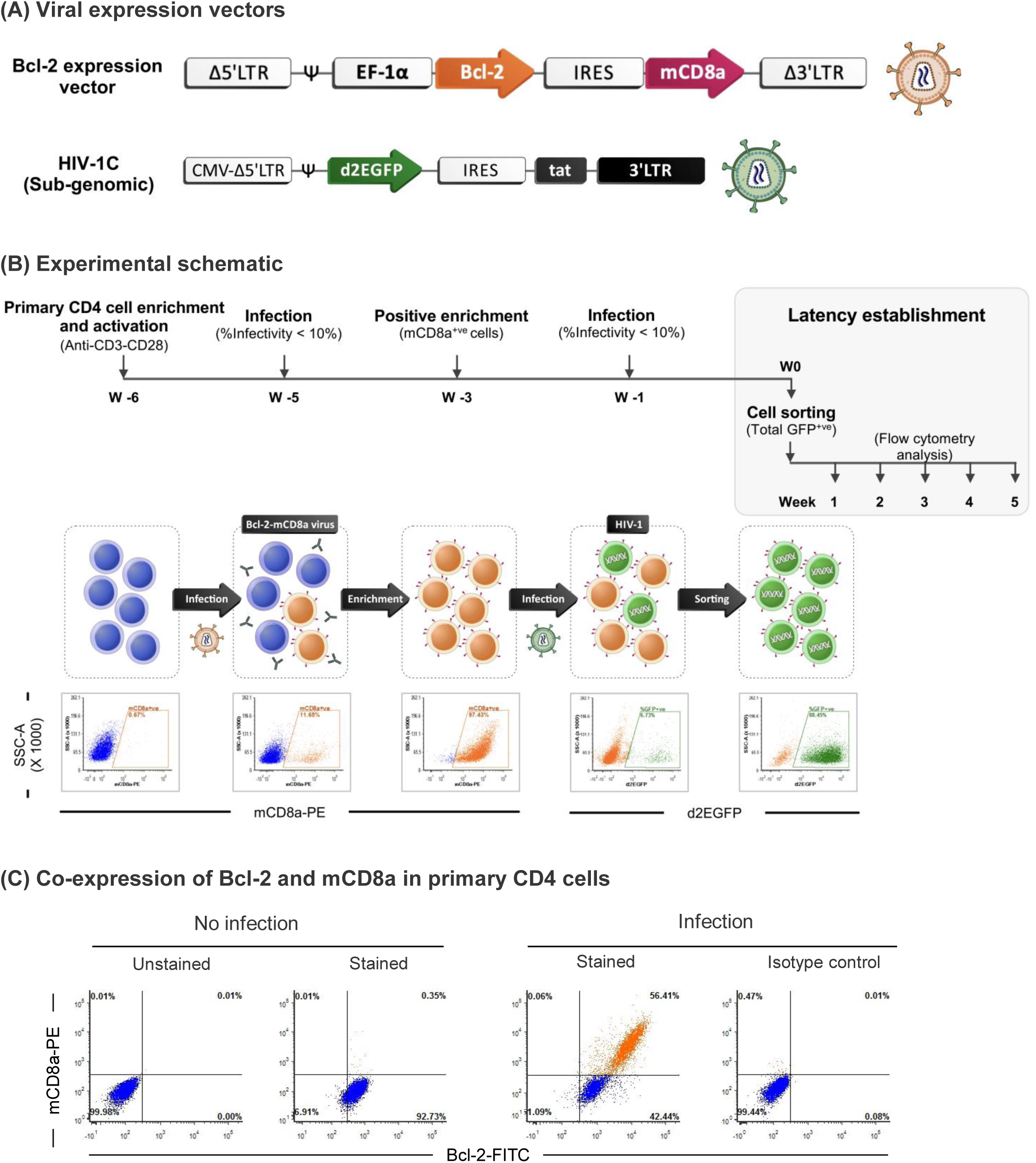
Long-term primary CD4 culture for HIV-1 latency establishment. (A) The schematic representation of the viral vectors. The lentiviral vector, EB-FLV-mCD8, co-expresses Bcl-2 and mouse CD8a.The sub-genomic HIV-1 reporter vector panel was as in Supplementary Figure 2. (A) The experimental schematic and representative flow cytometry plots. A schematic of the workflow, a timeline of the infection, selection of the infected cells, and latency establishment evaluation are depicted. Primary CD4 cells (blue) were first infected with the Bcl-2-expressing lentivirus (orange), infected cells were selected, and a pool of the cells was re-infected with the HIV-1 vector (green). Cells expressing Bcl-2 stably were enriched with anti-mCD8a-coated magnetic beads (orange, plot-3). HIV-1 vector-infected cells were selected using flow cytometry. The sorted GFP+ve cells (green, plot-5) were used in the latency experiments. The representative flow profiles presented here are derived from the infection of the RN3 viral strain, and the profiles of all the other promoter variants are comparable. (C) Co-expression of Bcl-2 and mCD8a in primary CD4 cells 48 h post-infection of EB-FLV-mCD8 lentivirus. Cells were stained using anti-Bcl-2 and anti-mCD8a antibodies and analyzed using flow cytometry. The plots with only Bcl-2 (top panel), only mCD8a (middle panel) or both in the same plot (bottom panel) are depicted. Isotype controls are shown.

**Supplementary Figure 8:**
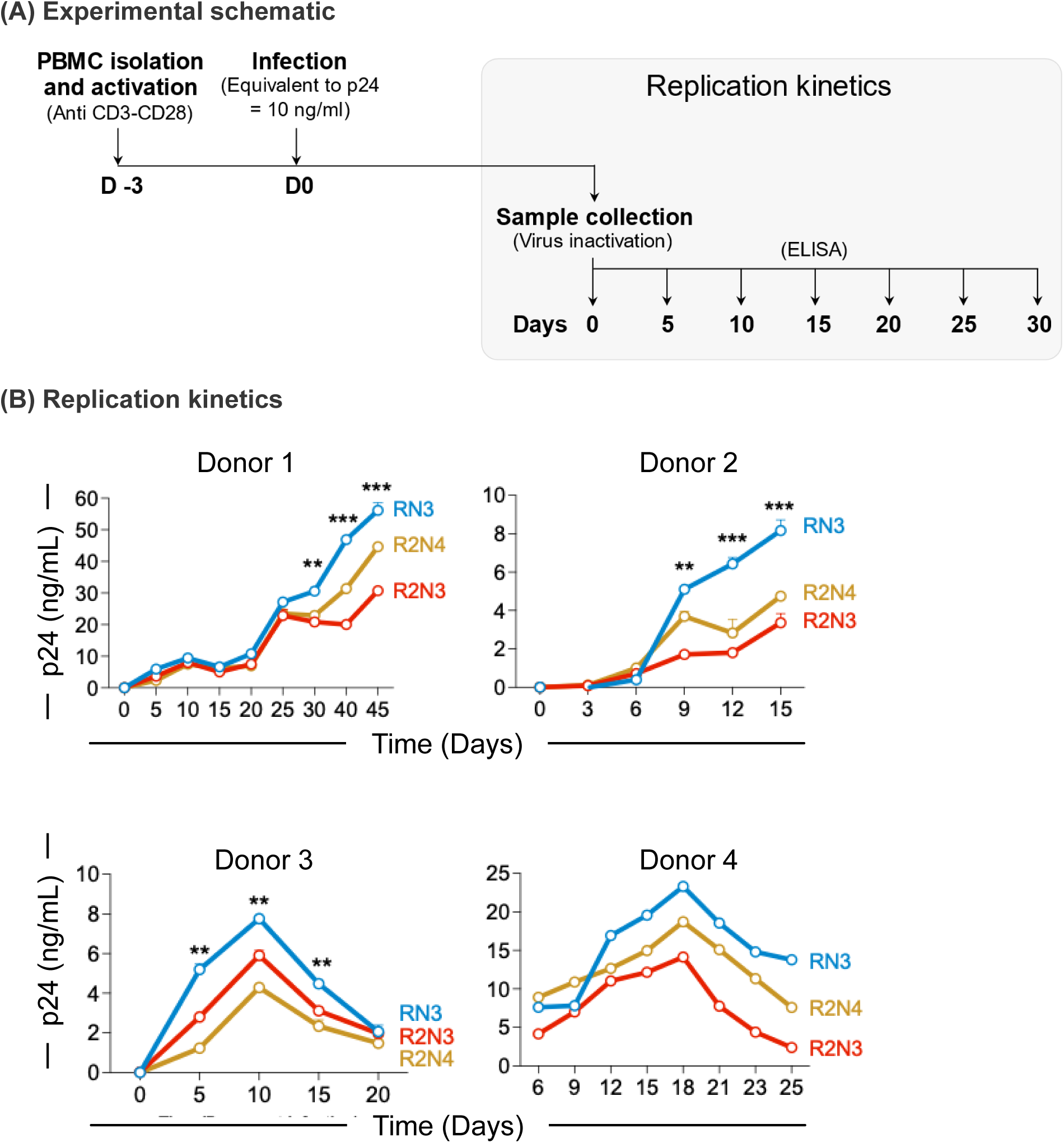
Replication kinetics of promoter-variant full-length viral strains in PBMCs of healthy donors. (A) A schematic of the workflow and timeline of the infection, sample collection, and p24 quantification by ELISA are depicted. (B) The replication kinetics of RN3, R2N3, and R2N4 in the PBMCs of four healthy donors. The statistical analysis represents a comparison between RN3 and R2N3 viral strains. p=0.033(*), p=0.002(**), p<0.001 (***), Two-way ANOVA with Dunnett multiple comparisons test.

**Supplementary figure 9:**
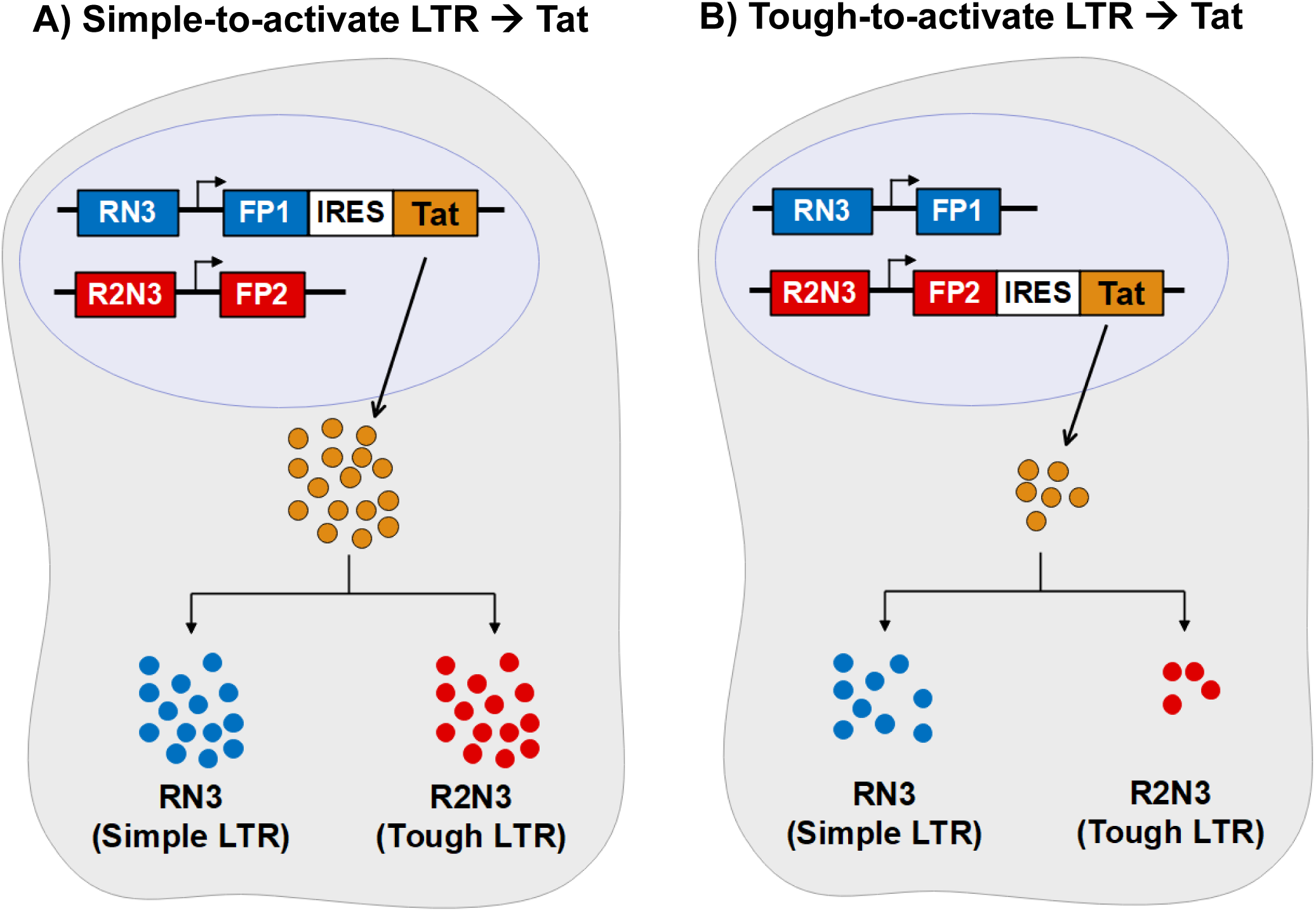
A schematic representation of the MTRC governing the ON/OFF decisions. The schematic portrays how transcriptional strength differences of the LTR can alter the ON/OFF decisions when all experimental parameters are normalized. Two different scenarios are projected when Tat is encoded by, **(A)** the simple-to-activate RN3-LTR or **(B)** the tough-to-activate R2N3-LTR. Cellular activation of a low or moderate intensity can induce significantly high levels of Tat expression (blue dots) from RN3-LTR, but not the R2N3-LTR. High-level intracellular Tat concentration (orange dots) driven by the RN3-LTR can induce reporter gene expression from both the LTRs efficiently. In contrast, sub-optimal Tat drives only a partial activation of the RN3-LTR and very low activation of the R2N3-LTR. The ON/OFF decisions are consistent and highly reproducible in the cell pool, thus normalizing integration site differences. The schematic is based on data presented in Figure 3. FP=Fluorescent protein.

**Supplementary figure 10:**
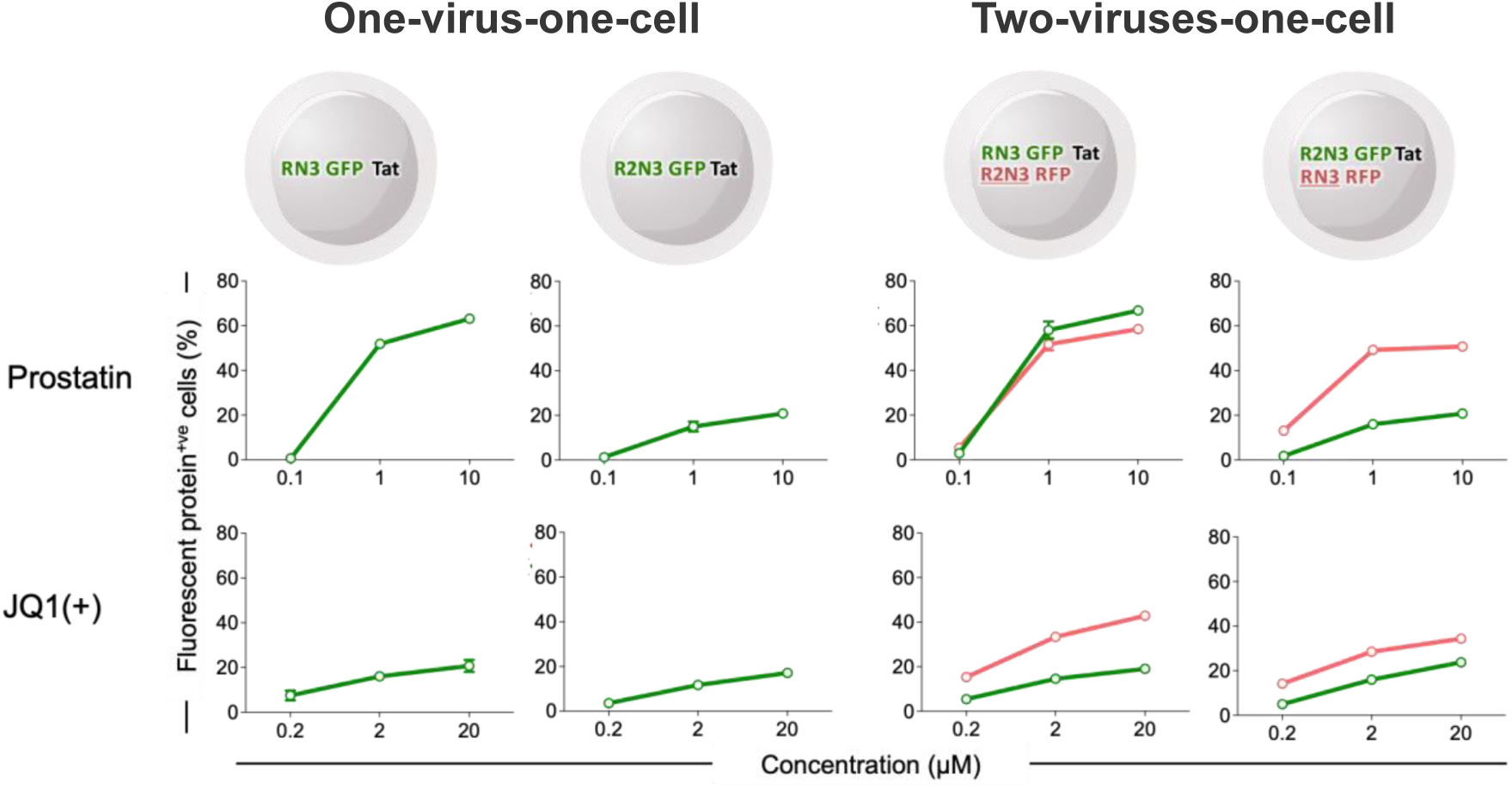
Latency reversal kinetics of RN3- and R2N3-LTRs using two LRAs. The two left panels represent the mono-infection models harbouring RN3 or R2N3 viral strains, co-expressing d2EGFP and Tat. The right two panels represent the two-viruses-one-cell models, where Tat is encoded by one of the two LTRs. The activation status of each LTR could be detected by monitoring the encoded fluorescent protein, d2EGFP or d2mScarlet. The latency reversal assays were performed in the presence of increasing concentrations of LRAs, Prostratin, or JQ1(+). The red and green curves represent the expression profiles of d2EGFP and d2mScarlet, respectively.

**Supplementary figure 11:**
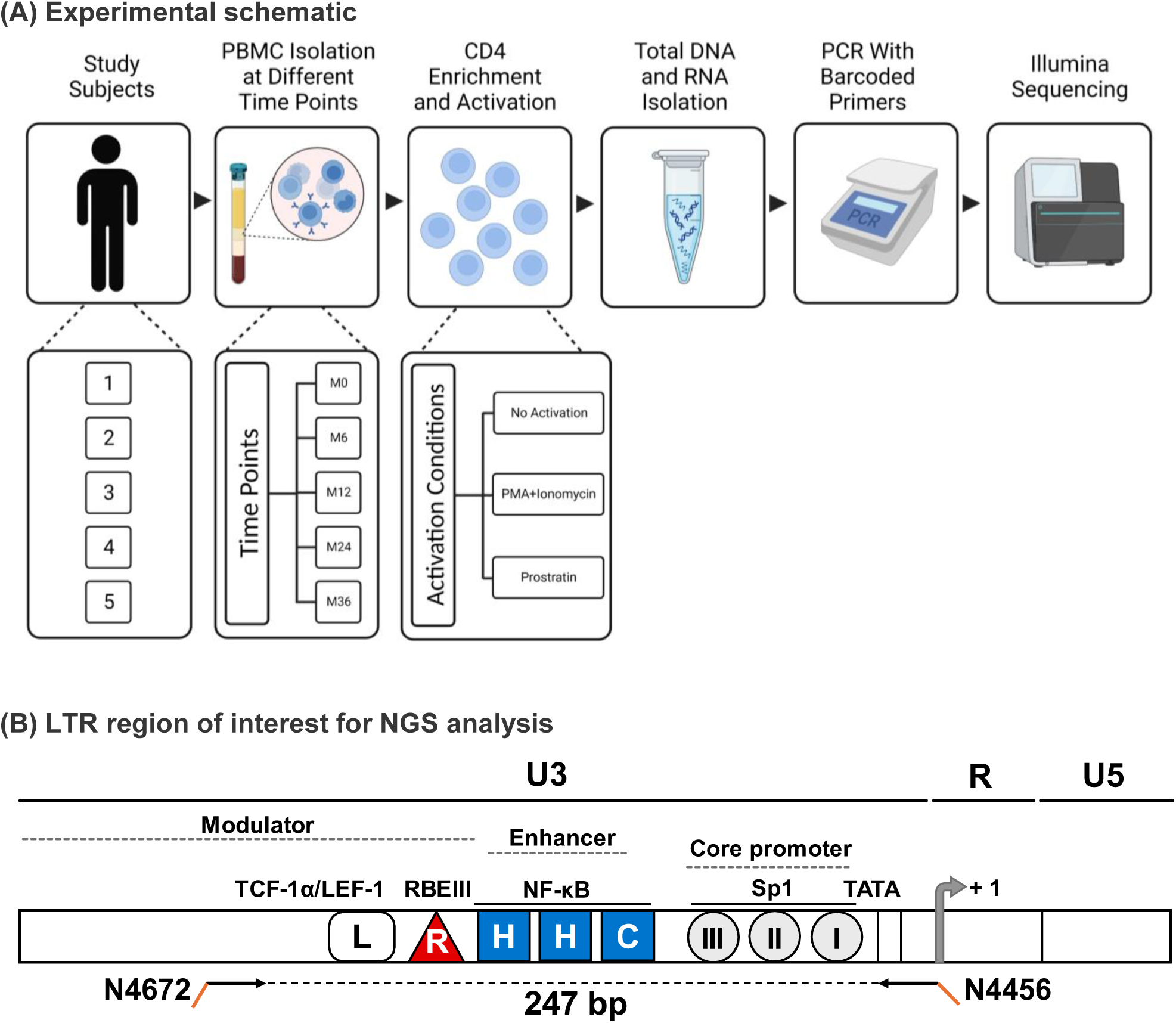
The experimental schematic of evaluating viral transcripts by NGS. **(A)** The workflow from sample collection to sequencing is depicted. CD4 T-cells were enriched from stored PBMCs collected at several follow-up points, subjected to diverse activation conditions, and genomic DNA or cellular RNA was extracted. Finally, the specific region of LTR was amplified and subjected to Illumina sequencing. Note that two biological replicate samples were used in parallel from all the samples under each experimental condition. **(B)** The PCR amplification schematic. A 247 bp region of the LTR comprising the regulatory elements of interest was amplified using the primer pair N4672-N4456 (the black arrows) depicted. Important TFBS are shown, with the RBEIII and NF-κB motifs, marked as red triangle and blue squares, respectively. The orange tags of the primers represent the 8 bp barcodes.

**Supplementary Figure 12:**
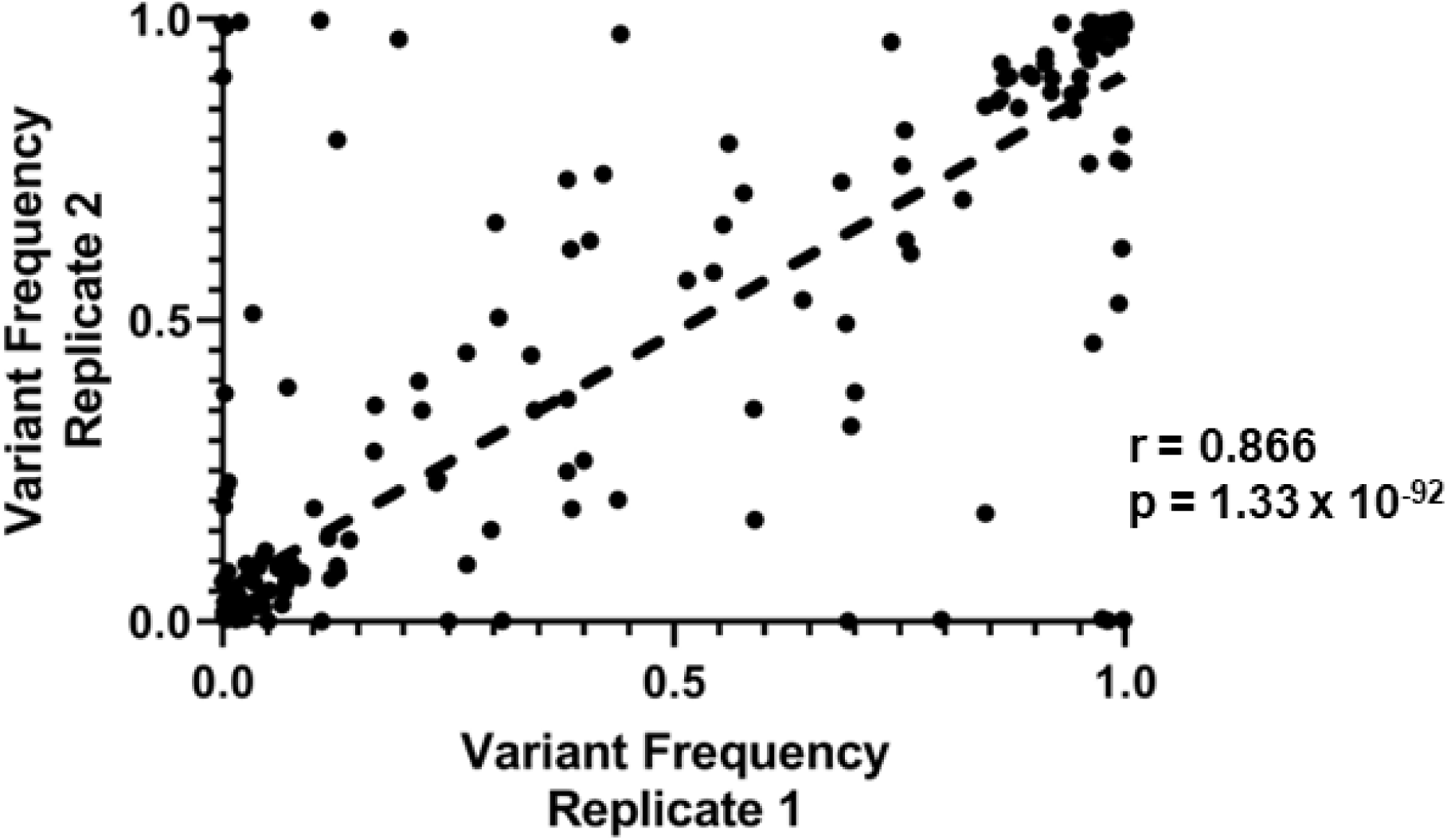
Scatter plot depicting variant frequency at all time points and all conditions assayed for the three major variants RN3, R2N3 and R2N4 for the NGS analysis. Each dot represents the frequency of a given variant in both replicates. The correlation coefficient and its associated p value are as shown.

**Supplementary Figure 13:**
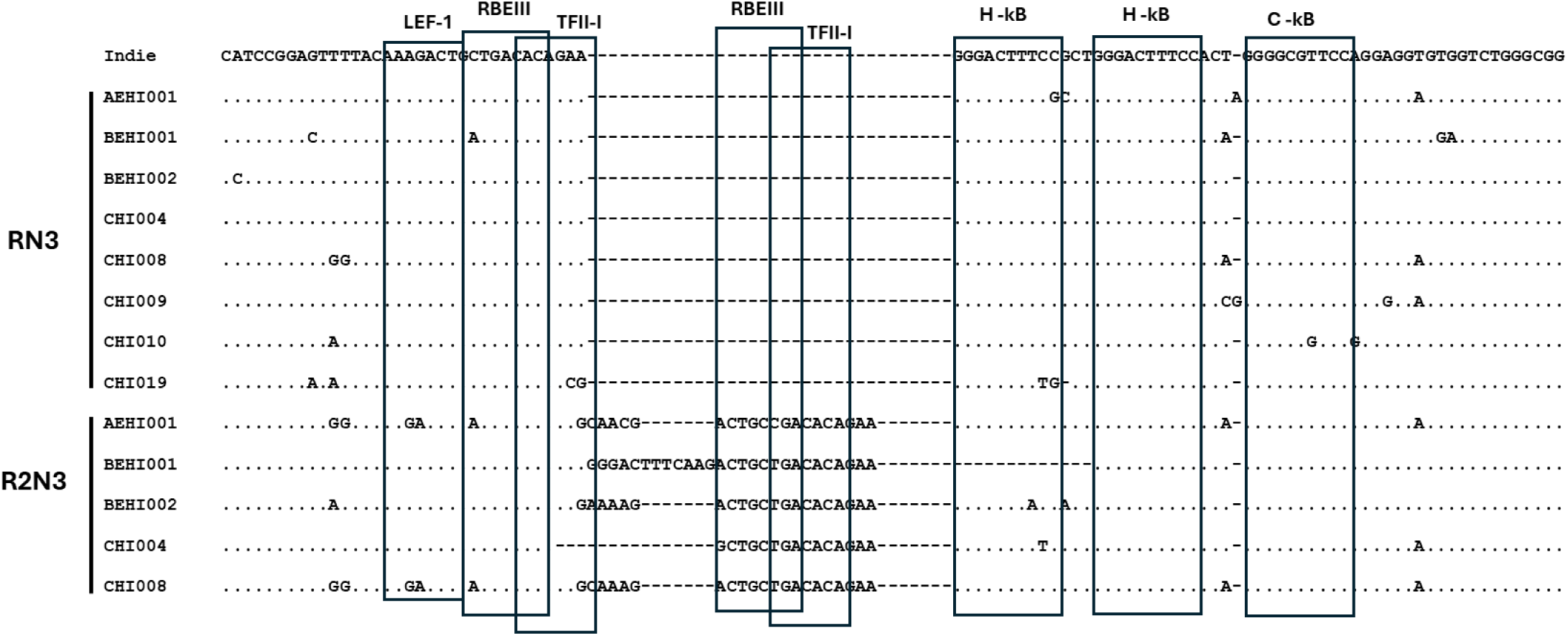
Sequence Alignments of LTR variants from. **Figures 5A** and **6A**. Representative sequences of the major LTR variants observed in the patients from figures – 5A and 6A, aligned with the HIV-1C reference sequence Indie. Dots represent identity and dashes representing deletions. The major transcription factor binding sites are indicated.

**Supplementary Figure 14.**
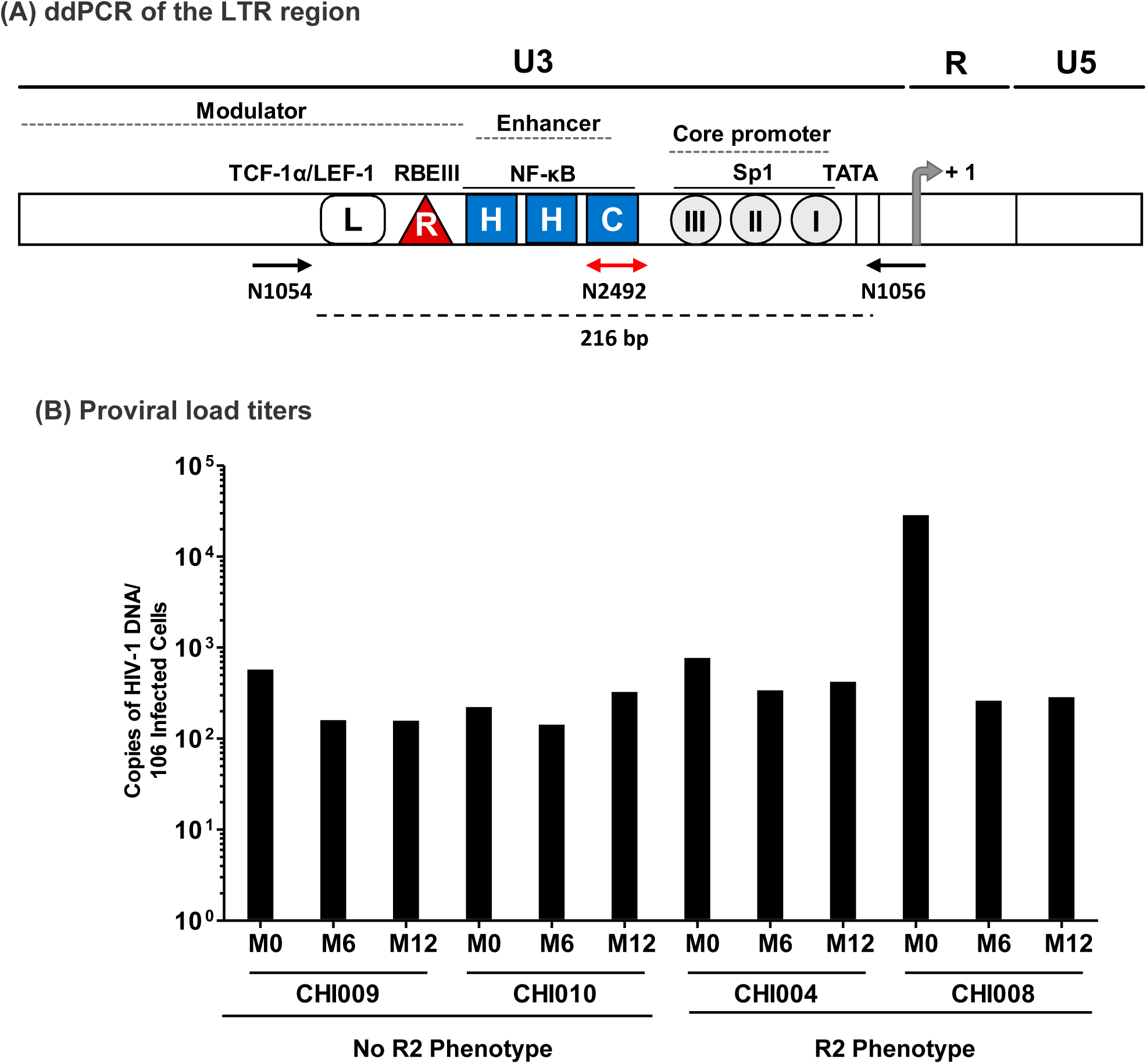
(A) The ddPCR amplification schematic. A 216 bp region of the LTR spanning the regulatory regions of interest is depicted. The primer pair N1054-N1056 (the black arrows) and a TaqMan probe N2492 (the red arrow) are shown. Important TFBS are shown, with the RBEIII and NF-κB motifs, marked as red triangles and blue squares, respectively. (B) Proviral load of four subjects as determined by dd PCR at the base line and two follow-up time points.

**Supplementary Figure 15:**
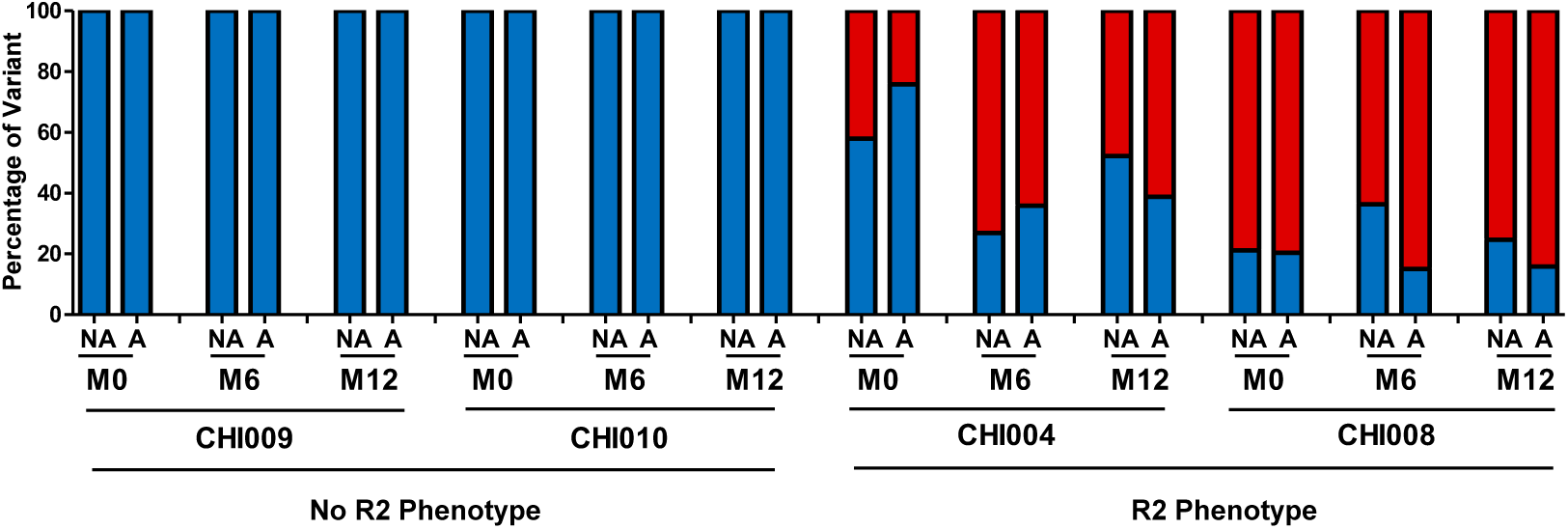
TILDA-seq data from figure 7C, representing absolute proportions of induced variants without normalization with the relative proportions in the viral reservoir.

**Supplementary Figure 16:**
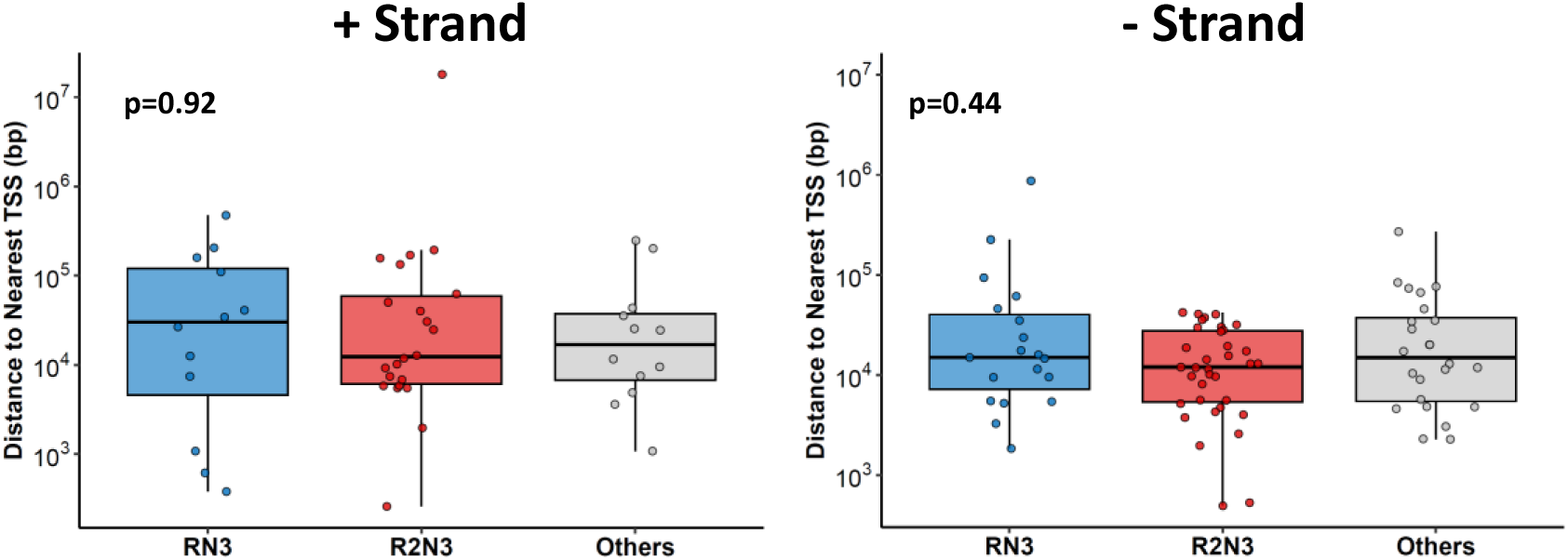
Distance of the integration sites from nearest TSS: The distance to the nearest Transcription Start Site (TSS) from each integration site was analysed using a custom code and classified into two groups based on its presence in the same (+ Strand) or the opposite (-Strand) of the 5’ LTR. The colour code is consistent across figures. The data was compared using Kruskal-Wallis test.

**Supplementary Figure 17:**
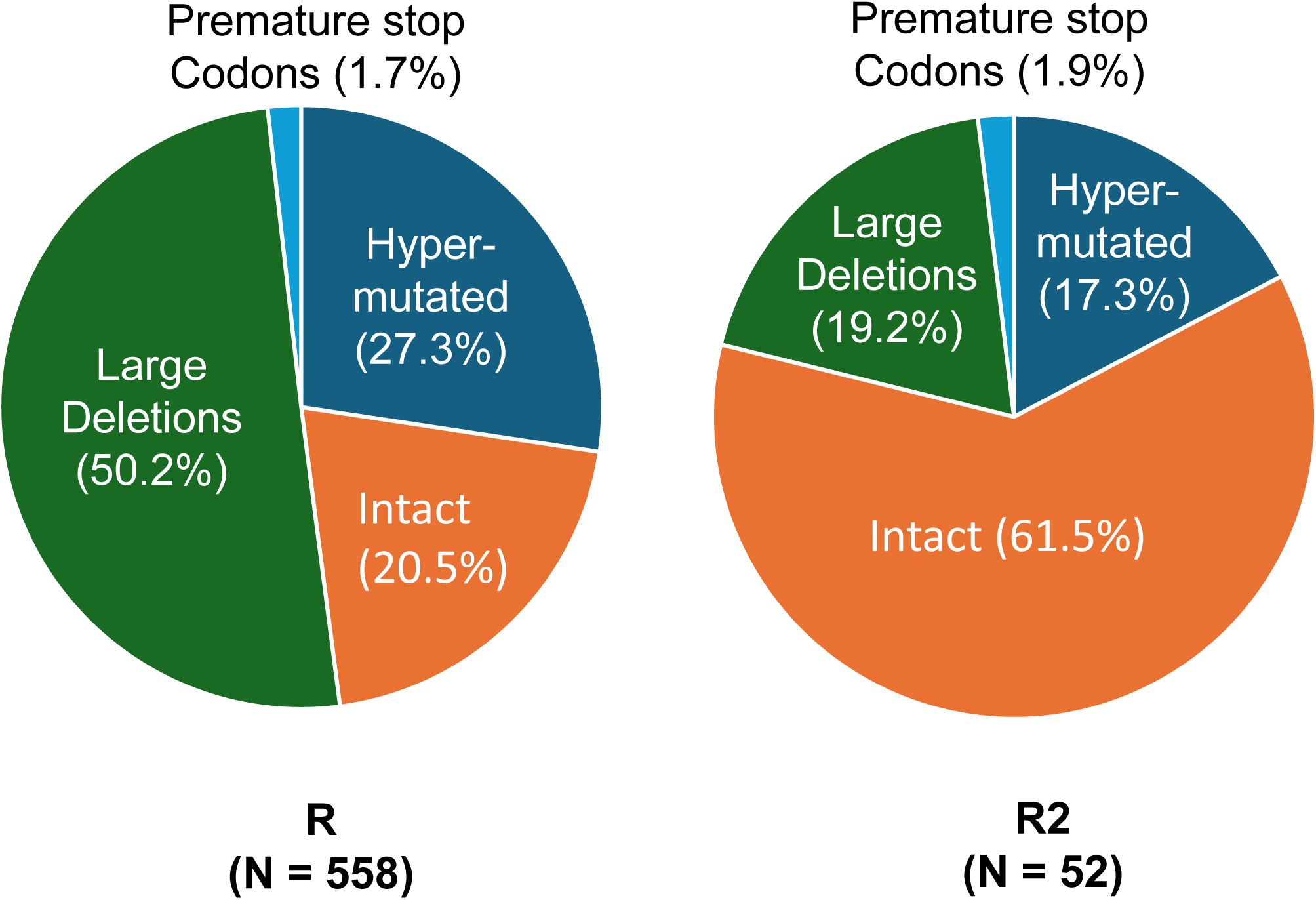
Proviral sequences derived from previous publications that reported several next-generation sequences of single-genome, near-full-length strains (Garcia-Broncano et. al, 2019; Lee et. al, 2019). The sequences were downloaded from GenBank and genotyped into R and R2 viral strains. Genome intactness was analysed using the ProSeq-IT tool (https://psd.cancer.gov/tools/pvs_annot.php) developed by the National Cancer Institute (NIH), and hypermutations were analysed using the Hypermut tool developed by the Los Alamos National Laboratories (https://www.hiv.lanl.gov/content/sequence/HYPERMUT/hypermut.html).

**Supplementary Figure 18:**
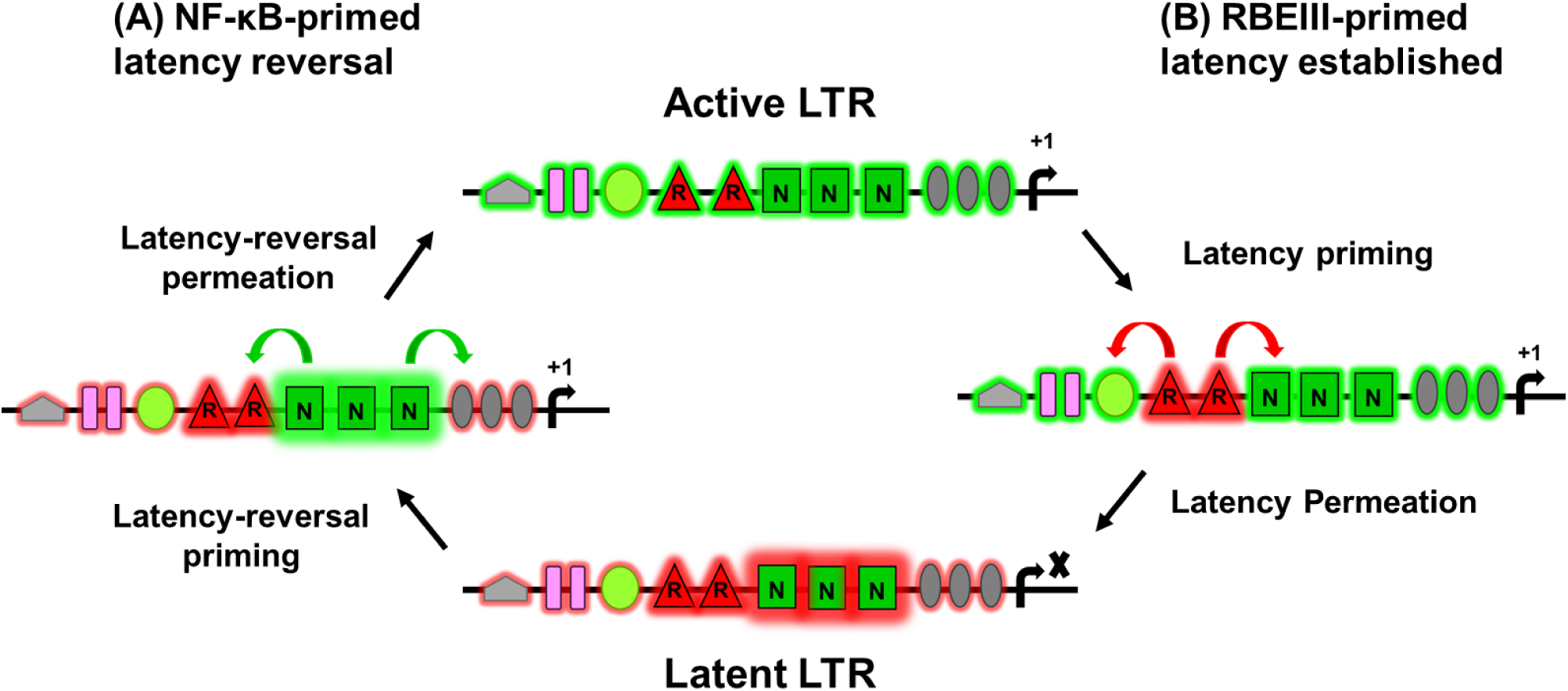
A schematic model for ON/OFF decision-making in HIV-1 gene expression: The interplay between NF-κB and RBEIII motifs governs the transcriptional activity of the HIV-1 LTR, determining whether the promoter remains active or enters latency. Our findings suggest that all TFBS function collectively to interpret environmental cues and drive the LTR towards either an active (ON) or suppressed (OFF) state. The establishment of latency is initiated by the RBEIII motifs and their associated repressive complexes, which gradually extend their influence across other TFBS regions for consorted silencing of transcription. Conversely, the NF-κB motifs play a pivotal role in reactivating latent LTRs by coordinating the activation of other TFBS clusters, resulting in a concerted transcriptional response.

**Supplementary Figure 19:**
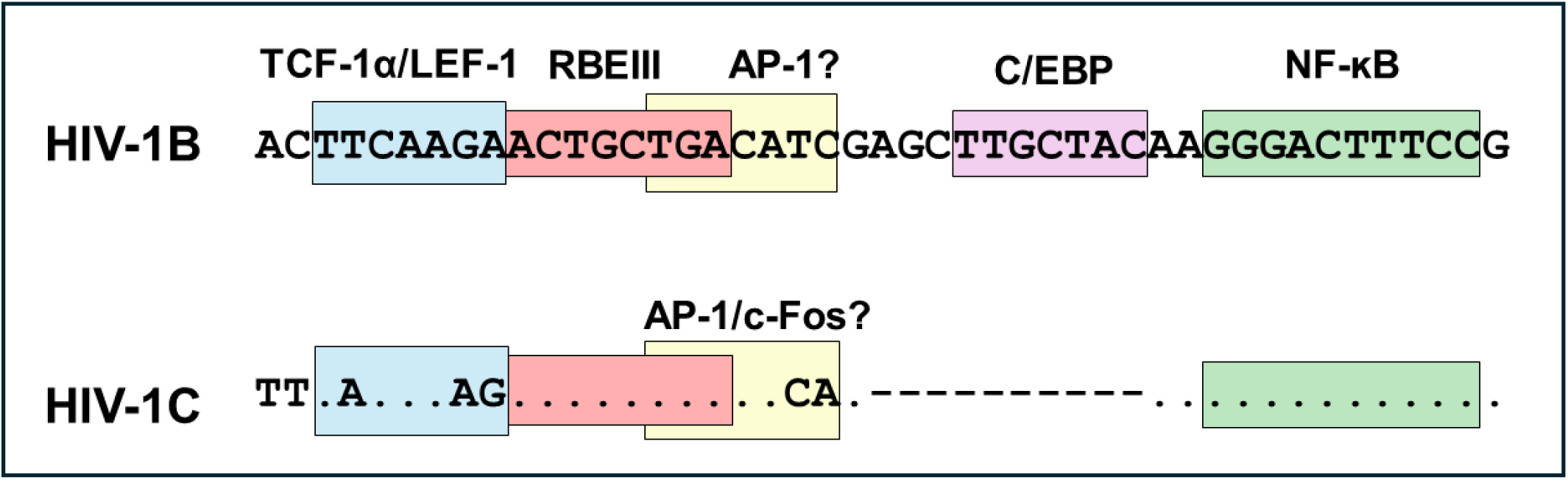
A schematic representation of subtype-specific TFBS variations between HIV-1B and HIV-1C LTR sequences. The comparative sequence analysis uses consensus sequences representative of the respective subtype centred on the RBEIII core motif. The NF-ĸB motif is depicted in green shade, RBEIII in yellow, TCF-1α/ E-1 in pink, AP-1/c-Fos in blue, and TFII-I in grey. Dots and dashes represent sequence homology and deletion, respectively. The coordinates are as per HXb-2. Note that while the RBEIII core motif is conserved between (and among all) the subtypes of HIV-1, the flanking motifs contain subtype-specific variations. ?: TF binding is bioinformatically predicted.

**Supplementary Table 1:** Characteristics of subjects evaluated in Figure 4.

| Subject ID | Age | Sex | PVL (number of RNA copies/ml) | | CD4 (cells/μl) | | |
| --- | --- | --- | --- | --- | --- | --- | --- |
|  |  |  | M0 | M12 | M0 | M6 | M12 |
| 4084 | 38 | F | 6,924 | 5,179 | 509 | 415 | 442 |
| 3767 | 40 | F | 3,742 | - | 566 | 292 | 661 |
| VFSJ15 | 40 | F | 1,914 | 1,710 | 518 | 468 | 367 |
| VFSJ17 | 50 | M | 7,287 | 13,080 | 817 | 954 | 992 |
| VFSJ43 | 36 | F | 39 | 204 | 827 | 827 | 882 |

**Supplementary Table 2:** Percentages of circulating variants in Figure 4A. Asterisk indicates time of ART initiation.

| Sample type | Activation Condition | Month | Replicate 1 |  |  |  |  | Replicate 2 |  |  |  |  |
| --- | --- | --- | --- | --- | --- | --- | --- | --- | --- | --- | --- | --- |
|  |  |  | RN2 | RN3 | R2N2 | R2N3 | R2N4 | RN2 | RN3 | R2N2 | R2N3 | R2N4 |
| Genomic DNA |  | 0 | 0.2 | 99.8 | 0.0 | 0.0 | 0.02 | 0.1 | 99.9 | 0.0 | 0.0 | 0.0 |
|  |  | 6 | 4.0 | 91.8 | 0.0 | 0.1 | 3.96 | 2.8 | 87.9 | 0.0 | 0.3 | 9.1 |
|  |  | 12* | 2.2 | 68.6 | 0.1 | 4.5 | 23.71 | 0.4 | 72.9 | 1.3 | 2.3 | 23.0 |
|  |  | 24 | 0.1 | 51.5 | 0.9 | 12.8 | 34.59 | 0.0 | 56.7 | 0.1 | 8.1 | 35.0 |
|  |  | 36 | 0.0 | 38.2 | 0.0 | 6.1 | 55.49 | 0.2 | 24.7 | 0.1 | 8.9 | 65.9 |
| Cellular RNA | No Activation | 0 | 0.1 | 99.9 | 0.0 | 0.0 | 0.04 | 0.1 | 99.8 | 0.0 | 0.0 | 0.1 |
|  |  | 6 | 1.6 | 97.9 | 0.0 | 0.1 | 0.45 | 0.1 | 99.3 | 0.0 | 0.1 | 0.4 |
|  |  | 12* | 0.1 | 75.7 | 0.0 | 2.0 | 22.09 | 0.1 | 63.3 | 0.0 | 1.3 | 35.1 |
|  |  | 36 | 0.0 | 38.2 | 0.0 | 6.1 | 55.49 | 0.0 | 26.7 | 0.0 | 2.1 | 71.2 |
|  | PMA + Ionomycin | 0 | 0.1 | 99.4 | 0.0 | 0.0 | 0.22 | 0.1 | 99.4 | 0.0 | 0.0 | 0.4 |
|  |  | 6 | 0.4 | 82.0 | 0.0 | 0.7 | 16.81 | 0.5 | 70.0 | 0.0 | 1.0 | 28.2 |
|  |  | 12* | 0.3 | 84.6 | 0.4 | 0.6 | 14.03 | 0.3 | 85.5 | 0.0 | 0.6 | 13.5 |
|  |  | 24 | 0.1 | 79.6 | 0.0 | 0.7 | 19.50 | 0.0 | 0.5 | 0.0 | 2.8 | 96.6 |
|  |  | 36 | 0.0 | 0.3 | 0.0 | 3.6 | 96.00 | 0.9 | 21.5 | 0.0 | 2.4 | 76.0 |
|  | Prostratin | 0 | 0.1 | 99.5 | 0.0 | 0.0 | 0.34 | 0.1 | 99.2 | 0.0 | 0.0 | 0.7 |
|  |  | 6 | 0.3 | 91.9 | 0.0 | 0.4 | 7.30 | 0.2 | 90.2 | 0.0 | 0.3 | 9.3 |
|  |  | 12* | 0.0 | 7.1 | 0.0 | 2.9 | 89.82 | 0.0 | 6.7 | 0.0 | 2.8 | 90.4 |
|  |  | 24 | 0.0 | 0.3 | 0.0 | 3.0 | 96.56 | 0.0 | 0.5 | 0.0 | 2.8 | 96.6 |
|  |  | 36 | 0.0 | 0.2 | 0.0 | 3.0 | 96.70 | 0.0 | 0.2 | 0.0 | 3.0 | 96.8 |

**Supplementary Table 3:** Percentages of circulating variants in Figure 4B.

| Patient | Sample Type | Activation Condition | RN2 | RN3 | R2N2 | R2N3 | R2N4 |
| --- | --- | --- | --- | --- | --- | --- | --- |
| 3767 | gDNA | - | 0.8 | 42.2 | 0.1 | 27.1 | 29.8 |
|  | RNA | No Activation | 0.3 | 12.7 | 2.0 | 84.5 | 0.2 |
|  |  | PMA + Ionomycin | 1.4 | 2.0 | 0.1 | 5.2 | 91.1 |
|  |  | Prostratin | 0.0 | 27.1 | 0.0 | 2.6 | 70.1 |
| 4084 | gDNA | - | 14.5 | 40.7 | 5.7 | 38.7 | 0.1 |
|  | RNA | No Activation | 0.3 | 98.9 | 0.0 | 0.5 | 0.2 |
|  |  | PMA + Ionomycin | 24.5 | 47.9 | 0.6 | 26.1 | 0.1 |
|  |  | Prostratin | 0.0 | 3.3 | 0.1 | 96.5 | 0.1 |
| VFSJ017 | gDNA | - | 0.0 | 30.2 | 0.1 | 69.7 | 0.0 |
|  | RNA | No Activation | 0.0 | 12.0 | 0.1 | 86.3 | 0.4 |
|  |  | PMA + Ionomycin | 0.0 | 2.5 | 0.1 | 97.4 | 0.0 |
|  |  | Prostratin | 0.0 | 7.7 | 0.1 | 86.7 | 1.5 |
| VFSJ043 | gDNA | - | 0.4 | 1.6 | 6.4 | 52.3 | 37.9 |
|  | RNA | No Activation | 0.5 | 2.8 | 10.5 | 84.2 | 1.2 |
|  |  | PMA + Ionomycin | 0.0 | 4.4 | 0.0 | 0.4 | 85.8 |
|  |  | Prostratin | 0.0 | 0.2 | 0.0 | 0.5 | 95.8 |

**Supplementary Table 4:** Characteristics of subjects involved in figure 5.

| Longitudinal Samples (Figure 5A) |  |  |  |  |  |  |  |  |
| --- | --- | --- | --- | --- | --- | --- | --- | --- |
| Subject ID | Age (Y) | Sex | PVL (number of RNA copies/ml) |  |  | CD4 (cells/μl) |  |  |
|  |  |  | M0* | M1 | M3 | M0 | M1 | M3 |
| AEHI001 | 30 | M | 816 | 4040 | <20 | 774 | 886 | 719 |
| AEHI006 | 29 | M | 4030 | 2580 | <20 | 813 | 174 | 921 |
| BEHI001 | 22 | M | 3499 | 4589 | <20 | 308 | 403 | 369 |
| BEHI002 | 20 | M | 251,076 | 795 | <20 | 419 | 672 | 523 |
| Cross-Sectional Samples (Figure 5B) |  |  |  |  |  |  |  |  |
| Sample Number |  |  |  |  |  | 49 |  |  |
| Mean Age (Years ± SD) |  |  |  |  |  | 32.8 ± 8.66 |  |  |
| Female : Male Ratio |  |  |  |  |  | 7:42 |  |  |
| Race |  |  |  |  |  | Asian (n=49) |  |  |
| Median Plasma Viral Load (RNA copies/ml [IQR]) |  |  |  |  |  | 81,048 [165,200] |  |  |
| Median CD4 count (cells/μl [IQR]) |  |  |  |  |  | 372.5 [214.5] |  |  |
| ART status at time of collection |  |  |  |  |  | Naive |  |  |
| IQR - Inter-Quartile Range |  |  |  |  |  |  |  |  |
| SD - Standard Deviation |  |  |  |  |  |  |  |  |

**Supplementary Table 5:** Percentages of circulating variants in Figure 5A. Asterisk indicates the time of ART initiation.

| Activation Condition | Month | AEHI001 |  |  |  |  | AEHI006 |  |  |  |  | BEHI001 |  |  |  |  | BEHI002 |  |  |  |  |
| --- | --- | --- | --- | --- | --- | --- | --- | --- | --- | --- | --- | --- | --- | --- | --- | --- | --- | --- | --- | --- | --- |
|  |  | RN2 | RN3 | R2N2 | R2N3 | R2N4 | RN2 | RN3 | R2N2 | R2N3 | R2N4 | RN2 | RN3 | R2N2 | R2N3 | R2N4 | RN2 | RN3 | R2N2 | R2N3 | R2N4 |
| Genomic DNA | 0* | 2.8 | 92.3 | 0.1 | 2.9 | 0.0 | 0.5 | 98.0 | 0.0 | 0.4 | 0.0 | 3.3 | 94.3 | 0.0 | 1.1 | 0.0 | 0.5 | 94.4 | 0.0 | 4.0 | 0.0 |
|  | 1 | 0.4 | 98.7 | 0.0 | 0.3 | 0.0 | 0.7 | 95.7 | 0.1 | 2.1 | 0.0 | 0.3 | 96.8 | 0.1 | 1.4 | 0.0 | 0.3 | 93.3 | 0.1 | 5.0 | 0.1 |
|  | 3 | 0.3 | 98.2 | 0.0 | 0.9 | 0.0 | 0.1 | 98.1 | 0.0 | 0.5 | 0.0 | 0.3 | 95.7 | 0.0 | 2.8 | 0.1 | 0.3 | 95.0 | 0.0 | 3.6 | 0.0 |
| No Activation | 0* | 2.6 | 95.8 | 0.0 | 0.6 | 0.0 | 0.3 | 98.0 | 0.0 | 0.4 | 0.0 | 2.2 | 92.1 | 0.1 | 4.8 | 0.1 | 1.1 | 20.9 | 0.2 | 77.1 | 0.5 |
|  | 1 | 5.9 | 92.4 | 0.2 | 0.2 | 0.0 | 7.2 | 87.1 | 1.5 | 2.6 | 0.1 | 0.5 | 97.8 | 0.1 | 0.5 | 0.0 | 75.5 | 18.1 | 0.4 | 5.7 | 0.1 |
|  | 3 | 2.4 | 94.4 | 0.0 | 2.2 | 0.0 | 1.9 | 94.3 | 0.0 | 2.4 | 0.0 | 0.3 | 98.5 | 0.0 | 0.4 | 0.0 | 0.2 | 97.5 | 0.0 | 1.3 | 0.0 |
| PMA + Ionomycin | 0* | 2.7 | 91.0 | 0.1 | 5.3 | 0.1 | 1.7 | 93.5 | 0.0 | 3.6 | 0.0 | 0.2 | 98.4 | 0.0 | 0.6 | 0.0 | 0.1 | 2.4 | 0.1 | 96.6 | 0.8 |
|  | 1 | 0.8 | 97.0 | 0.3 | 0.7 | 0.0 | 1.3 | 88.7 | 1.8 | 5.3 | 0.1 | 0.8 | 94.7 | 0.7 | 2.4 | 0.0 | 0.6 | 95.2 | 0.3 | 2.7 | 0.0 |
|  | 3 | 2.8 | 95.2 | 0.0 | 1.1 | 0.0 | 4.1 | 89.9 | 0.1 | 4.6 | 0.0 | 0.4 | 98.1 | 0.0 | 0.6 | 0.0 | 4.5 | 68.4 | 0.3 | 25.8 | 0.2 |
| Prostratin | 0* | 3.9 | 86.5 | 0.3 | 8.3 | 0.1 | 0.5 | 97.6 | 0.0 | 0.9 | 0.0 | 1.0 | 47.0 | 0.1 | 50.9 | 0.5 | 0.9 | 90.9 | 0.2 | 7.1 | 0.1 |
|  | 1 | 4.5 | 93.5 | 0.0 | 1.0 | 0.0 | 0.2 | 98.3 | 0.0 | 0.4 | 0.0 | 0.6 | 95.8 | 0.1 | 2.5 | 0.1 | 68.7 | 22.3 | 0.1 | 8.6 | 0.1 |
|  | 3 | 1.2 | 97.0 | 0.1 | 0.7 | 0.0 | 1.7 | 96.3 | 0.1 | 0.8 | 0.0 | 3.9 | 81.6 | 0.6 | 7.3 | 0.1 | 3.0 | 77.2 | 0.5 | 18.4 | 0.2 |
| Plasma | 0* | 0.4 | 97.8 | 0.0 | 0.3 | 0.0 | 0.3 | 96.8 | 0.0 | 0.3 | 0.0 | 1.2 | 91.0 | 0.1 | 5.7 | 0.1 | 0.5 | 15.0 | 0.5 | 83.1 | 0.7 |
|  | 1 | 1.6 | 96.1 | 0.0 | 0.7 | 0.0 | 3.5 | 93.5 | 0.0 | 0.0 | 0.0 | 0.1 | 98.4 | 0.0 | 0.2 | 0.0 | 0.5 | 94.3 | 0.0 | 3.6 | 0.0 |
|  | 3 | 4.0 | 93.9 | 0.4 | 0.7 | 0.0 | 3.4 | 94.1 | 0.4 | 0.7 | 0.0 | 3.0 | 92.9 | 0.3 | 2.5 | 0.1 | 3.6 | 73.7 | 0.6 | 20.9 | 0.3 |

**Supplementary Table 6:** Percentages of circulating variants in Figure 5B.

| Patient ID | Variant |  |  |  |  |  |
| --- | --- | --- | --- | --- | --- | --- |
|  | RN3 | RN2 | RN4 | R2N2 | R2N3 | R2N4 |
| **RN3 (Can.) - 34/49 (69.4%)** |  |  |  |  |  |  |
| 5 | 99.7 | 0.2 | 0.2 | 0.0 | 0.0 | 0.0 |
| 8 | 99.8 | 0.1 | 0.1 | 0.0 | 0.0 | 0.0 |
| 11 | 99.6 | 0.2 | 0.2 | 0.0 | 0.0 | 0.0 |
| 12 | 99.6 | 0.1 | 0.3 | 0.0 | 0.0 | 0.0 |
| 13 | 99.6 | 0.3 | 0.1 | 0.0 | 0.0 | 0.0 |
| 17 | 99.6 | 0.2 | 0.2 | 0.0 | 0.0 | 0.0 |
| 18 | 99.7 | 0.1 | 0.2 | 0.0 | 0.0 | 0.0 |
| 19 | 99.5 | 0.2 | 0.2 | 0.0 | 0.2 | 0.0 |
| 20 | 99.5 | 0.2 | 0.2 | 0.0 | 0.0 | 0.0 |
| 21 | 99.6 | 0.2 | 0.2 | 0.0 | 0.0 | 0.0 |
| 22 | 99.7 | 0.2 | 0.1 | 0.0 | 0.1 | 0.0 |
| 25 | 99.6 | 0.3 | 0.2 | 0.0 | 0.0 | 0.0 |
| 27 | 99.9 | 0.1 | 0.1 | 0.0 | 0.0 | 0.0 |
| 31 | 99.8 | 0.1 | 0.1 | 0.0 | 0.0 | 0.0 |
| 33 | 99.6 | 0.2 | 0.2 | 0.0 | 0.0 | 0.0 |
| 34 | 99.7 | 0.2 | 0.1 | 0.0 | 0.0 | 0.0 |
| 36 | 99.7 | 0.1 | 0.2 | 0.0 | 0.0 | 0.0 |
| 37 | 99.6 | 0.1 | 0.2 | 0.0 | 0.0 | 0.0 |
| 38 | 99.6 | 0.2 | 0.2 | 0.0 | 0.0 | 0.0 |
| 41 | 99.3 | 0.5 | 0.2 | 0.0 | 0.0 | 0.0 |
| 42 | 99.6 | 0.3 | 0.2 | 0.0 | 0.0 | 0.0 |
| 44 | 99.7 | 0.2 | 0.1 | 0.0 | 0.0 | 0.0 |
| 45 | 99.6 | 0.2 | 0.1 | 0.0 | 0.0 | 0.0 |
| 47 | 99.6 | 0.2 | 0.2 | 0.0 | 0.0 | 0.0 |
| 48 | 99.5 | 0.2 | 0.3 | 0.0 | 0.0 | 0.0 |
| 49 | 99.7 | 0.2 | 0.1 | 0.0 | 0.0 | 0.0 |
| 50 | 99.7 | 0.2 | 0.1 | 0.0 | 0.0 | 0.0 |
| 58 | 99.7 | 0.2 | 0.2 | 0.0 | 0.0 | 0.0 |
| 59 | 99.6 | 0.1 | 0.3 | 0.0 | 0.0 | 0.0 |
| 60 | 99.6 | 0.3 | 0.2 | 0.0 | 0.0 | 0.0 |
| 61 | 99.7 | 0.2 | 0.2 | 0.0 | 0.0 | 0.0 |
| 62 | 99.8 | 0.1 | 0.1 | 0.0 | 0.0 | 0.0 |
| 70 | 99.5 | 0.1 | 0.3 | 0.0 | 0.1 | 0.0 |
| 71 | 97.9 | 0.4 | 1.6 | 0.0 | 0.0 | 0.0 |
| **R2 Phenotype - 9/49 (18.4%)** |  |  |  |  |  |  |
| 14 | 96.7 | 3.2 | 0.1 | 0.1 | 0.0 | 0.0 |
| 23 | 15.0 | 0.0 | 0.0 | 0.2 | 84.7 | 0.2 |
| 28 | 45.0 | 0.0 | 0.1 | 0.1 | 54.6 | 0.2 |
| 32 | 95.2 | 4.5 | 0.2 | 0.1 | 0.0 | 0.0 |
| 39 | 94.1 | 5.7 | 0.2 | 0.0 | 0.0 | 0.0 |
| 40 | 88.6 | 0.0 | 0.4 | 0.0 | 11.0 | 0.0 |
| 63 | 81.2 | 4.2 | 0.1 | 0.0 | 14.5 | 0.0 |
| 72 | 83.7 | 16.0 | 0.2 | 0.1 | 0.0 | 0.0 |
| 73 | 14.3 | 0.0 | 0.0 | 0.2 | 85.2 | 0.3 |
| **Others - 6/49 (12.2%)** |  |  |  |  |  |  |
| 29 | 53.7 | 0.1 | 46.3 | 0.0 | 0.0 | 0.0 |
| 30 | 0.7 | 0.0 | 99.3 | 0.0 | 0.0 | 0.0 |
| 51 | 81.3 | 2.0 | 16.7 | 0.0 | 0.0 | 0.0 |
| 53 | 10.3 | 0.0 | 89.2 | 0.0 | 0.4 | 0.0 |
| 55 | 4.3 | 0.0 | 95.6 | 0.0 | 0.0 | 0.0 |
| 57 | 36.5 | 0.1 | 63.3 | 0.0 | 0.0 | 0.0 |

**Supplementary Table 7:** Characteristics of subjects involved in figure 6.

| Longitudinal Samples (Figure 6A) |  |  |  |  |  |  |  |  |
| --- | --- | --- | --- | --- | --- | --- | --- | --- |
| Subject ID | Age (Y) | Sex | PVL<br>(number of RNA copies/ml) |  |  | CD4 (cells/μl) |  |  |
|  |  |  | M0* | M6 | M12 | M0 | M1 | M3 |
| CHI009 | 23 | M | 4,980 | <20 | <20 | 541 | 634 | 659 |
| CHI010 | 26 | M | 121,000 | <20 | <20 | 250 | 425 | 576 |
| CHI004 | 38 | M | 42,543 | <20 | <20 | 257 | 387 | 511 |
| CHI008 | 34 | M | 796,433 | <20 | <20 | 247 | 506 | 537 |
| Cross-Sectional Samples (Figure 6B) |  |  |  |  |  |  |  |  |
| Sample Number |  |  |  |  |  | 40 |  |  |
| Mean Age (Years ± SD) |  |  |  |  |  | 33.02 ± 10.55 |  |  |
| Female : Male Ratio |  |  |  |  |  | 12:28 |  |  |
| Race |  |  |  |  |  | Asian (n=40) |  |  |
| Median Plasma Viral Load (RNA copies/ml [IQR]) |  |  |  |  |  | 98,905 [598,019] |  |  |
| Median CD4 count (cells/μl [IQR]) |  |  |  |  |  | 337 [485] |  |  |
| ART status at time of collection |  |  |  |  |  | Naive |  |  |
| IQR - Inter-Quartile Range |  |  |  |  |  |  |  |  |
| SD - Standard Deviation |  |  |  |  |  |  |  |  |

**Supplementary Table 8:** Percentages of circulating variants in Figure 6A. Asterisk indicates time of ART initiation.

| Activation Condition | Month | CHI009 |  |  |  |  | CHI010 |  |  |  |  | CHI004 |  |  |  |  | CHI008 |  |  |  |  |
| --- | --- | --- | --- | --- | --- | --- | --- | --- | --- | --- | --- | --- | --- | --- | --- | --- | --- | --- | --- | --- | --- |
|  |  | RN2 | RN3 | R2N2 | R2N3 | R2N4 | RN2 | RN3 | R2N2 | R2N3 | R2N4 | RN2 | RN3 | R2N2 | R2N3 | R2N4 | RN2 | RN3 | R2N2 | R2N3 | R2N4 |
| Genomic DNA | 0* | 0.6 | 97.8 | 0.0 | 0.5 | 0.0 | 1.8 | 96.6 | 0.1 | 0.7 | 0.0 | 13.0 | 42.8 | 0.2 | 42.1 | 0.6 | 0.4 | 26.0 | 0.5 | 71.6 | 1.0 |
|  | 6 | 0.3 | 97.5 | 0.0 | 0.8 | 0.0 | 0.3 | 97.7 | 0.1 | 0.7 | 0.0 | 6.8 | 54.2 | 0.2 | 37.6 | 0.4 | 0.2 | 2.4 | 0.7 | 96.6 | 0.1 |
|  | 12 | 0.4 | 97.1 | 0.0 | 1.4 | 0.0 | 0.4 | 97.1 | 0.1 | 1.4 | 0.0 | 1.7 | 10.3 | 0.0 | 87.8 | 0.1 | 1.0 | 6.4 | 0.5 | 90.8 | 1.2 |
| No Activation | 0* | 0.8 | 97.2 | 0.0 | 0.9 | 0.0 | 1.9 | 97.0 | 0.0 | 0.3 | 0.0 | 13.0 | 52.9 | 0.3 | 32.7 | 0.5 | 0.2 | 15.8 | 1.0 | 82.2 | 0.7 |
|  | 6 | 4.7 | 93.7 | 0.2 | 0.5 | 0.0 | 4.8 | 85.6 | 4.6 | 0.4 | 1.6 | 9.3 | 52.3 | 0.2 | 37.5 | 0.3 | 0.8 | 29.9 | 0.6 | 68.0 | 0.6 |
|  | 12 | 0.1 | 98.7 | 0.0 | 0.1 | 0.0 | 0.9 | 97.2 | 0.1 | 0.9 | 0.0 | 8.8 | 48.5 | 0.8 | 41.2 | 0.3 | 0.2 | 26.5 | 0.2 | 72.4 | 0.5 |
| PMA + Ionomycin | 0* | 0.3 | 98.2 | 0.0 | 0.7 | 0.0 | 1.7 | 97.6 | 0.0 | 0.6 | 0.0 | 3.1 | 55.5 | 0.4 | 40.1 | 0.4 | 0.5 | 19.8 | 0.6 | 78.2 | 0.7 |
|  | 6 | 41.9 | 61.5 | 0.5 | 1.5 | 0.0 | 1.1 | 92.3 | 1.5 | 3.2 | 0.1 | 1.0 | 95.5 | 0.2 | 2.2 | 0.0 | 0.1 | 13.1 | 0.4 | 85.5 | 0.8 |
|  | 12 | 45.4 | 51.3 | 0.1 | 2.5 | 0.0 | 0.5 | 98.2 | 0.0 | 0.4 | 0.0 | 29.9 | 40.9 | 0.1 | 28.3 | 0.4 | 0.4 | 15.7 | 0.3 | 82.6 | 0.7 |
| Prostratin | 0* | 0.2 | 98.5 | 0.0 | 0.5 | 0.0 | 1.7 | 96.8 | 0.0 | 0.6 | 0.0 | 3.6 | 87.9 | 0.1 | 7.4 | 0.1 | 0.2 | 30.1 | 0.2 | 67.9 | 1.3 |
|  | 6 | 0.8 | 97.6 | 0.0 | 0.6 | 0.0 | 1.3 | 94.7 | 0.1 | 2.9 | 0.1 | 21.3 | 34.8 | 0.6 | 42.7 | 0.2 | 0.1 | 9.3 | 0.2 | 89.5 | 0.8 |
|  | 12 | 1.5 | 95.9 | 0.1 | 1.3 | 0.0 | 0.6 | 97.9 | 0.1 | 0.6 | 0.0 | 44.3 | 27.2 | 0.3 | 27.8 | 0.1 | 2.8 | 10.4 | 11.9 | 74.1 | 0.8 |
| Plasma | 0* | 0.3 | 98.1 | 0.0 | 0.4 | 0.0 | 2.8 | 94.4 | 0.1 | 1.7 | 0.0 | 14.7 | 67.5 | 0.2 | 16.5 | 0.5 | 0.2 | 31.9 | 1.0 | 65.6 | 0.8 |
|  | 6 | 1.1 | 96.9 | 0.0 | 0.4 | 0.0 | 0.9 | 97.3 | 0.1 | 0.4 | 0.0 | 10.2 | 65.7 | 0.2 | 22.6 | 0.2 | 0.4 | 57.8 | 1.2 | 39.6 | 0.5 |
|  | 12 | 0.5 | 97.7 | 0.0 | 0.5 | 0.0 | 3.1 | 95.0 | 0.3 | 0.4 | 0.0 | 1.7 | 94.4 | 0.1 | 2.5 | 0.0 | 0.7 | 92.5 | 0.2 | 5.4 | 0.1 |

**Supplementary Table 9:** Percentages of circulating variants in Figure 6B.

| Patient ID | Variant |  |  |  |  |  |
| --- | --- | --- | --- | --- | --- | --- |
|  | RN2 | RN3 | RN4 | R2N2 | R2N3 | R2N4 |
| **RN3 (Can.) - 12/40 (30%)** |  |  |  |  |  |  |
| 7 | 0.0 | 98.6 | 0.5 | 0.2 | 0.0 | 0.0 |
| 9 | 0.0 | 98.9 | 0.6 | 0.0 | 0.0 | 0.3 |
| 10 | 0.1 | 98.7 | 0.9 | 0.0 | 0.2 | 0.0 |
| 23 | 0.4 | 98.5 | 0.9 | 0.0 | 0.3 | 0.0 |
| 37 | 2.0 | 96.3 | 1.3 | 0.0 | 0.3 | 0.1 |
| 55 | 1.2 | 96.9 | 0.7 | 0.7 | 0.5 | 0.1 |
| 65 | 1.6 | 96.4 | 1.8 | 0.0 | 0.1 | 0.1 |
| 72 | 0.1 | 98.6 | 1.1 | 0.0 | 0.2 | 0.0 |
| 73 | 0.4 | 98.3 | 1.3 | 0.0 | 0.0 | 0.0 |
| 84 | 0.9 | 98.1 | 0.6 | 0.0 | 0.3 | 0.1 |
| 85 | 0.2 | 99.0 | 0.6 | 0.0 | 0.3 | 0.0 |
| 86 | 0.1 | 98.6 | 1.1 | 0.0 | 0.2 | 0.0 |
| **R2 Phenotype - 19/40 (47.5%)** |  |  |  |  |  |  |
| 1 | 1.8 | 24.0 | 0.2 | 0.4 | 72.9 | 0.7 |
| 4 | 17.4 | 37.2 | 0.0 | 0.1 | 45.1 | 0.0 |
| 5 | 0.2 | 3.1 | 0.1 | 0.9 | 95.3 | 0.4 |
| 6 | 0.4 | 96.0 | 0.3 | 0.0 | 3.3 | 0.0 |
| 8 | 0.2 | 23.1 | 0.9 | 0.1 | 75.2 | 0.2 |
| 11 | 0.5 | 39.5 | 0.6 | 0.1 | 58.1 | 1.2 |
| 15 | 0.8 | 8.3 | 1.8 | 1.5 | 1.7 | 86.0 |
| 21 | 5.3 | 87.7 | 5.1 | 0.1 | 1.8 | 0.1 |
| 22 | 4.2 | 40.2 | 0.6 | 1.6 | 49.2 | 4.2 |
| 25 | 7.2 | 6.8 | 0.1 | 5.1 | 80.5 | 0.4 |
| 32 | 10.7 | 8.4 | 0.1 | 1.0 | 79.6 | 0.2 |
| 35 | 0.5 | 12.7 | 0.3 | 1.5 | 84.3 | 0.8 |
| 63 | 0.7 | 16.6 | 0.3 | 11.6 | 70.4 | 0.4 |
| 67 | 4.8 | 87.9 | 1.8 | 0.7 | 4.8 | 0.1 |
| 70 | 0.7 | 94.9 | 1.1 | 0.0 | 3.2 | 0.0 |
| 71 | 13.0 | 61.7 | 0.6 | 0.2 | 22.3 | 2.2 |
| 81 | 6.8 | 66.2 | 0.6 | 0.0 | 0.8 | 25.5 |
| 87 | 3.9 | 28.5 | 62.6 | 0.1 | 2.9 | 2.0 |
| 99 | 0.6 | 63.6 | 0.7 | 0.1 | 33.9 | 1.1 |
| **Others - 9/40 (22.5%)** |  |  |  |  |  |  |
| 41 | 5.9 | 92.9 | 0.9 | 0.0 | 0.3 | 0.0 |
| 54 | 0.7 | 95.2 | 3.9 | 0.0 | 0.2 | 0.0 |
| 58 | 0.6 | 96.2 | 2.9 | 0.0 | 0.3 | 0.1 |
| 74 | 9.9 | 83.6 | 6.4 | 0.0 | 0.0 | 0.0 |
| 75 | 17.1 | 80.9 | 1.8 | 0.1 | 0.2 | 0.0 |
| 79 | 2.2 | 95.4 | 1.0 | 0.0 | 1.3 | 0.2 |
| 88 | 3.1 | 95.2 | 1.2 | 0.0 | 0.4 | 0.2 |
| 89 | 2.4 | 94.7 | 2.3 | 0.0 | 0.5 | 0.1 |
| 90 | 3.8 | 78.8 | 17.1 | 0.0 | 0.2 | 0.1 |

## Notes

### Competing Interest Statement

The authors have declared no competing interest.

### Summary of Updates

This version of the manuscript has been revised to improve the figure quality for several figures.

